# Fine-tuning an intracellular allosteric pocket programs GPCR signaling bias

**DOI:** 10.64898/2026.10.22.760766

**Authors:** Xinchao Shi, Ruidi Zhu, Kewei Chang, Peng Yu, Qingning Yuan, Yunyun Zhang, Xun Lu, Ziwei Zhou, Chunhao Zhu, Xiaobing Lan, Mingyu Li, Rui Li, Xin Qiao, Weidi Xie, Youwen Zhuang, H Eric Xu, Wanchao Yin, Mingzhu Zhao, Jian Zhang

## Abstract

Biased signaling of G protein-coupled receptors (GPCRs) offers a route for precise therapeutics with fewer adverse effects^1^. Intracellular allosteric ligands can impose such bias, yet how subtle chemical changes within these pockets reshape receptor-transducer coupling and downstream signaling remains unclear. Here, we systematically fine-tuned the intracellular allosteric pocket of neurotensin receptor 1 (NTSR1) through chemical modification of the β-arrestin-biased ligand SBI-553^2, 3^. Minor changes to the ligand scaffold altered transducer-coupling preferences, generating distinct profiles of G-protein and β-arrestin signaling. Cryo-electron microscopy structures of NTSR1–β-arrestin2 complexes bound to designed allosteric probes revealed ligand-dependent remodeling of the intracellular cavity that reshapes the receptor-effector interface. Alanine scanning identified transducer-sensitive residues within and beyond the immediate ligand-contact region. In mice, SBI-553 and N065 produced distinct physiological profiles, with N065 improving survival in a lethal endotoxemia model. Finally, phosphoproteomic profiling revealed partially overlapping but distinct downstream signaling profiles among the three modulators. Together, these findings establish the intracellular allosteric pocket as a tunable determinant of transducer coupling and show that closely related modulators can generate distinct signaling and physiological profiles.

## Introduction

G protein-coupled receptors (GPCRs) constitute the largest family of membrane signaling proteins and represent targets for roughly one-third of all approved drugs^4^. A central concept in GPCR pharmacology is biased signaling, in which different ligands stabilize distinct receptor conformations that preferentially engage specific downstream transducers such as G proteins or β-arrestins^5, 6^. By selectively activating beneficial signaling pathways while avoiding those associated with adverse effects, biased ligands hold considerable promise for improving the therapeutic index of GPCR-targeting drugs^7, 8^.

Most efforts to achieve signaling bias have focused on orthosteric ligands that bind within the endogenous agonist pocket^9–12^. Allosteric modulators, which bind to spatially distinct receptor sites, provide an alternative strategy to regulate receptor signaling^13–15^. Among these, intracellular allosteric pockets located at the interface between GPCRs and their downstream effectors have recently emerged as particularly effective regulatory nodes. Ligands occupying these cavities can directly reshape the interaction surface for downstream transducers, either stabilizing effector engagement through additional contacts or sterically hindering coupling to specific signaling partners^16^. Recent structural and pharmacological studies have demonstrated that such intracellular biased allosteric modulators (BAMs) can selectively biasing GPCR coupling toward specific G-protein subtypes or arrestins, suggesting the potential of these sites for precise control of receptor signaling^3, 17–24^. In particular, the intracellular neurotensin receptor 1 (NTSR1) modulator SBI-553 and related analogues were shown to tune G-protein coupling by acting as molecular glues or molecular bumpers, establishing the intracellular allosteric pocket it occupies as a tunable node for transducer selectivity^16^. Whether signaling bias can also be tuned by systematically perturbing distinct regions of the intracellular pocket, and whether such local chemical perturbations influence receptor conformational microstates beyond the immediate transducer interface, remain less well understood.

In this study, we refined the chemical scaffold of SBI-553 to probe how subtle modifications within the intracellular cavity reshape NTSR1 signaling. By combining ligand design, cryo-electron microscopy (cryo-EM) structural analysis, biochemical assays and *in vivo* physiology, we show that minor chemical perturbations in the intracellular allosteric pocket can redistribute G-protein and β-arrestin coupling to generate distinct signaling signatures. Alanine scanning of the allosteric pocket together with adjacent conserved motifs further revealed potential transducer-sensitive hotspots that shapes transducer signaling output of NTSR1. These residues were distributed both within the pocket and in conserved motifs outside the pocket, suggesting that intracellular allosteric ligands could reshape NTSR1 signaling not only by directly modulating the receptor–transducer interface, but also by shifting the conformational ensemble associated with receptor activation. Such signaling differences may translate *in vivo* into divergent NTSR1-dependent phenotypes^2,25–27^, including altered thermoregulation, blood pressure and metabolic responses. Allosteric regulation mediated by the modified molecule N065 that partially attenuates G_q_-dependent thermogenic signaling stabilizes an adaptive hypothermic state and improves survival in a murine endotoxemia model. Together, our findings reveal how fine-tuning of intracellular allostery programs GPCR signaling in physiological and pathological contexts.

## Results

### Allosteric molecules program signaling bias in the intracellular pocket

NTSR1 is a class A GPCR that engages multiple transducers, including G_i/o_, G_q/11_, G_12/13_, and β-arrestins, upon binding its orthosteric peptide ligand neurotensin (NTS) (Fig. 1A)^28^. Prior structural studies have identified a distinct intracellular allosteric site on the cytoplasmic face of NTSR1, where the BAM SBI-553 acts as a negative allosteric modulator (NAM) for G protein signaling and an agonist positive allosteric modulator (ago-PAM) for β-arrestin signaling^3, 23^. SBI-553 was suggested to occupy an intracellular cavity within the hydrophobic core of NTSR1, contacting key microswitches of the conserved class A GPCR activation machinery^3, 23^. It is positioned next to the TM3 ERY (DRY) motif and extends toward the TM7 NPxxY motif that undergo coordinated rearrangements during receptor activation. Building on this structural framework (Extended Data Fig. 1A), we hypothesized that precise chemical modification on SBI-553 within this pocket could remodel the effector interface or stabilize receptor conformations favoring specific transducers (Fig. 1A). We therefore designed four SBI-553 analogs to introduce localized modifications across distinct regions of the intracellular pocket to probe their contributions to signaling bias. These four analogs, N065, N253, N191, and N209, serve as chemical probes for the entrance-facing head region, the mid-pocket core, the cytoplasmic-facing side chain, and the deep anchoring region, respectively (Fig. 1B).

**Fig. 1.**
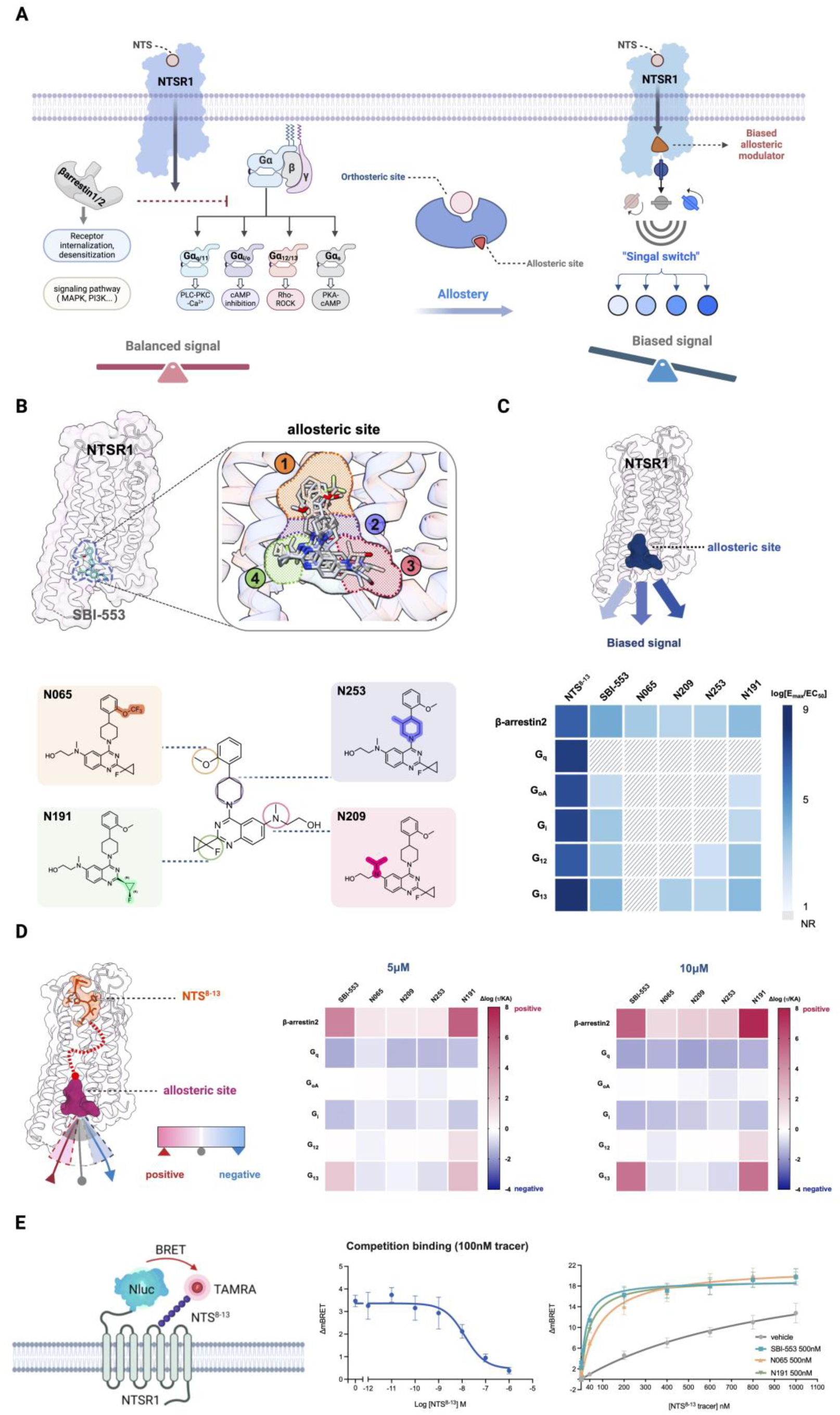
| Intracellular allostery at NTSR1. **(A)** Conceptual schematic of NTSR1 signaling. Left, orthosteric NTS binding supports balanced coupling to β-arrestins and multiple G-protein families. Right, occupancy of an intracellular allosteric site by a BAM acts as a “signal switch,” redistributing signaling output toward a biased profile. **(B)** Localization of SBI-553 at the intracellular allosteric site of NTSR1 and structural decomposition of the pocket into four subregions (labeled 1–4) that accommodate distinct substituent vectors. Chemical structures of SBI-553 and four representative analogs (N065, N209, N253, N191) are shown with highlighted substituent variations. **(C)** Structural depiction of the intracellular allosteric pocket mapped onto NTSR1 (dark blue) and pathway efficiency heat map for NTS^8–13^, SBI-553 and representative analogs across β-arrestin2 and G-protein pathways. Signaling efficiency is displayed as log (*E_max_/EC_50_*); NR, no response. **(D)** Pathway-selective allosteric modulation of NTS^8–13^ signaling by intracellular ligands at fixed concentrations (5 μM and 10 μM). Heat maps report Δlog (*τ/K_A_*) values derived from operational model analysis; positive values indicate potentiation and negative values indicate inhibition of the orthosteric response. **(E)** NanoBRET binding assay for orthosteric ligand engagement. Left, schematic of Nluc-NTSR1 and TAMRA-NTS^8–13^ tracer BRET configuration. BRET signals are expressed as milliBRET units (mBU; BRET ratio × 1,000). Middle, competition binding of unlabeled NTS^8–13^ against 100 nM tracer. Right, tracer binding in the presence of vehicle or 500 nM intracellular ligands (SBI-553, N065, N191). Data are presented as mean ± s.e.m. from three independent experiments.

To profile their signaling properties, we assessed intrinsic agonism of SBI-553 and all four analogs across β-arrestin2 (βarr2) and multiple G protein pathways (G_q_, G_i_, G_oA_, G_12_ and G_13_) using BRET-based assays (Extended Data Fig. 1C, Fig. 1C). We found that none of the compounds activated G_q_. SBI-553 and N191 exhibited agonist activity toward G_i_ and G_oA_, with more pronounced signaling output on G_12/13_. Additionally, N191 demonstrated greater βarr2 recruitment efficacy than SBI-553. The remaining analogs showed more restricted G protein subtype engagement. Among them, N253 displayed weak activation on G_12/13_, N209 exhibited weak agonism selectively at G_13_, and N065 produced no detectable G protein activity, indicating a highly convergent signaling profile. Across all five compounds, the hierarchy of G protein preference follows the trend G_12/13_ > G_i/o_ > G_q_ (Fig. 1C, Extended Data Figs. 1C–D). Despite divergent G protein activity, all intracellular ligands enhanced β-arrestin2 signaling, consistent with their ago-PAM activity at NTSR1 (Extended Data Fig. 2G).

In the presence of NTS, all five compounds reduced G_q_ potency and efficacy, except N065, which selectively decreased potency with minimal effect on efficacy (Extended Data Fig. 2B). Across G_i/o_ and G_12/13_ pathways, N065, N209, and N253 primarily acted as NAMs, although N253 exhibited weak PAM activity at G_12_ signaling of NTS. In contrast, SBI-553 and N191 exhibited bidirectional allosteric effects consistent with their intrinsic agonism in these pathways (Extended Data Figs. 2C–F). In G_12/13_, both compounds displayed ago-PAM behavior at higher concentrations but reduced NTS potency at sub-saturating levels. A similar pattern was observed in G_i/o_ pathways, where intrinsic agonism coexisted with reductions in NTS potency or efficacy (Extended Data Figs. 2C–F). These complex effects likely arise from combined classical receptor allostery and intracellular ligand-mediated effects, including molecular glue-like cooperativity and steric hindrance of transducer engagement. Accordingly, we selected N065, which showed selective agonist activity toward βarr2 with no detectable G-protein activity, and N191, a higher-efficacy βarr2–biased agonist relative to SBI-553, as representative probes to dissect transducer-selective signaling relative to the parental scaffold. Despite distinct signaling outputs, all the three intracellular ligands similarly enhanced binding of the fluorescent orthosteric ligand (Fig. 1E, Fig S1), supporting positive cooperativity between intracellular pocket occupancy and orthosteric ligand binding^2^.

### Structures of NTS–NTSR1–βarr2 complexes with intracellular modulators

Building upon our functional observations that SBI-553 and its structural analogues exert profound, differentiated modulation of NTSR1 signaling outputs, we sought to unveil the structural underpinnings by which chemically distinct intracellular BAMs engage NTSR1 when coupled to βarr2. To this end, we determined cryo-EM structures of NTSR1–βarr2 complexes simultaneously bound to NTS^8–13^ and one of the three modulators (SBI-553, N065, or N191) at global resolutions of 2.88 Å, 2.54 Å, and 2.63 Å, respectively (Figs. 2A–C, Figs. S3–S6). All three BAM compounds localize within the shared intracellular cavity formed at the composite NTSR1–βarr2 protein-protein interface, occuping an overall spatial region analogous to the BAM-binding pocket previously characterized in NTSR1–β-arrestin1 (βarr1) assemblies (Figs. 2E–F) ^29^. Despite this shared binding site, the three modulators differ subtly in their binding depth, three-dimensional orientation, and the spectrum of receptor sub-pocket microdomains (Fig. S7).

**Fig. 2.**
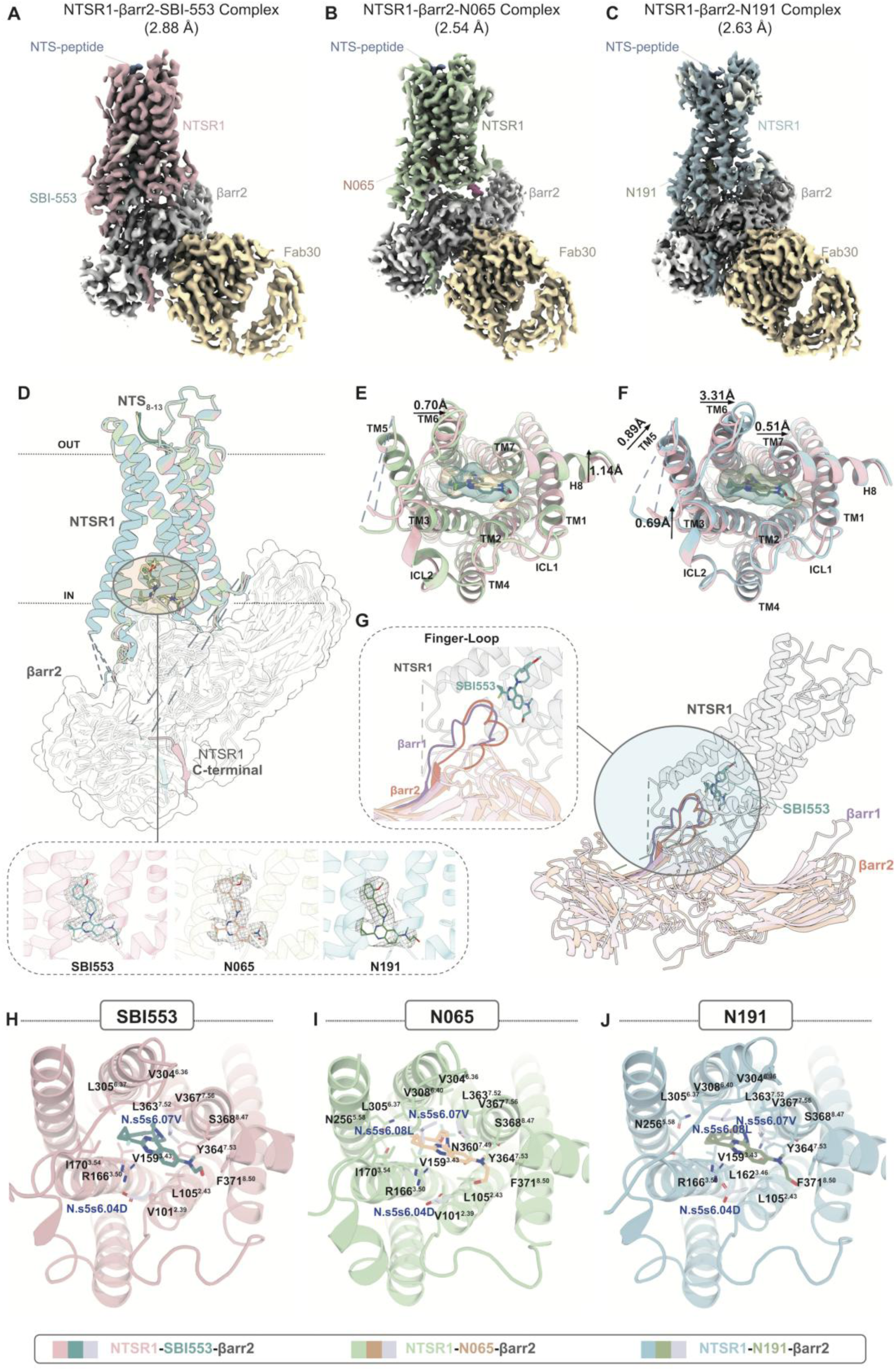
| Cryo-EM structures of NTSR1–β-arrestin2 complexes bound to BAMs. (**A–C**) Cryo-EM density maps of NTSR1–β-arrestin2 complexes bound to SBI-553, N065, and N191 at resolutions of 2.88 Å, 2.54 Å, and 2.63 Å, respectively. NTSR1, βarr2, NTS peptide, Fab30 and the intracellular BAMs are shown in the indicated color scheme. **(D)** Superposition of the three BAM-bound NTSR1–βarr2 complexes after alignment on NTSR1, illustrating the conserved overall receptor–arrestin architecture. Insets show the local cryo-EM density surrounding SBI-553, N065 and N191, respectively, with the ligands shown as sticks and the corresponding density as mesh. **(E–F)** Pairwise structural comparisons of the NTSR1 transmembrane bundle in the SBI-553-bound complex with the N065-bound (E) and N191-bound (F) complexes. Selected local differences in transmembrane-helix positions are indicated in Å. **(G)** Comparison of SBI-553-bound NTSR1–arrestin complexes highlighting differences in arrestin engagement. The enlarged view shows the relative positioning of the βarr1 and βarr2 finger loops at the intracellular receptor interface. **(H–J)** Intracellular binding modes of SBI-553 (H), N065 (I) and N191 (J). Ligands and selected pocket-lining residues are shown as sticks, with NTSR1 displayed as a semi-transparent cartoon.

In each of our three NTSR1–βarr2 reconstructions, βarr2 adopts a canonical core-engagement coupling mode, reminiscent of that observed in previously reported NTSR1–βarr1 structures determined in the presence or absence of SBI-553^28, 29^. This is consistent with the overlapping yet complementary roles of βarr1 and βarr2 *in vivo*. Notably, these BAM-bound NTSR1–βarr2 structures differ markedly from the recently described loop-engagement configuration captured for SBI-553-stabilised NTSR1– βarr1 complexes^29^, with an RMSD greater than 5 Å for the aligned complex (PDBs: 8ZYU and 8ZYY). Whether this difference in arrestin engagement reflects intrinsic differences between βarr1 and βarr2, distinct conformational flexibility of NTSR1, or a combination of both remains to be determined. Despite this difference in arrestin engagement, structural alignments show that the receptor conformations are highly similar across these structures, with an RMSD < 1 Å for all receptor Cα atoms. The main local differences are observed around the intracellular loop (ICL) 1 and ICL3, which may contribute to the distinct binding angles adopted by βarr2 relative to NTSR1 in different BAM-bound states. These structural features illustrate that the endogenous agonist NTS and the synthetic intracellular modulators jointly shape NTSR1 conformational states, resulting in both shared and distinct transducer-binding features.

Despite the presence of different intracellular allosteric partners, the orthosteric peptide NTS maintains a largely conserved overall binding pose across all three ternary complexes, comparable to that observed in BAM-free activated NTSR1 (PDB: 6UP7). Structural comparison, however, reveals modulator-dependent fine-tuning of specific peptide-receptor contact networks (Extended Data Fig. 3A). A set of intermolecular interactions is conserved across all three assemblies. In particular, the C-terminal L13 of NTS (L13^NTS^) forms a salt-bridge with the side-chain of R322^6.54^, alongside a hydrogen-bond donated by the backbone amide of L13^NTS^ toward the side-chain hydroxyl of Y145^3.29^. Notwithstanding these conserved anchor points, subtle positional oscillations are observed for the side-chains of NTS residues R8, R9, I12 as well as for the carboxylate moiety of L13. These modulator-specific changes in the local orthosteric interaction network provide a structural basis for the differential allosteric regulation of NTS binding by distinct BAMs.

Although SBI-553, N065 and N191 share the same intracellular binding cavity, they adopt subtly distinct binding poses and establish different contact footprints across pocket sub-regions. Rather than forming direct contacts in BAM-free NTSR1 active structures^28^, R166^3.50^ and Y364^7.53^ reside adjacent to the bound BAM modulators and, together with a set of shared interactions involving residues from the central intracellular cavity, define a common anchoring environment for BAM recognition (Figs. 2G–I; Extended Data Fig. 3F). Beyond this shared framework, the three ligands display distinct spatial engagement of the cavity. Relative to the SBI-553-bound NTSR1–βarr1 complex^29^, SBI-553 inserts deeper into the TM3-TM7 cleft, and approaches the βarr2 finger loop, making extensive contacts with both receptor residues and βarr2 V71 and L72, thereby directly spanning the receptor–βarr2 interface. In comparison, N191 similarly contacts with L305^6.37^, V308^6.40^, Y364^7.53^, V367^7.56^, S368^8.47^ and F371^8.50^, while retaining interactions with the βarr2 finger loop. Uniquely among the three analogues, N065 shows a distinct feature in which its fluorinated substituent projects toward N360^7.49^ on the TM7 side of the pocket and form a short-range ligand-specific halogen bond that is not observed for the other two modulators (Extended Data Fig. 3G). Collectively, these structures show that subtle chemical changes within a common BAM scaffold redistribute ligand occupancy across distinct microdomains of the intracellular cavity. In combination with the modulator-dependent remodeling of the orthosteric NTS interaction network, these differences illustrate how chemically related intracellular modulators can remodel both sides of the receptor allosteric network while preserving the overall NTSR1–βarr2 architecture.

### Potential hotspots around the allosteric pocket reshape transducer output

To reveal how the intracellular allosteric pocket of NTSR1 shapes signal transduction, we performed alanine scanning across this region together with key activation motifs and evaluated the resulting phenotypes at three levels: orthosteric activation, intrinsic activity of the allosteric ligands, and orthosteric-allosteric cooperativity (Figs. S8–S21). We treated NTS and intracellular modulators as conformational probes that bias NTSR1 into distinct ligand-conditioned states. Distinct intracellular receptor states expose differential transducer-sensitive nodes. Among expression-competent mutants, alanine scanning identified a limited set of transducer-sensitive hotspots clustered around the intracellular allosteric pocket and adjacent activation network (Figs. 3A-B). Mapping these effects across pathways uncovered spatially separated microdomains that contribute unequally to different downstream signaling outputs (Figs. 3C-D). We found that residues H102^2.40^, I170^3.54^, I259^5.61^, G301^6.33^, I359^7.48^ emerged as functional hotspots across multiple signaling branches (Fig. 3C). Notably, alanine substitution at these positions produced bidirectional effects in concentration-response profiles rather than uniform loss of function (Fig. 3D, Figs. S8–S12). Because alanine substitutions more commonly weaken local interactions or packing, such enhanced or directionally divergent responses were particularly informative, as they are less readily explained by nonspecific structural disruption and instead suggest release or redistribution of local conformational constraints that alter transducer compatibility. At the pathway level, gain-of-function effects in the G_i_ branch were mapped to I259^5.61^, G301^6.33^ and W316^6.48^, whereas enhanced G_13_ output was primarily associated with H102^2.40^, G301^6.33^ and W316^6.48^. By contrast, increased β-arrestin2 recruitment and G_q_ activation was more selectively linked to I170^3.54^ and I359^7.48^, respectively (Fig. 3D). Thus, different signaling branches were preferentially sensitized by partially overlapping but non-identical residue sets, identifying them as pathway-selective determinants of transducer coupling.

**Fig. 3.**
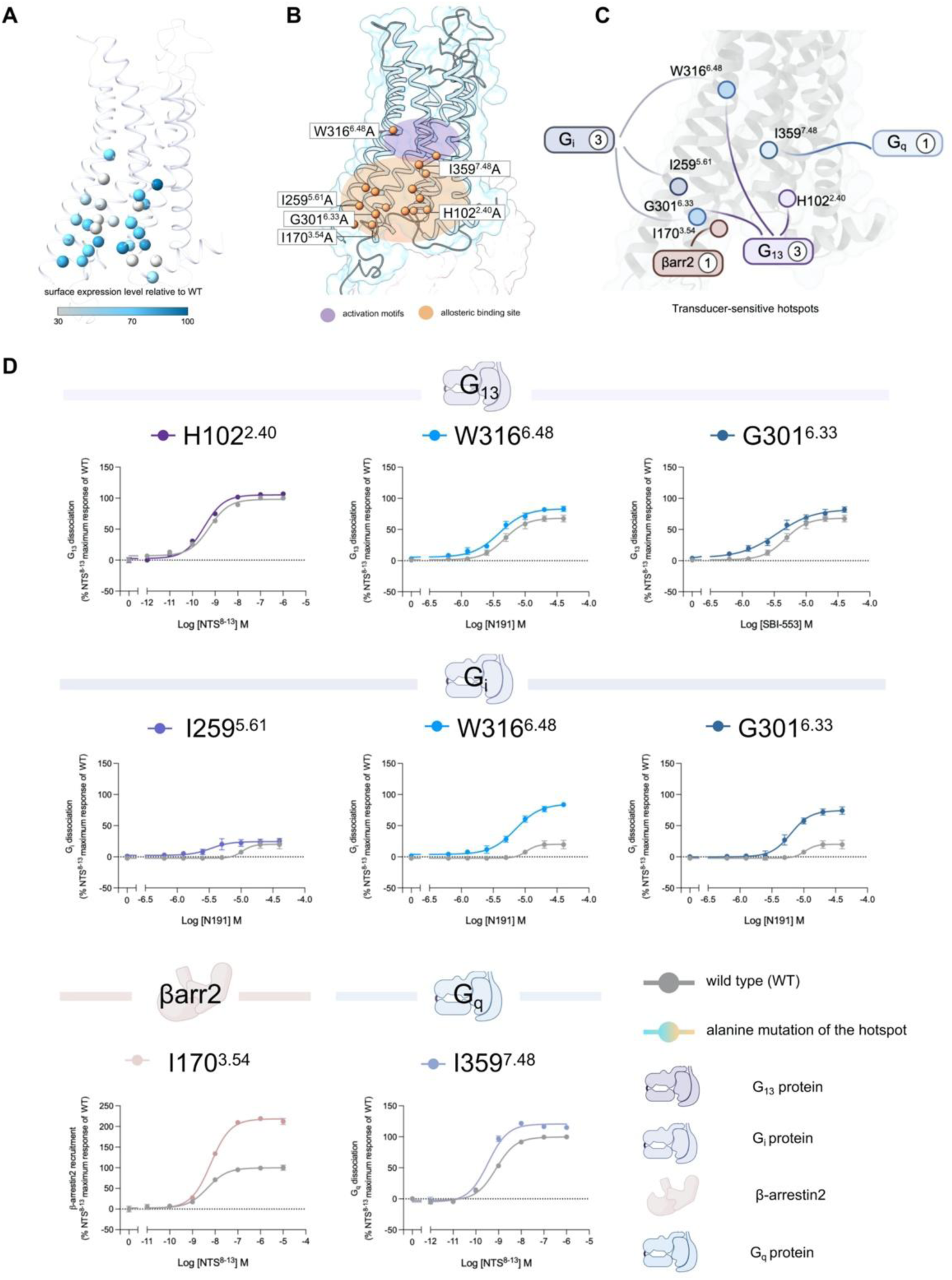
| Alanine scanning-based mapping of transducer-sensitive hotspots around the intracellular allosteric pocket of NTSR1. **(A)** Relative cell-surface expression of alanine mutants mapped onto the NTSR1 structure. Residues are shown as spheres and colored according to surface expression relative to wild type (WT), as indicated by the color scale. **(B)** Structural localization of representative residues selected for functional analysis. The intracellular allosteric binding site is shown in orange, and adjacent conserved activation motifs are shown in purple. Labeled residues indicate potential hotspots. **(C)** Schematic summary of pathway-associated residues on the intracellular face of NTSR1. Circles indicate representative residues associated with G_i_, G_13_, G_q_, or β-arrestin2 signaling phenotypes. Numbers indicate the number of residues assigned to each signaling branch in this summary. **(D)** Representative concentration-response curves for selected alanine mutants in different signaling assays, organized by transducer class. Colored curves represent the indicated alanine mutant and gray curves represent WT. For each residue, the panels show representative assay conditions in which the mutation enhanced relative to WT. G_13_-related responses are shown for H102^2.40^, W316^6.48^ and G301^6.33^. G_i_-related responses are shown for I259^5.61^, W316^6.48^ and G301^6.33^. β-arrestin2-related responses are shown for I170^3.54^; and G_q_-related responses are shown for I359^7.48^. Responses are plotted as percentages of the maximal WT response of NTS^8–13^. Data are presented as mean ± s.e.m. from three independent experiments.

Structurally, these hotspots cluster into several spatially separable regions surrounding the intracellular pocket. Residues H102^2.40^ and I170^3.54^ are located on the TM2-TM3 side near the upper rim, I259^5.61^ lines the intracellular face of TM5, G301^6.33^ sits within the shallow TM6-side transducer-facing interface and is strongly associated with G-protein outputs, and I359^7.48^ lies near the NPxxY region of TM7 and links the activation network to the upper edge of the intracellular pocket. Interestingly, mutation of the canonical toggle switch W316^6.48^, which located outside the allosteric ligand-binding pocket, weakened orthosteric ligand activation while enhancing agonism by the intracellular ligands (Fig. 3D). These bidirectional effects, particularly at W316^6.48^, raise the possibility that BAMs differentially stabilize local microstates of conserved NTSR1 activation motifs. Such microstate tuning could alter the relative compatibility of the receptor with distinct transducers and thereby contribute to signaling bias, in addition to direct remodeling of the receptor–transducer interface. Consistent with this view, the allosteric effects of all three ligands were extensively reprogrammed across the mutant panel. For each pathway, mutant responses could be classified relative to wild type as weakened modulation, enhanced modulation, mode-shifted regulation, or reversal of modulation direction (Figs. S13–S21). This reassignment was strongly ligand– and pathway-dependent. Although most mutations attenuated allosteric effects, specific residue contexts enabled the same ligand to switch its mode of action, with reversals occurring preferentially at core activation motifs such as W^6.48^ and D^2.50^, and nearby residues such as N^2.56^.

### *In vivo* physiological effects of N065 and SBI-553 on NTSR1-related functions

Given their divergent signaling profiles, we then investigated whether N065 and SBI-553 produce distinct *in vivo* effects on NTSR1-related physiological functions, including core body temperature, sedation, motor activity and coordination, cardiovascular parameters, and energy metabolism^30–33^ (Fig. 4A).

**Fig. 4.**
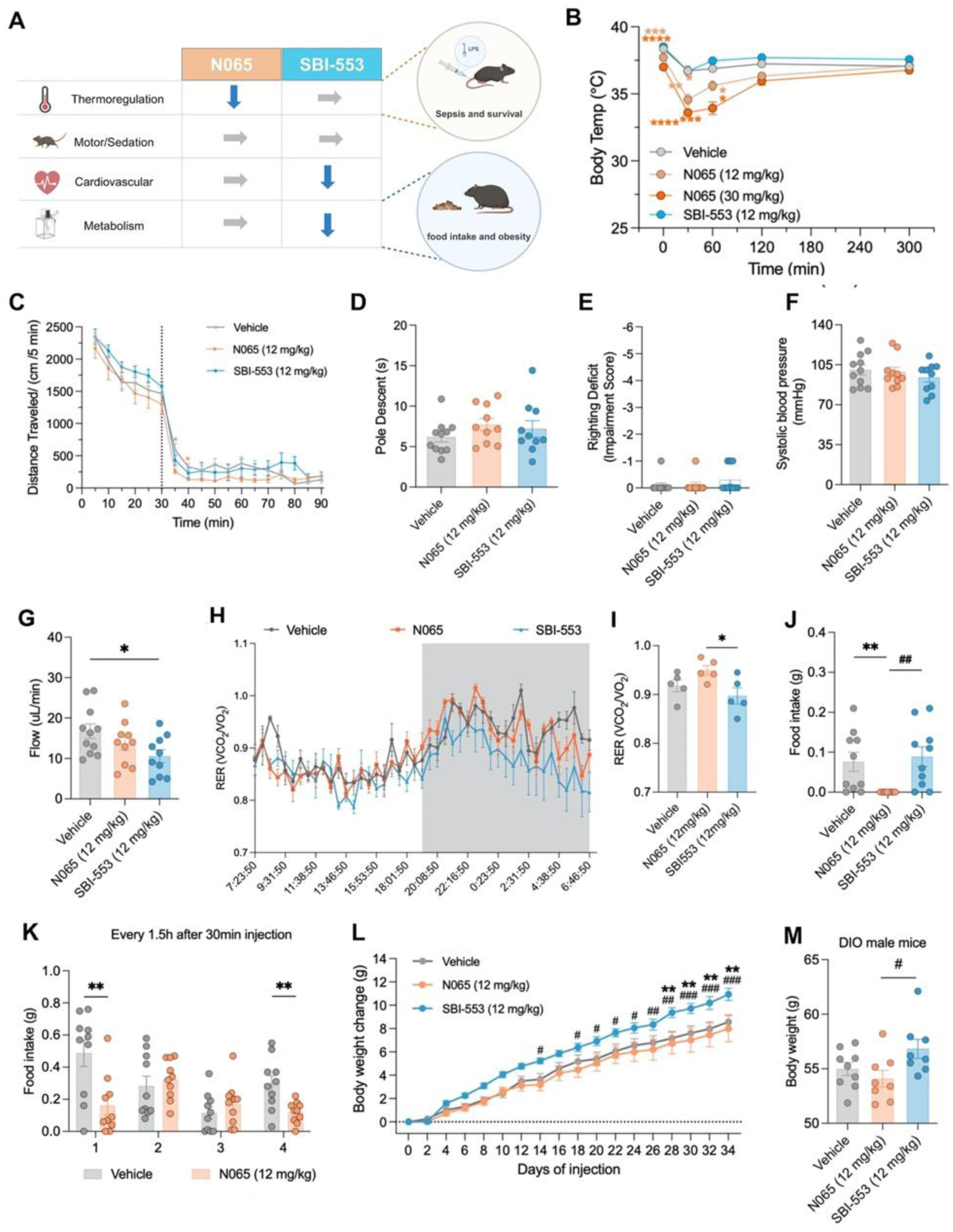
| Effects of N065 and SBI-553 on Basic Physiological Phenotypes in mice. **(A)** Summary of differences between N065 and SBI-553 in NTSR1-related phenotypes. **(B)** Core body temperature. Animals received vehicle, N065 or SBI-553 (i.p.) at Time 0. **(C)** Open field test. Animals were placed into the open field for 30 min prior to administration of either vehicle, N065 or SBI-553. Distance moved in open field test was recorded. **(D)** Pole descent test. Time required for the mouse to descend from the pole to the home cage. Animals received vehicle, N065 or SBI-553 (i.p.) 30 min prior to testing. **(E)** Righting reflex. Contact righting deficits were scored on a six-point scale. Animals received vehicle, N065 or SBI-553 (i.p.) 30 min prior to testing. **(F-G)** Systolic blood pressure (mmHg) (F), blood flow (μL/min) (G) was continuously recorded by tail-cuff system CODA in mice 30 min after treatment with vehicle, N065 or SBI-553 (i.p.). Baseline was set to 0. **(H-I)** Metabolic cage analysis of mice treated with vehicle, N065 or SBI-553. Respiratory exchange ratio (H) and average RER (I). For details on statistical comparisons, see Table S1. **(J-K)** Regular food intake in C57BL/6 mice were evaluated under short-term fasted (6 h, J), long-term fasted (17 h, K) states. Mice were administered N065 (12 mg/kg), SBI-553 (12 mg/kg), or vehicle 30 min before the feeding test. In fasting studies, drug administration occurred at 5.5 h (short-term, J) or 16.5 h (long-term, K) into the fast. Food intake was measured for 1 hour in short-term fasted mice (J), and for 6 hours in fed and long-term fasted mice (K). **(L)** Body weight change in DIO mice fed a high-fat diet treated with vehicle, N065 or SBI-553. DIO mice were treated daily with vehicle, N065, SBI-553. Body weight was recorded daily. **(M)** Body weight of DIO mice treated with vehicle, N065 or SBI-553 from day 34 of HFD diet. For details on statistical comparisons, see Table S1 and Table S2.

To assess whether these divergent *in vivo* effects could be attributed simply to differences in drug exposure, we compared the pharmacokinetic and brain-penetration profiles of N065 and SBI-553. Following intraperitoneal administration at 12 mg/kg, the two compounds reached similar peak plasma concentrations, although N065 showed greater overall plasma exposure, whereas SBI-553 exhibited substantially higher brain exposure and brain-to-plasma ratios (Fig. S22). Thus, the distinct physiological effects of N065 and SBI-553 do not track simply with systemic or brain exposure, arguing against pharmacokinetic differences as the sole explanation and supporting a contribution from their distinct allosteric pharmacology.

In mice model, N065 (12 mg/kg, 30 mg/kg, i.p.) produced a rapid (within approximately 30 min) and sustained (lasting at least 2 h) hypothermia, whereas SBI-553 (12 mg/kg, i.p.) had no effect on core body temperature (Fig. 4B). Since NTSR1-mediated hypothermia is thought to be primarily driven by the G_q_-PLC-IP_3_/Ca²⁺ signaling axis^30^, and neither N065 nor SBI-553 directly activates G_q_, this divergence likely reflects their differential allosteric effects on NTS-evoked G_q_ signaling (Fig. 1E). Although both compounds enhance NTS binding, SBI-553 acts as a potent G_q_ NAM that markedly reduces NTS potency and efficacy, whereas N065 is a weaker G_q_ NAM that largely preserves NTS-driven G_q_ activation (Fig. 1E, Extended Data Fig. 2B). Consequently, only N065 permits endogenous NTS-evoked G_q_ signaling to surpass the threshold required for hypothermia.

Across distinct behavioral tests, neither N065 or SBI-553 affected open-field locomotion, pole descent latency, wire hang duration, or righting reflex response time (Figs. 4C–E, Fig. S23A), indicating the absence of motor, muscle coordination, or sedative effects. Furthermore, no significant differences in diastolic or systolic blood pressure, heart rate, or blood flow volume were observed between N065– and SBI-553-treated mice. However, SBI-553-treated male mice exhibited a selective reduction in blood flow velocity, suggesting a compound-specific cardiovascular effect (Figs. 4F– G, Figs. S23C–D). Given the critical role of NTSR1 in regulating energy balance^34^, we next assessed energy metabolism using metabolic cages. The results showed that, compared with control mice, N065 treatment did not significantly alter oxygen consumption, carbon dioxide (CO_2_) production, or heat generation (Fig. 4H, Figs. S23E–G). In contrast, SBI-553 significantly reduced the respiratory exchange ratio (RER), indicating a shift in energy substrate utilization (Fig. 4I). Since N065 can regulate feeding behavior, we further investigated whether it could exert anti-obesity effects in obese mice. Diet-induced obese (DIO) mice were treated daily with N065 or SBI-553 for one month. Interestingly, N065 did not reduce weight gain, but SBI-553-treated DIO mice exhibited significantly accelerated weight gain and a trend toward increased fat mass (Figs. 4L–M, Figs. S23I–K), suggesting that SBI-553 may promote adiposity in the context of diet-induced obesity.

Given that NTSR1 is present in memory-relevant areas such as the PFC and anterior cingulate^35^, we assessed cognitive and emotional behaviors in mice under SBI-553 or N065 treatment. SBI-553 impaired performance in the Morris water maze and reduced spontaneous alternation in the Y-maze, indicating compromised working memory, whereas N065 showed minimal effects (Fig. S24A–C). In the novel object recognition test, neither compound altered the discrimination index (Fig. S24D). Since NTSR1 is also linked to emotional behaviors^36^, we then assessed anxiety– and depression-like phenotypes. In the elevated-O-maze and forced swim test, neither N065 nor SBI-553 significantly affected anxiety– and depression-related behaviors (Fig. S24A–C). In the novelty-suppressed feeding test, N065 significantly reduced food consumption relative to SBI-553 without affecting latency to approach food (Fig. S24F–G), indicating that N065-induced hypophagia is not associated with emotional state (Fig. S24A–C). Consistent with this, N065 robustly suppressed food intake under multiple conditions. It reduced regular food intake in short-term and long-term fasted mice, as well as in ad libitum-fed animals (Figs. 4J–K, Fig. S23H). Together, these results demonstrate that intracellular allosteric modulators with distinct signaling biases translate into divergent physiological outcomes *in vivo*, with N065 selectively engaging thermoregulatory and anorectic pathways, and SBI-553 exhibiting distinct metabolic and cognitive liabilities.

### N065 improves survival in a murine endotoxemia model

Because NTSR1-mediated hypothermia is primarily linked to G_q_ signaling^30^, we first compared N065 and SBI-553 in counteracting hypothermia induced by the NTSR1 agonist PD149163. Mice were pretreated with either compound or their respective vehicles before PD149163 administration (Fig. 5A). PD149163 induced rapid (within 60 min) and sustained (≥8 h) hypothermia, and both compounds attenuated this response at 12 mg/kg, with greater alleviation at 30 mg/kg (Fig. 5B). SBI-553 promoted temperature recovery more effectively than N065 at both doses, consistent with its stronger G_q_ NAM activity in vitro.

**Fig. 5.**
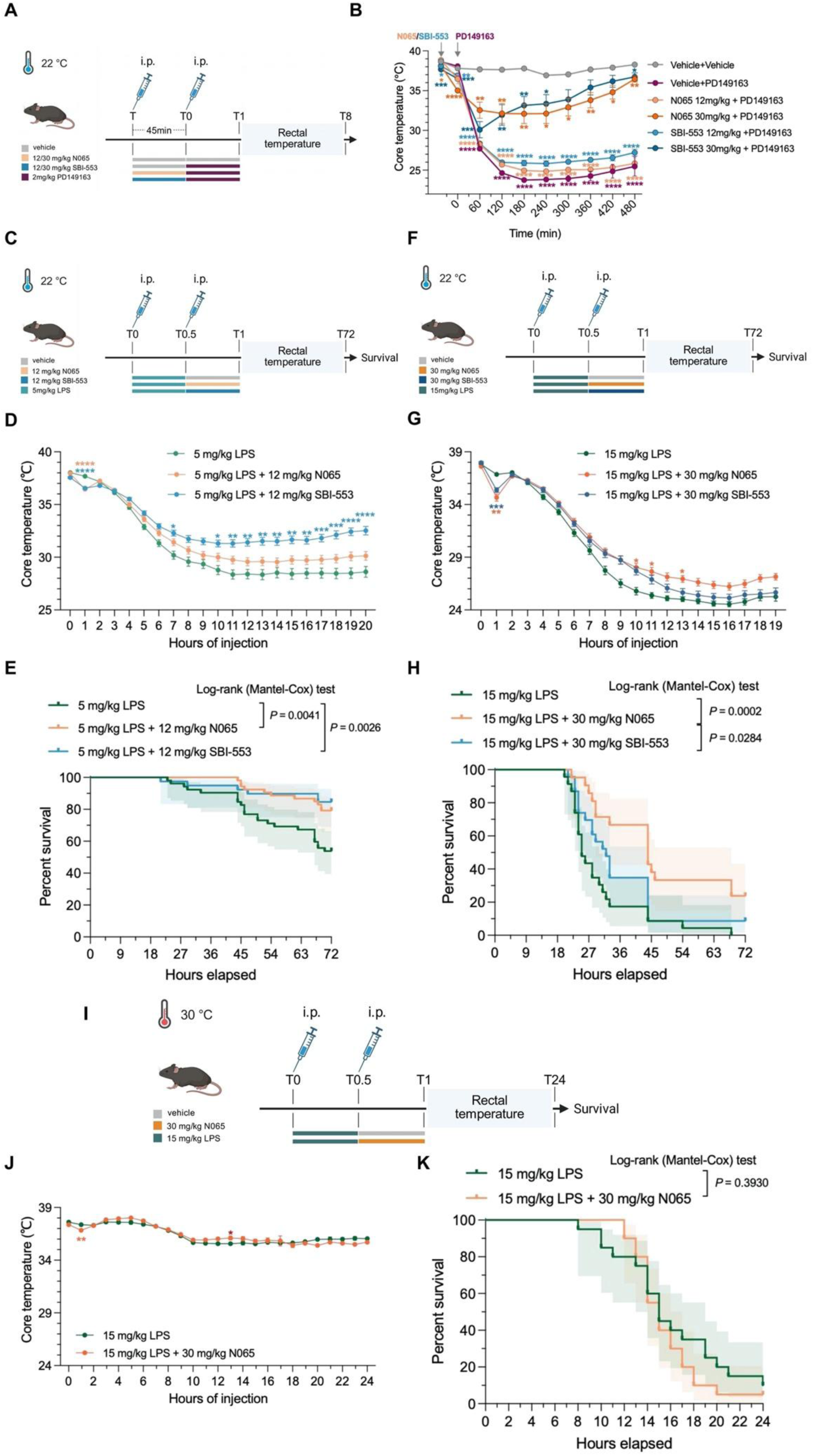
| N065 induced pharmacological hypothermia and improved mortality. **(A-B)** Effects of N065 and SBI-553 on PD149163-induced hypothermia. (A) Animals received vehicle 1 (5% DMSO + 40% PEG300 + 5% Tween 80 + 50% saline, i.p.), N065 (12 or 30 mg/kg, i.p.) or SBI-553 (12 or 30 mg/kg, i.p.) 45 min prior to the study (gray arrow). Animals received vehicle 2 (0.2% DMSO in saline, i.p.) or PD149163 (2 mg/kg, i.p.) at Time 0. Animals remained at 22℃ the whole experiment and were observed over a period of up to 8 h. (B) Core body temperature was monitored at multiple time-points. **(C-E)** Effects of N065 and SBI-553 on survival rate reduction induced by low-dose LPS. (C) Animals received LPS (5 mg/kg, i.p.) at Time 0. Animals received vehicle, N065 (12 mg/kg, i.p.) or SBI-553 (12 mg/kg, i.p.) 30 min after LPS injection. Animals remained at 22℃ the whole experiment and were observed over a period of up to 72 h. (D) Core body temperature was monitored at multiple time-points. (E) Kaplan-Meier cumulative survival plots. Data were analyzed using a log-rank (Mantel-Cox) test with results indicated on a graph. **(F-H)** Effects of N065 and SBI-553 on survival rate reduction induced by high-dose LPS. (F) Animals received LPS (15 mg/kg, i.p.) at Time 0. Animals received vehicle, N065 (30 mg/kg, i.p.) or SBI-553 (30 mg/kg, i.p.) 30 min after LPS injection. Animals remained at 22℃ the whole experiment and were observed over a period of up to 72 h. (G) Core body temperature was monitored at multiple time-points. (H) Kaplan-Meier cumulative survival plots. Data were analyzed using a log-rank (Mantel-Cox) test with results indicated on a graph. **(I-K)** Effects of N065 on survival rate reduction induced by high-dose LPS at 30℃. (I) Animals received LPS (15 mg/kg, i.p.) at Time 0. Animals received vehicle, N065 (30 mg/kg, i.p.) or SBI-553 (30 mg/kg, i.p.) 30 min after LPS injection. The mice were switched to 30℃ immediately after LPS and observed over a period of up to 24 h. (J) Core body temperature was monitored at multiple time-points. (K) Kaplan-Meier cumulative survival plots. Data were analyzed using a log-rank (Mantel-Cox) test with results indicated on a graph. Data are presented as mean core temperature ± SEM. For details on statistical comparisons, see Table S3.

LPS-induced hypothermia in mice is partly mediated by neurotensin-NTSR1 signaling downstream of peripheral TLR4 activation, and pharmacological induction of hypothermia has been shown to improve survival in endotoxemia models^37^. We therefore tested whether N065-induced hypothermia confers protection against LPS challenge. In the low-dose model (LPS 5 mg/kg, compound 12 mg/kg), both N065 and SBI-553 augmented the LPS-induced temperature drop at 30 min post-injection (Fig. 5D). By 10 h, SBI-553-treated mice showed significantly higher body temperatures than LPS-only controls, while N065-treated mice showed a non-significant trend toward recovery. Survival analysis revealed that although both compounds reduced LPS-induced mortality, N065 more effectively delayed time to death (Fig. 5E). In the lethal-dose model (LPS 15 mg/kg, compound 30 mg/kg), both compounds again augmented the early hypothermic response (Fig. 5G). However, by 10 h, N065-treated mice maintained significantly higher body temperatures than LPS-only controls, whereas SBI-553-treated mice showed no significant difference from the LPS group. Critically, survival analysis demonstrated that N065 significantly improved survival, whereas SBI-553 had no significant effect on lethal-dose LPS-induced mortality (Fig. 5H). To determine whether the survival benefit of N065 was mediated by its hypothermic effect, mice were housed at 30°C following lethal-dose LPS injection to attenuate N065-induced hypothermia. Under thermoneutral conditions, both the hypothermic effect of N065 and its survival benefit were abolished (Figs. 5I–K), directly implicating hypothermia as the primary protective mechanism. Together, these findings demonstrate that N065 independently induces hypothermia, confers protection in a lethal endotoxemia model by lowering body temperature, and prolongs survival time of the model mice, effects not shared by SBI-553.

### Distinct intracellular allosteric modulators generate divergent phospho-signaling states

The divergent physiological profiles of allosteric modulators prompted us to examine whether fine-tuning the intracellular allosteric pocket is also accompanied by differences in downstream signaling states. To address this question in a receptor-defined cellular context, we profiled protein phosphorylation in NTSR1-expressing HEK293T cells. We selected a 1-h stimulation period to capture a downstream phosphorylation state established after sustained ligand exposure rather than rapid receptor-proximal events. A phospho-kinase array showed that SBI-553 alone did not activate ERK but attenuated NTS-induced ERK phosphorylation, consistent with previous findings^2^, while also altering a subset of other NTS-responsive phosphorylation events (Extended Data Figs. 4A–B).

We next performed phosphoproteomic profiling following treatment with NTS, each intracellular modulator alone, or NTS in combination with SBI-553, N065 or N191 (Extended Data Fig. 4C). Biological replicates showed good within-group concordance and comparable coefficient-of-variation distributions across conditions (Extended Data Fig. 4D, Fig. S25A). Focusing on the biologically relevant NTS background, all three modulators altered subsets of NTS-regulated phosphosites, with both shared and ligand-specific responses (Extended Data Figs. 4 E–G). Pairwise comparisons under the NTS background further identified phosphosites that distinguished N065 and N191 from SBI-553.

Kinase-substrate enrichment analysis (KSEA) further revealed that these differences propagate to kinase activity networks. Compared with SBI-553, N065 produced broader negative kinase enrichment under NTS stimulation, including kinase activities associated with MAPK, mTOR, and GSK3 signaling (Extended Data Fig. 4H, Fig. S25D). In contrast, GRK2 (ADRBK1), a kinase closely associated with β-arrestin recruitment, showed positive enrichment with all three allosteric ligands, though less strongly with N065 than SBI-553 or N191. KEGG analysis further identified differences in pathways including mTOR and PI3K–Akt signaling, nucleocytoplasmic transport, cellular senescence, and cytoskeleton-related processes (Extended Data Fig. 4I, Figs. S25E–H). These data show that intracellular allosteric modulators produce partially overlapping but distinguishable phosphoproteomic and kinase-enrichment profiles under a common NTS stimulus. These differences provide a broader signaling context for the distinct pharmacological and physiological activities of the modulators.

## Discussion

Biased signaling at GPCRs offers a strategy for selectively modulating therapeutically relevant pathways while limiting undesired signaling. Intracellular allosteric sites are particularly attractive because they lie close to the receptor–transducer interface and can therefore influence coupling without competing directly with endogenous agonists. Previous work on SBI-553 established that modifications of this scaffold can alter G-protein subtype selectivity through molecular-glue or molecular-bumper mechanisms at the transducer interface. Here, we extend this concept by chemically perturbing distinct microdomains across the same intracellular pocket and profiling their effects across multiple G-protein and β-arrestin pathways. Our results show that relatively small, spatially distributed modifications within one pocket are sufficient to generate distinct transducer-response profiles, indicating that the NTSR1 intracellular cavity behaves as a spatially tunable regulatory landscape rather than a single uniform allosteric site.

The cryo-EM structures provide a structural framework for this chemical sensitivity. SBI-553, N065 and N191 occupy the same intracellular cavity but differ in binding depth, orientation and the sub-pocket regions they engage, while the overall NTSR1– βarr2 architecture remains highly similar. Thus, the observed signaling differences are more consistent with local remodeling of the intracellular cavity and receptor– transducer interface than with formation of globally distinct receptor states. The orthosteric peptide NTS also retains a largely conserved binding pose, with BAM-dependent differences confined mainly to local peptide–receptor contacts. These observations suggest that intracellular bias can arise not only from direct molecular-glue or molecular-bumper effects at the effector interface, but also from how different regions of the intracellular pocket are selectively engaged and locally remodeled.

The alanine-scanning data further support functional heterogeneity within this intracellular network. Several mutations produced pathway-dependent gain-of-function or bidirectional effects rather than uniform loss of signaling, identifying residues whose local structural constraints influence transducer compatibility. Importantly, these data do not establish redistribution of the global receptor conformational ensemble, but instead identify transducer-sensitive positions and provide a testable model for how local receptor microstates may participate in biased signaling.

The *in vivo* studies further demonstrate that closely related intracellular modulators can produce distinct physiological profiles. N065 and SBI-553 differed in thermoregulation, feeding, metabolic and cognitive readouts, whereas their pharmacokinetic profiles indicate that these differences do not simply track systemic exposure or brain penetration. N065 showed greater overall plasma exposure, while SBI-553 achieved substantially higher brain exposure, arguing against pharmacokinetic differences as the sole explanation for their divergent effects. In the endotoxemia model, N065 induced hypothermia and improved survival following lethal LPS challenge, and this protection was abolished under thermoneutral conditions that attenuated hypothermia. These findings support hypothermia as an important mediator of protection, while avoiding a direct causal assignment of survival to any single proximal signaling parameter. The PK data are consistent with the physiological differences reflecting, at least in part, distinct allosteric pharmacology rather than exposure alone.

Finally, phosphoproteomic profiling showed that the three BAMs produce distinct phosphorylation and kinase-associated signaling states under a common NTS-stimulated background. Because these measurements were obtained at a single 1-h time point in a heterologous system, they are best interpreted as sustained downstream signaling states rather than as a temporal map of receptor-proximal events. They also do not establish a causal relationship between the observed phosphorylation changes and the physiological phenotypes in mice. Together, our findings define the NTSR1 intracellular allosteric pocket as a chemically and functionally heterogeneous regulatory landscape in which local chemical perturbations can differentially shape transducer coupling, local receptor microstates and downstream signaling. This pocket-centered view provides a framework for designing intracellular GPCR modulators with more precisely differentiated signaling properties, while emphasizing that the relationships among proximal bias, downstream signaling and physiological outcome must be established in each biological context.

## Methods

### Chemistry

All commercial reagents and solvents were used directly without further purification, unless otherwise stated. All reactions were conducted in round-bottomed flasks equipped with Teflon-coated magnetic stirring bars. Experiments involving air-sensitive compounds were performed under a nitrogen atmosphere. NMR spectra (¹H and ¹³C) were recorded on a Bruker Avance 400 MHz spectrometer using deuterated dimethyl sulfoxide (DMSO-d6) as the solvent. The synthesized compounds were purified by silica gel flash chromatography, eluting with ethyl acetate/petroleum ether mixtures. LCMS analyses were performed on a Waters 2695 system, and high-resolution mass spectrometry (HRMS) was conducted on a Sciex X500B QTOF system equipped with an electrospray ionization (ESI) source. The purity of all final compounds was confirmed by high-performance liquid chromatography (HPLC) on an Agilent 1260 system equipped with a UV detector (254 nm) and a reversed-phase C18 column (4.6 × 150 mm, 5 μm). A gradient elution of acetonitrile and water (containing 0.1% trifluoroacetic acid) was used as the mobile phase at a flow rate of 1.0 mL/min, and the column temperature was maintained at ambient temperature.

### Tango assay

Human NTSR1 (1–418) was cloned into pcDNA6 vector consisting of an expression cassette with tobacco etch virus (TEV) protease cleavage site and the transcriptional activator tTA at the C terminus. A TEV protease cDNA was fused to the N terminus of ARRB1 (1-418) or ARRB2 (1-409). Interaction between NTSR1 and β-arrestins leads to the cleavage of the TEV site, thus releasing tTA to trigger tTA-dependent luciferase reporter gene expression. HTL cells were cultured in 6-cm dishes at a density of 1.2×10^6^ cells/dish for 24 h, and then transfected with 1 µg NTSR1-tTA and 1 µg TEV-ARRB1/2 expression plasmids using the ExFect Transfection Reagent (Vazyme). After transfection for 24 h, cells were incubated overnight with PBS (vehicle) or different concentrations of ligands. Then luciferase activities were evaluated according to manufacturer’s protocols of the Bright-Lumi Luciferase Kit (Beyotime). Experiments were performed three times and data were presented as mean ± s.e.m.

### BRET assay for G protein dissociation and β-arrestin recruitment

HEK293T cells were cultured in DMEM supplemented with 10% FBS. Cells were seeded at a density of 3–4 million cells per dish in 10-cm dishes. After 18-24 h, cells were transfected at a confluency of 70–80%, using a 1:1:1:1 DNA ratio of NTSR1: Gα-RLuc8: Gβ: Gγ-GFP2 (2 µg per construct for 10-cm dishes, the plasmids encoding subunits of the heterotrimeric G protein complex named TRUPATH^38^ were provided by Bryan Roth laboratory, Addgene kit #1000000163), or using a 1:1:10 DNA ratio of NTSR1-RLuc8: GRK2: GFP2-β-arrestin (total 8 µg for 10-cm dishes). ExFect Transfection Reagent (Vazyme) was used to complex the DNA at a ratio of 2 µl of ExFect per µg of DNA, Opti-MEM (Gibco) at a concentration of 10 ng of DNA per µl of Opti-MEM. The next day, cells were reseeded in white opaque-bottom 96-well assay plates (Beyotime) at a density of 30,000–50,000 cells per well. 24 hours later, medium was replaced with 80 µl assay buffer (HBSS + 20 mM HEPES, pH 7.4). After 2 min equilibration, cells were stimulated with 20 µl compounds (serial dilutions) for 5 min.

After addition of 5 µM coelenterazine 400a (Nanolight Technology), the plates were read on a Synergy Neo plate reader (BioTek) using 410-nm (RLuc8-coelenterazine 400a) and 515-nm (GFP2) emission filters. BRET2 ratios were calculated as GFP2/Rluc8, and signals were normalized to the reference ligand NTS^8–13^ (0% at baseline; 100% at maximal NTS^8–13^ response). Data are presented as mean ± s.e.m. from three independent experiments. For direct agonism analyses, data were analyzed in GraphPad Prism 10 by nonlinear regression using a four-parameter dose-response model. For allosteric modulation studies, dose–response curves were fitted to the Black-Leff-Ehlert’s allosteric operational model^39, 40^ in Prism 10. To more clearly present the overall effects of these allosteric ligands, we applied operational model analysis^41–43^ and extracted Δlog(*τ*⁄*KA*) for each pathway as a quantitative index.

### NanoBRET binding assay

TAMRA-PEG6-NTS^8–13^ was synthesized by TGpeptide. To verify specificity, HEK293T cells with or without NTSR1 expression were incubated with 1 µM fluorescent ligand for 5 min, washed with PBS, and imaged on a fluorescent cell imager (ZOE, Bio-Rad) using bright-field and red fluorescence channels. For NanoBRET binding, HEK293T cells were transiently transfected with NanoLuc-NTSR1 (1 µg plasmid per 6-cm dish; 1.2×10^6^ cells) and assayed 36 h post-transfection. Cells were resuspended in BRET assay buffer and plated into white opaque 96-well plates (Beyotime) at 80 µL per well. For saturation binding, 10 µL serial dilutions of fluorescent ligand and 10 µL vehicle or allosteric modulators were added, then incubated for 30 min at 37 °C. 20 µL furimazine (Macklin) was then added (final 10 µM) and luminescence at 460 nm and 580 nm was measured on an Envision plate reader (PerkinElmer). NanoBRET ratios were calculated as 580 nm/460 nm and multiplied by 1000 to yield mBRET. Data were fitted to a one site binding model in GraphPad Prism 10. For competition binding, 100 nM fluorescent ligand was incubated with serial dilutions of the unlabeled NTS^8–13^ (20 µL) for 30 min at 37 °C, followed by furimazine addition and signal acquisition as above. Data were fitted to the one site-Fit logIC_50_ model in Prism 10. Data are presented as mean ± s.e.m. from three independent experiments.

### Alanine mutation scanning and membrane receptor expression detection

Human NTSR1 (1-418) was cloned into a modified pcDNA3.1 vector with a HiBiT tag (Promega) at the N-terminus. NTSR1 mutations were generated through site-directed mutagenesis using the Mut Express II Fast Mutagenesis Kit V2 (Vazyme). The successful introduction of the mutations in the polymerase chain reaction products was confirmed through DNA sequencing. The receptor surface expression levels were detected by the Nano-Glo HiBiT Extracellular Detection System (Promega). N-terminal HiBiT-tagged receptors (500 ng/well) were transfected into HEK293T cells seeded in a 12-well plate. After 36 h, the cells were reseeded into white opaque-bottom 96-well plates in HBSS (15,000 cells in 30 µl/well). Then, 30 µl of assay buffer (dilute the LgBiT protein 1:100 and the substrate 1:50 into an appropriate volume) was added to each well and incubated for 10 min. Luminescence counts were determined using the Synergy Neo plate reader (BioTek). The expression level was presented as a percentage relative to that of the wild-type receptor. Mutants with membrane expression levels no less than 65% of WT were used in subsequent experiments. The transfection dose of the mutants was adjusted according to the membrane expression level to achieve the WT expression level.

### Construction and Expression of the NTSR1-Arrestin Complex

The NTSR1-arrestin complex was expressed and purified using a modified version of a previously described method^44^. Briefly, a chimeric construct was created in the pFastBac1 baculovirus expression vector for insect cell expression. This construct encoded a fusion protein composed of the following elements, in order: an N-terminal hemagglutinin (HA) signal sequence followed by a cytochrome b562 RIL domain (BRIL) fused to the N-terminus of human neurotensin receptor 1 (NTSR1; UniProtKB accession: P30989, residues 49-418); a truncated human β-arrestin2 (βarr2-ΔCT; UniProtKB accession: P32121, residues 1-393, with L278F, S280A, C242V and C251V mutations) linked to the C-terminus of NTSR1 via a short GSA linker; and the light chain of the Fab30 antibody (Fab30L) fused to the C-terminus of βarr2-ΔCT using a longer flexible linker (GSAGSAGSAGSA). This BRIL-NTSR1-βarr2-ΔCT-Fab30L fusion protein was employed to facilitate stable complex formation. The Fab30 heavy chain was separately cloned into a pFastBac1 vector and modified to include a C-terminal octahistidine (8 × His) tag. For receptor phosphorylation, two additional constructs were created in separate pFastBac1 vectors: a modified version of human GRK5 (UniProtKB accession: P34947) carrying the Q41L, K454A, and R455A mutations, and mouse PIP5K (UniProtKB accession: P70181). All genes were codon-optimized for insect cell expression (Genewiz). Mutations were introduced via PCR, and the sequences of all constructs were confirmed by DNA sequencing.

The BRIL-NTSR1-βarr2-ΔCT-Fab30L fusion construct, Fab30 heavy chain (Fab30H-His8), GRK5, and PIP5K were co-expressed in Spodoptera frugiperda (Sf9) insect cells using the Bac-to-Bac Baculovirus Expression System (Invitrogen)^45^. Sf9 cells, routinely tested for mycoplasma contamination, were cultured in SIM SF serum-free medium (Sino Biological) to a density of 3-4 × 10^6^ cells/mL. Cultures were then infected with the four baculovirus constructs (BRIL-NTSR1-βarr2-ΔCT-Fab30L, Fab30H-His8, GRK5, and PIP5K) at a ratio of 2:2:1:1, respectively, with a total multiplicity of infection (MOI) of ∼6. Cells were harvested by centrifugation 60 hours post-infection at 27 °C, and the pellets were stored at –80 °C.

### Purification of the NTSR1-Arrestin Complex

Cells expressing the NTSR1-arrestin complex were lysed in hypotonic buffer (10 mM HEPES pH 7.4, 10 mM MgCl_2_, 20 mM KCl) supplemented with EDTA-free protease inhibitor cocktail (TargetMol). After Dounce homogenization, lysates were centrifuged at 30,000 rpm and 4°C for 35 min. The membrane pellet was washed twice by Dounce homogenization in high-osmotic buffer (1.0 M NaCl, 10 mM HEPES pH 7.4, 10 mM MgCl_2_, 20 mM KCl, and EDTA-free protease inhibitor cocktail). Membrane solubilization was performed for 3-4 hours at 4°C in a buffer containing 20 mM HEPES pH 7.4, 200 mM NaCl, 25 mM imidazole, 10% glycerol, 1 µM PAMs, 1 µM NTS (8-13) peptide, 25 mM ATP, 0.5% DDM, 0.1% CHS, and protease inhibitors.

The solubilized membrane fraction was clarified by ultracentrifugation at 30,000 rpm and 4°C for 45 min. The supernatant was incubated with TALON^®^ Superflow Metal Affinity Resin (TaKaRa) for 2 hours at 4°C. The resin was washed sequentially with two buffers: first, 10 column volumes (CV) of washing buffer I (20 mM HEPES pH 7.4, 200 mM NaCl, 25 mM imidazole, 10% glycerol, 1 µM compound, 1 µM NTS (8-13), 0.075% LMNG, and 0.0015% CHS); and second, 5 CV of washing buffer II (same as buffer I, but with 50 mM imidazole, 0.0075% LMNG, and 0.00015% CHS). The complex was eluted with washing buffer II supplemented with 250 mM imidazole.

The eluted NTSR1-βarr2 complex was concentrated using a 100 kDa molecular weight cut-off (MWCO) Amicon Ultra centrifugal device. The concentrated complex was then subjected to size-exclusion chromatography (SEC) on a Superose^®^ 6 Increase 10/300 GL column (GE Healthcare) pre-equilibrated with 20 mM HEPES pH 7.4, 200 mM NaCl, 0.00075% LMNG, and 0.000015% CHS. Monomeric fractions of the NTSR1-βarr2 complex were collected and pooled.

### Cryo-EM Sample Preparation and Data Collection

For cryo-EM sample preparation, 3.5 µL of purified NTSR1–βarr2 complex was applied to glow-discharged Quantifoil holey carbon grids (ANTcryo R1.2/1.3, Au 300 mesh). Grids were blotted for 5 s with a blot force of 1 using a Vitrobot Mark IV (FEI) at 4°C and 100% humidity, then plunge-frozen in liquid ethane. Cryo-EM data were collected on a Titan Krios G4 microscope (300 kV, E-CFEG) at the Advanced Center for Electron Microscopy (Shanghai Institute of Materia Medica, Chinese Academy of Sciences) using EPU software. Images were recorded at a calibrated pixel size of 0.73 Å. The following numbers of movies were collected for the NTSR1-βarr2 complex bound to different compounds: 14,639 (SBI-553), 7,924 (N065), and 9,486 (N191).

### Image Processing

Motion correction and dose-weighting of raw movie stacks for the SBI-553-bound NTSR1-βarr2 complex (14,639 movies) were performed using MotionCor2^46^. Contrast transfer function (CTF) parameters were estimated with cryoSPARC’s Patch CTF^47^. Particle picking began with a 160 Å diameter blob picker, followed by template picking and Topaz training. Seven rounds of 2D classification selected 1,616,901 particles for 3D reconstruction. Multiple rounds of seed-facilitated multi-reference 3D classification and resolution gradient classification yielded 968,592 particles. To improve local resolution of the receptor region, local refinement focused on the receptor domain was carried out. The global consensus map and receptor-focused local-refined map were fitted and merged in ChimeraX, and the resultant composite map was used for subsequent atomic model building. Non-uniform and local refinement of these particles produced a final map at 2.88 Å resolution (Supplementary Fig. 3). Datasets for the N065– and N191-bound NTSR1-βarr2 complexes were processed similarly, yielding maps with global resolutions of 2.54 Å and 2.63 Å, respectively (Supplementary Figs. 4 and 5).

### Model Building and Refinement

Atomic modeling of the NTSR1–βarr2 complex was initiated using the previously determined NTSR1–βarr1 structure (PDB ID: 6PWC) as a reference^44^. The starting model was positioned within the corresponding cryo-EM density using Chimera, ISOLDE, and ChimeraX^48^ and subsequently adjusted and rebuilt in Coot (version 0.9.6)^49^. To facilitate interpretation of regions with enhanced local density, a composite map was generated by combining the consensus reconstruction with receptor-focused and receptor/βarr2-focused refinement maps, and was used for model building.

The resulting model was subjected to real-space refinement in PHENIX (version 1.20)^50^ and evaluated using comprehensive structural validation procedures. Stereochemical quality was assessed with MolProbity (version 4.5)^51^. Model-to-map agreement was evaluated using Fourier shell correlation (FSC), while potential overfitting was examined by comparing the refined model with the two independently generated half-maps. The final refinement statistics are reported in Table S5. Molecular graphics were generated using Chimera, ChimeraX, and PyMOL 2.5.2 (https://pymol.org/2/).

### Phospho-kinase array

HEK293T cells expressing NTSR1 were cultured in 10-cm dishes and treated with vehicle (HBSS supplemented with 20 mM HEPES and 0.01% BSA), NTS^8–13^ (1μM), SBI-553 (20 μM), or the combination of NTS^8–13^ and SBI-553 for 1 h at 37°C. After treatment, cells were lysed using the lysis buffer provided in the Proteome Profiler Human Phospho-Kinase Array Kit (ARY003C, R&D Systems). Total protein concentration was determined using a BCA Protein Assay Kit (Epizyme), and sample volumes were adjusted so that each membrane was incubated with 500 μg total protein. The phospho-kinase array was performed essentially according to the manufacturer’s instructions. This membrane-based antibody array uses paired Part A and Part B membranes. Briefly, all reagents were first equilibrated to room temperature, and the membranes were blocked in Array Buffer 1 for 1 h at room temperature. Cell lysates containing 500 μg total protein were then diluted to a final volume of 2 mL with Array Buffer 1 and incubated with paired Part A and Part B membranes overnight at 2–8°C on a rocking platform. After washing, Part A and Part B membranes were incubated with the corresponding biotinylated detection antibody cocktails for 2 h at room temperature. Following additional washes, membranes were incubated with Streptavidin-HRP for 30 min at room temperature. Signals were developed using equal volumes of the supplied chemiluminescent detection reagents, and subsequent steps were carried out according to the manufacturer’s protocol.

Membranes were imaged using an Amersham ImageQuant 800 imaging system (Cytiva). Spot intensities were quantified in ImageJ. Duplicate spots for each analyte were averaged, background was corrected by subtracting the signal from blank spots, and the resulting values were normalized to the reference spots for comparison of relative phosphorylation changes across treatment groups.

### Phosphoproteomic analysis

#### Sample preparation and phosphopeptide enrichment

HEK293T cells expressing NTSR1 were cultured in 10-cm dishes and treated with vehicle (HBSS supplemented with 20 mM HEPES and 0.01% BSA), NTS^8–13^ (1μM), each allosteric ligand alone (SBI-553, N065, or N191; 20 μM each), or in combination with NTS^8–13^, followed by incubation at 37°C for 1 h. Samples were lysed in protein lysis buffer containing 8 M urea, 1% SDS and protease inhibitors. Lysates were homogenized using a cryogenic grinder, sonicated at low temperature for 30 min, and centrifuged at 14,000 × g for 15 min at 8 °C. Supernatants were collected, and protein concentrations were determined by BCA assay.

For digestion, 30–50 μg protein from each sample was adjusted to 100 mM TEAB, reduced with 10 mM TCEP at 37 °C for 60 min, and alkylated with 40 mM iodoacetamide at room temperature for 40 min in the dark. Proteins were precipitated with acetone at −20 °C for 2 h, washed with 90% acetone, air-dried, and digested overnight at 37 °C with trypsin at an enzyme-to-protein ratio of 1:40 (w/w). Peptides were acidified, desalted, lyophilized and stored at −80 °C until phosphopeptide enrichment.

Phosphopeptides were enriched using enrichment beads. Peptides were resuspended in loading buffer containing 1 M glycolic acid, 80% acetonitrile and 5% trifluoroacetic acid, and incubated with equilibrated beads for 1 h. The beads were washed sequentially with loading buffer, 80% acetonitrile/1% trifluoroacetic acid, and 10% acetonitrile/0.2% trifluoroacetic acid. Phosphopeptides were eluted with 1% ammonium hydroxide, acidified to approximately pH 3 with 20% trifluoroacetic acid, and quantified using a NanoDrop One instrument.

#### LC–MS/MS acquisition

Equal amounts of enriched phosphopeptides were analyzed by DIA-based LC–MS/MS on a Vanquish Neo system coupled to a timsTOF HT mass spectrometer. Peptides were separated on a homemade analytical column (15 cm × 100 μm, 1.7 μm) using mobile phase A, 2% acetonitrile/0.1% formic acid, and mobile phase B, 80% acetonitrile/0.1% formic acid. A 14.4-min gradient was used. Data were acquired using Compass HyStar in positive-ion DIA mode with a capillary voltage of 1.6 kV, an ion mobility range of 1/K0 = 0.7–1.3 V·s cm⁻², and an MS scan range of 300–1500 m/z. LC–MS/MS analysis was performed by Majorbio Bio-Pharm Technology Co. Ltd. (Shanghai, China).

#### Data processing and quality control

Raw DIA data were processed using Spectronaut v19 against a human UniProt sequence database. Subsequent phosphosite-level statistical and bioinformatic analyses were performed using the Majorbio Cloud platform (www.majorbio.com). Protein– and peptide-level identifications were controlled at a false-discovery rate of 1%. Phosphorylation of serine, threonine and tyrosine was included in the post-translational-modification analysis, and localized phosphosites were quantified at the site level.

Quantitative phosphosite values were log2-transformed before downstream analysis. Measurements not passing the quantitative filtering criteria were treated as missing values. Sample reproducibility was assessed by pairwise Pearson correlation of phosphosite intensities and by within-group coefficient-of-variation analysis. Hierarchical clustering of the sample correlation matrix was used to evaluate within-group reproducibility and treatment-dependent separation.

#### Differential phosphorylation and phosphosite remodeling

Differential phosphorylation was assessed by pairwise comparison of biological replicates between the indicated treatment conditions. The primary contrasts used to evaluate intracellular allosteric modulation were NTS + SBI-553 versus NTS, NTS + N065 versus NTS, and NTS + N191 versus NTS. Comparisons among the three NTS + modulator conditions were additionally performed to identify modulator-specific phosphosite responses.

For visualization of global phosphorylation patterns, phosphosites were ranked according to their variation across treatment conditions, and the most variable sites were displayed after row-wise scaling of log2-transformed intensities.

To quantify phosphosite remodeling under a common orthosteric background, the effect of each intracellular modulator was defined as

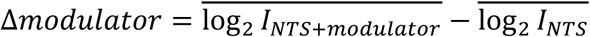

where *I* denotes phosphosite abundance. Phosphosites showing the largest absolute remodeling effects across SBI-553, N065 and N191 were selected for visualization.

To assess departures from additivity, a descriptive interaction term was calculated on the log2 scale as

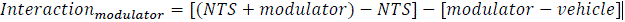

The interaction term was used as a descriptive effect-size measure and was not interpreted independently as statistical evidence of receptor allostery.

Differential phosphosite sets obtained from NTS + SBI-553 versus NTS, NTS + N065 versus NTS and NTS + N191 versus NTS were compared to identify shared and modulator-specific phosphorylation responses. Direct comparisons of NTS + N065 and NTS + N191 with NTS + SBI-553 were used to identify phosphosites that differed among modulators under the same NTS background.

#### Kinase-substrate enrichment analysis

Kinase-substrate enrichment analysis (KSEA) was performed at the phosphosite level to infer changes in kinase-associated signaling from coordinated phosphorylation changes among annotated kinase substrates. For each treatment comparison, quantitative changes in phosphosites were mapped to curated kinase–substrate relationships, and an enrichment z-score was calculated for each kinase based on the collective phosphorylation changes of its matched substrate sites. Only kinases represented by at least five quantified substrate phosphosites were included in the analysis. Kinase sets with P < 0.05 were retained for visualization. KSEA z-scores were compared across treatment contrasts to identify kinase-associated phosphorylation programs differentially affected by the intracellular modulators. Because KSEA infers kinase-associated activity from substrate phosphorylation patterns, these scores were not interpreted as direct measurements of kinase recruitment or receptor–kinase association.

#### Pathway enrichment analysis

Proteins associated with differential phosphosites were subjected to KEGG pathway enrichment analysis. The principal analyses compared NTS + SBI-553, NTS + N065 and NTS + N191 individually with NTS alone, thereby assessing each intracellular modulator against a common orthosteric background. Adjusted P values were used to assess pathway-level significance. Enrichment results were visualized according to pathway rich factor, the number of mapped differential proteins and adjusted P value. For descriptive comparison of pathway repertoires, pathways identified after NTS stimulation were used as the reference set. Pathways detected after both NTS and NTS + modulator treatment were classified as retained, pathways detected only after co-treatment as gained, and pathways detected after NTS alone but not after co-treatment as lost. This classification was used descriptively and was not considered a formal statistical test of allosteric interaction.

### Animals

C57BL/6 mice (8-16 weeks) were purchased from Shanghai Lingchang BioTech Co., Ltd. (Shanghai, China). Body temperature, wire hang, pole descent, blood pressure, contact righting studies and monitoring of metabolism employed male mice. Morris water maze, Y maze, Novel object recognition, forced swim test, Elevated-O-maze, Novelty-suppressed feeding test, food intake utilized male C57BL/6J mice. In all studies, littermates of the same sex were randomly assigned to experimental groups. Mice were maintained on a 12:12 hour light: dark cycle. Experiments were conducted at the beginning of the light cycle. Tap water and standard laboratory chow were supplied *ad libitum*, except during testing. DIO C57BL/6J mice (male, 13 weeks on high fat diet since week 5) were purchased from Beijing GemPharmatech Co., Ltd. Animals were acclimatized for 1 week prior to the experiment. Mice were housed 5/cage and 12-hour lighting conditions (8:00 am to 8:00 pm) in a controlled environment (18∼26 °C). DIO animals had free access to purified water and high fat diet (Research Diets Cat: D12492). All animal studies and experimental procedures were approved by the Animal Care and Use Committees at Shanghai Jiao Tong University School of Medicine (policy number: DLAS-MP-ANIM, 01-05).

### *In vivo* drug treatment

SBI-553 HCL (SBI-553) (12 mg/kg) and N065 HCL (N065) (12 or 30 mg/kg) were solubilized in 5% DMSO, 40% PEG300, 5% Tween-80, and 50% saline to a stock concentration of 1.2 mg/mL and injected intraperitoneally (i.p.) according to the experimental schedule at a volume of 5 mL/kg or 10 mL/kg body weight. Vehicle (5% DMSO + 40% PEG300 + 5% Tween 80 + 50% saline) was administered to control mice. Method-specific compounds are described with their respective procedures. Behavioral testing was conducted at the times of peak drug effect: 30 min post-treatment for the SBI-553 group and N065 group.

Lipopolysaccharide (LPS) (25 mg; Sigma L2880 Escherichia coli 055: B5) was prepared in 5 mL of sterile pyrogen-free 0.9% saline, aliquoted and stored at 20 ℃. On the day of study, 600 μL of LPS was thawed and diluted in 400 μL of sterile saline. Each mouse received a dose equivalent to 15 mg/kg of body weight i.p at a volume of 5 mL/kg. Control animals received 0.9% sterile saline. LPS administration mimics the endotoxaemia seen in meningococcaemia, bacteraemia antibacterial therapy, and the early cytokine storm and blood pressure drop seen during vascular shock^37^.

### Pharmacokinetics of N065 and SBI-553

ICR male mice weighing 35-41 g (n = 3 per time point) were used for the pharmacokinetic study. Both N065 and SBI-553 were administered using the same regimens: intravenously (i.v.) at 1 mg/kg and intraperitoneally (i.p.) at 12 mg/kg, with an administration volume of 10 mL/kg. The i.v. formulation was 5% DMSO + 5% Solutol + 90% saline, and the i.p. formulation was 5% DMSO + 40% PEG300 + 5% Tween-80 + 50% saline. At designated time points (0.083, 0.25, 0.5, 1, 2, 4, 8, and 24 h post-dose), blood samples (approximately 50 µL) were collected from the retro-orbital venous plexus into heparin sodium tubes and kept on wet ice. Plasma was separated by centrifugation at 8000 rpm for 5 min at 4 °C and stored at –80 °C until analysis. Whole brain tissues were collected only from the i.p. administration groups at 0.5 h and 1 h post-dose, rinsed with phosphate-buffered saline, and flash-frozen. Brain samples were stored at –80 °C until analysis.

Before LC-MS/MS analysis, brain samples were homogenized with two volumes of water (brain: water = 1:2), and brain homogenate concentrations were multiplied by the dilution factor of 3. A 10-µL aliquot of plasma or brain homogenate was mixed with 300 µL acetonitrile containing an internal standard (IS), vortexed for 1 min, and centrifuged at 12,000 rpm for 5 min. Supernatant was token for injection. Calibration standards and QC samples were prepared in blank ICR mouse plasma or blank ICR mouse brain homogenate. The calibration range was 1-5000 ng/mL, with an LLOQ of 1 ng/mL, and linear regression with 1/x² weighting was used. LC-MS/MS analysis was performed on a Shimadzu LC30AD HPLC system coupled to a SCIEX Triple Quad 5500 mass spectrometer with ESI+. MRM transitions were: N065, m/z 504.7 → 439 (CE 65, DP 100); SBI-553, m/z 450.8 → 405.2 (CE 55, DP 100); IS, m/z 393.3 → 147.1 (CE 30, DP 100).

### Behavior testing

All mice were tested by a researcher in a blinded manner starting at eight weeks of age. Mice were allowed to settle for at least 2 days after cage changing before they were tested. Tests were performed 1 day apart to enable the mice to rest between tests. The mice were acclimatized to the behavior-testing room for 1 h before testing. Mice were tested during the light phase of the light cycle.

#### Pole descent test

To evaluate drug-induced alterations in gross motor function, pole descent was conducted as described previously^52^. A 0.5 m long pole, 1 cm in diameter, was wrapped with non-adhesive shelf liner to improve the grip of the mice, which were placed into the home cage. Mice were acclimated over a two-day training period to descend from the pole into their cage. On the first training day, each mouse performed three trials: initially placed head-down at one-third of the pole’s height, then at two-thirds, and finally at the top. On the second day, all mice completed three consecutive trials descending head-first from the top of the pole. On the second day of training, the mice were given three trials to descend, head-down, from the top of the pole. On the third day, animals received either N065 (12 mg/kg), SBI-553 (12 mg/kg), or vehicle (5% DMSO, 40% PEG300, 5% Tween-80 in saline). Behavioral testing was conducted at 30 min post-treatment. The animals were placed head-down on the top of the pole and timed to descend back into the home cage. Timing began when the experimenter released the mouse and ended when one hindlimb reached the home cage base.

#### Wire hang test

To assess drug-induced myorelaxant effects and/or coordination changes, the horizontal wire test was conducted as previously described^52^. On the experimental day, animals were administered N065 (12 mg/kg), SBI-553 (12 mg/kg), or vehicle. Behavioral assessment was performed 30 min post-treatment. The mice were positioned at the center of a square wire-mesh screen (30 × 30 cm, 1 cm grid). The screen was then rotated head-over-tail and placed on supports approximately 40 cm above an open cage containing clean bedding. The time until the mouse released its grip was recorded, with a maximum cutoff of 60 s.

#### Contact righting test

To detect drug-induced sedation, changes in the righting reflex were assessed. On the day of the experiment, mice were treated with N065 (12 mg/kg), SBI-553 (12 mg/kg), or vehicle. Behavioral testing was performed 30 min after administration. Each mouse was placed upright inside a clear glass tube (6.5 cm diameter × 43 cm length), and the tube was swiftly rotated 180° for six consecutive trials. The latency to regain an upright posture was recorded for each trial. Responses were scored according to the following scale: −6 for failure to right, –5 for >5 s delay in righting, –4 for >2 s delay in righting, –3 for >1 s delay in righting, –2 for righting from back <1 s, –1 for immediately rolling to the side and righting within 1 s, and 0 for no delay in righting.

#### Monitoring of metabolism

To assess drug-induced alterations in metabolism, mice were monitored using the Columbus Instruments Comprehensive Lab Animal Monitoring System (CLAMS). Prior to monitoring, animals were administered either N065 (12 mg/kg), SBI-553 (12 mg/kg), or vehicle once daily for seven consecutive days. Following treatment, mice were individually housed in metabolic cages for a 5-day period under a controlled 12-hour light/dark cycle (lights on from 7:00 to 19:00) with ad libitum access to food and water. Using CLAMS, indirect calorimetry was performed to measure the respiratory exchange ratio (RER), oxygen consumption rate (VO₂), and carbon dioxide production rate (VCO₂). Data acquired from CLAMS were analyzed with CalR software. Data are presented for the last 24 h or 48 h of the 5-day period.

#### Blood pressure measurement

The effects of SBI-553 and N065 on blood pressure were assessed in mice using the CODA non-invasive blood pressure system (Kent Scientific, Torrington, CT, USA). This system employs volume-pressure recording (VPR) technology to detect tail-volume changes corresponding to systolic and diastolic BP, utilizing two tail cuffs: an occlusion cuff (O-cuff) and a volume-pressure recording cuff (VPR-cuff). On the test day, animals were administered N065 (12 mg/kg), SBI-553 (12 mg/kg), or vehicle. Measurements were performed 30 min after treatment. Each mouse was placed in an adjustable restrainer to minimize movement during recording and allowed to acclimate for at least 10 min before starting. Following acquisition of 10 consecutive measurements, the animal was promptly removed from the restrainer. For each mouse, the averages of systolic blood pressure, diastolic blood pressure, heart rate, flow velocity, and volume were calculated individually.

#### Locomotor activity

Locomotor activity alterations induced by drugs were evaluated using the open field test following an established protocol. The assay was conducted in 40 × 40 cm² white Plexiglas arenas. An overhead camera recorded the sessions, and the EthoVision XT 4 software package (Noldus Information Technology) was employed for tracking and analysis. In the experiment, male mice were first placed in the arena for a 30-minute acclimation period, followed by intraperitoneal injection of vehicle, N065 (12 mg/kg), or SBI-553 (12 mg/kg). Each mouse was immediately returned to the arena after injection, and locomotor activity was monitored for a total duration of 1.5 hours, which included the 30-minute pre-injection acclimation. The total distance travelled were analyzed.

#### Elevated-O-maze test

To evaluate drug-induced alterations in anxiety-like behavior, the elevated-O-maze test was conducted according to a published method^53^. The circular maze (radius = 55 cm), elevated 60 cm above the floor, comprised two open and two enclosed arms of identical dimensions. The enclosed arms were equipped with 15-cm-high walls. On the test day, mice were administered vehicle, N065 (12 mg/kg), or SBI-553 (12 mg/kg). Behavioral assessment began 30 min post-injection. Each mouse was placed in the maze under dim lighting for a 5-min session, during which the time spent in the open arms was recorded.

#### Core body temperature measurement

Core body temperature measurements were obtained from male and female C57BL/6J mice using a rectal probe thermometer (BAT-12 Microprobe Thermometer, Physitemp Instruments, Clifton, NJ). To habituate the animals to the procedure, they were handled and their temperatures were taken during three separate sessions in the week preceding the experiment.

Single-compound dosing. Following baseline temperature recording, mice received an intraperitoneal (i.p.) injection of N065 (12 mg/kg), SBI-553 (12 mg/kg), or vehicle at a volume of 10 ml/kg. Temperatures were subsequently measured at 30, 60, 120, and 300 min post-injection.

Multiple-compound dosing. Following baseline temperature recording, mice were first pretreated with an i.p. injection of N065 (12 mg/kg), SBI-553 (12 mg/kg), or vehicle at 5 mL/kg. 45 minutes later, core body temperature was recorded (designated as Time 0). Immediately following this measurement, mice were administered a second i.p. injection of either PD149163 (2 mg/kg) or its vehicle (0.2% DMSO in saline), also at 5 mL/kg. Core body temperature was then monitored at 60, 120, 180, 240, 300, 360, 420, and 480 min following the PD149163/vehicle administration.

#### Home cage food consumption test

The effects of N065 and SBI-553 on food intake were evaluated under fed, short-term fasted (6 h), and long-term fasted (17 h) states. Mice were administered N065 (12 mg/kg), SBI-553 (12 mg/kg), or vehicle 30 minutes before the feeding test. In fasting studies, drug administration occurred at 5.5 h (short-term) or 16.5 h (long-term) into the fast. Food intake was measured for 1 hour in short-term fasted mice, and for 6 hours in fed and long-term fasted mice.

#### Novel objective recognition test

The novel object recognition test was conducted to evaluate the effects of N065 and SBI-553 on recognition memory. Mice were first habituated to the empty testing arena (40 × 40 × 40 cm) for 5 min and then rested in the experimental room for 24 h prior to the formal test. On the first day of testing, animals were treated with N065 (12 mg/kg), SBI-553 (12 mg/kg), or vehicle. Behavioral testing was conducted at 30 min post-treatment. Objects A and B (with different shapes and colors) were positioned in opposite corners of the arena. Each mouse was introduced from the center and allowed to freely explore both objects for 5 min. After a 24-hour interval, on the test day (Day 2), mice received the same drug treatment. 30 min later, they were returned to the arena, in which one familiar object (B) had been replaced with a novel object (C). The exploration time directed toward objects A, B, and C was recorded. The recognition index was calculated as TA / (TA + TC) and TC / (TA + TC) on the second day. TA and TC represent the time the mice explored objects A (old) and C (new), respectively.

#### Morris water maze Test

The Morris water maze (MWM) test was employed to assess the impact of N065 and SBI-553 on spatial memory. The apparatus (Techman Software Co., Ltd., Chengdu, China) included a circular pool (diameter: 1.2 m; height: 50 cm) filled with water maintained at 22 ± 1 °C and rendered opaque with milk powder. A 10-cm diameter escape platform was submerged 0.5 cm below the water surface in a constant position within the target quadrant. Distinct visual cues were positioned around the pool, and an overhead camera recorded all sessions. Prior to testing, mice were acclimated to the behavior room. The acquisition phase consisted of 5 consecutive days, with each animal completing four 60-second trials per day. If a mouse located the hidden platform within 60 s, it remained there for 20 s; if not, it was gently guided to the platform and allowed to stay for the same duration. Escape latency (time to find the platform) was recorded for each trial. 24 hours after the final training session, a probe trial was conducted with the platform removed. Each mouse was allowed to swim freely for 60 s and time in the target quadrant was recorded.

#### Y maze test

Y maze test was conducted to evaluate the effect of N065 and SBI-553 on working memory. The apparatus consisted of three enclosed arms (each 28 cm long, 6 cm wide, and 18 cm high) arranged at 120° angles from a central area, labeled A, B, and C. Mice were placed in the center and allowed to freely explore the maze for 8 minutes. An entry was recorded when all four paws passed into an arm. Spontaneous alternation was defined as the continuous entry of mice into three arms, such as ABC, BCA, or CAB. The percentage of spontaneous alternation rate was calculated: % spontaneous alternation rate = number of spontaneous alternations/ (total number of arm entries − 2) × 100.

#### Echo magnetic resonance imaging

The EchoMRI-1100 system (EchoMRI LLC, Houston, TX, USA) was used to evaluate the effects of N065 and SBI-553 on body composition in DIO mice. Each mouse was placed in an adjustable plastic cylinder to restrict movement. The cylinder (1.25 inches in diameter) had holes on each end to allow for the animal to breathe normally. The cylinder containing the mouse was then inserted into the instrument, and a scan lasting 30-75 seconds was performed. The obtained readings included lean mass, fat mass, total body water, and free body water.

#### Forced swim test

The effects of N065 and SBI-553 on depression-related behaviors were evaluated using the forced swim test. Briefly, a cylindrical tank (20 cm in diameter, 30 cm in height) was filled with water to a depth of 15 cm and maintained at 25 ± 1 °C. Each mouse was placed individually into the water for a 6-minute session. After an initial 2-minute adaptation period, immobility time was recorded over the remaining 4 minutes. Immobility was defined as passive floating with only slight movements necessary to keep the head above water.

### Statistical analysis

All data are expressed as mean ± standard error of the mean (SEM) unless otherwise described. Statistical analyses were performed using GraphPad Prism and the significance of differences was assessed by two tail unpaired Student’s t test or one-way or two-way analysis of variance (ANOVA) followed by Tukey’s multiple comparisons test. Parametric statistics were used for data that met normal (gaussian) distribution assumptions, as determined by a Shapiro-Wilk test. Wire hang and contact righting data, which were not normally distributed, were analyzed using nonparametric statistics (i.e., one-way ANOVA on ranks). The statistical parameters can be found in the figures and figure legends. Significance was set as *p* < 0.05 and expressed as *\*p* < 0.05, *\*\*p* < 0.01, and *\*\*\*p* < 0.001. The number of biological replicates is indicated in the figures. For phosphoproteomic analysis, all data were uploaded to the Majorbio Cloud Platform (cloud.majorbio.com) for analysis.

### Figure illustrations

Schematics were created using BioRender. Protein structure images were generated using ChimeraX (UCSF, v1.6) or PyMOL. Chemical structures were drawn using ChemDraw.

## Data availability

The density maps and structure coordinates have been deposited to the Electron Microscopy Data Bank (EMDB) and the Protein Data Bank (PDB) with accession numbers EMD-67631 and 21FR, respectively, for the NTSR1-β-arrestin2 complex bound to SBI-553; EMD-67595 and 21DN, respectively, for the NTSR1-β-arrestin2 complex bound to N065; and EMD-67592 and 21DM, respectively, for the NTSR1-β-arrestin2 complex bound to N191. The mass spectrometry proteomics data have been deposited to the ProteomeXchange Consortium with the dataset identifier PXD078536.

## Funding

Innovative Drug Research and Development National Science and Technology Major Project (2025ZD1803100 & 2025ZD1803104 to J.Z.)

Shanghai Municipal Health Commission (2025ZHYL038 to J.Z.)

Lingang Laboratory (LG8888 to J.Z.)

National Natural Science Foundation of China (82441035, 22237005, 824B2105, 82504673 to J.Z.; 32130022, 82495184, 82121005 to H.E.X.; 32401002 to Y.Z.)

National Key R&D Program of China (2023YFC3605500 to W.Y.; 2022YFC2703105, 2022YFE0203600 to H.E.X.; 2024YFA1307504 to Y.Z.)

Zhongshan Municipal Bureau of Science and Technology (CXTD2023010 to W.Y.) CAS Strategic Priority Research Program (XDB37030103 to H.E.X.)

Shanghai Municipal Science and Technology Major Project (2019SHZDZX02 to H.E.X.)

The HE Research Fellowship (HERF2025011 to Y.Z.)

Shanghai Leading Talent Program of Eastern Talent Plan (QNKJ2025058 to Y.Z.)

The Innovative Research Team of High-Level Local Universities in Shanghai (Y.Z.).

## Author Contributions

J.Z. conceived and supervised the project. J.Z. and W.Y. designed the experiments. X.S., C.Z. and X. Lan performed the BRET assays, Tango assays, site-directed mutagenesis and related analyses under the supervision of J.Z. M.Z. designed and synthesized the small molecules. P.Y. purified the protein complexes. Q.Y., Z.Z. and P.Y. performed cryo-EM data analysis, structural refinement and preparation of structural figures under the supervision of W.Y. R.Z., X.S., X. Lu, X.Q., R.L. and M.L. performed structural analysis, data analysis and bioinformatics analysis. K.C. and Y.Z. performed animal experiments and data analysis under the supervision of J.Z. and Y.Z. X.S., R.Z., K.C. and M.Z. wrote the manuscript, and all authors contributed to specific sections of the manuscript. J.Z., W.Y., H.E.X., Y.Z. and W.X. discussed the results and revised the manuscript.

## Declaration of generative AI and AI-assisted technologies in the writing process

During the preparation of this work, the authors used ChatGPT by OpenAI to improve language. After using this tool, the authors reviewed and edited the content as needed and take full responsibility for the content of the publication.

## Extended data

**Extended data Fig. 1.**
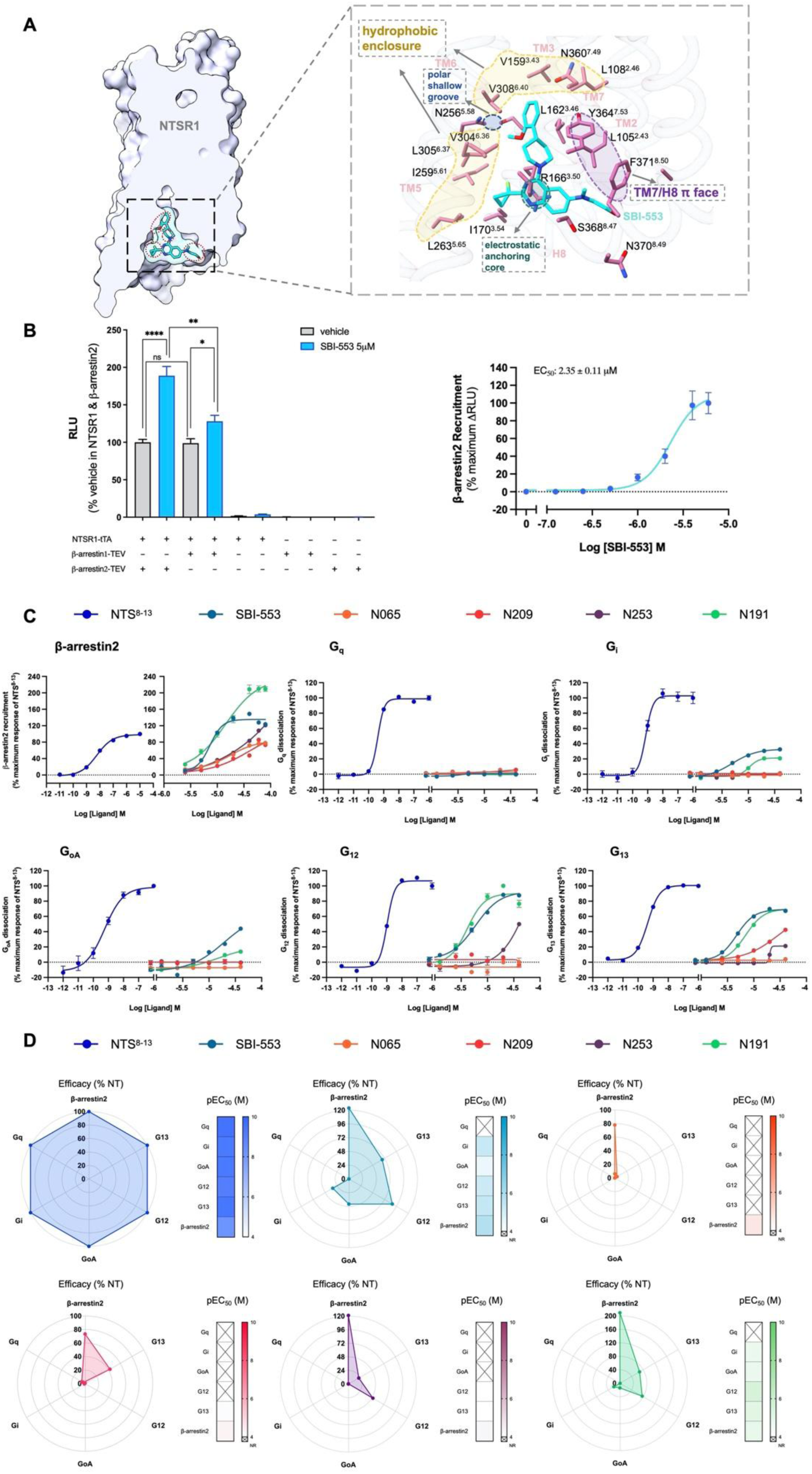
| Design, screening and profiling of SBI-553 analogs. (**A**) Surface and close-up views of SBI-553 bound to the NTSR1 intracellular allosteric pocket. Key residues shaping four interaction microdomains are annotated. **(B)** β-arrestin2 recruitment assay for SBI-553. Left, reporter output (RLU) comparing vehicle and SBI-553 (5 μM) under indicated component combinations. Right, concentration–response curve for SBI-553. Data are presented as mean ± s.e.m. from three independent experiments. **(C)** Intrinsic agonism profiling across β-arrestin2 and G-protein pathways (G_q_, G_i_, G_oA_, G_12_, G_13_) for NTS^8–13^, SBI-553 and representative analogs. Responses are shown as concentration–response curves (four-parameter sigmoidal curves) normalized to the maximal response of NTS^8–13^ within each pathway. Data are presented as mean ± s.e.m. from three independent experiments. **(D)** Radar summaries of efficacy (% of NTS^8–13^) and pEC_50_ across pathways for each ligand.

**Extended data Fig. 2.**
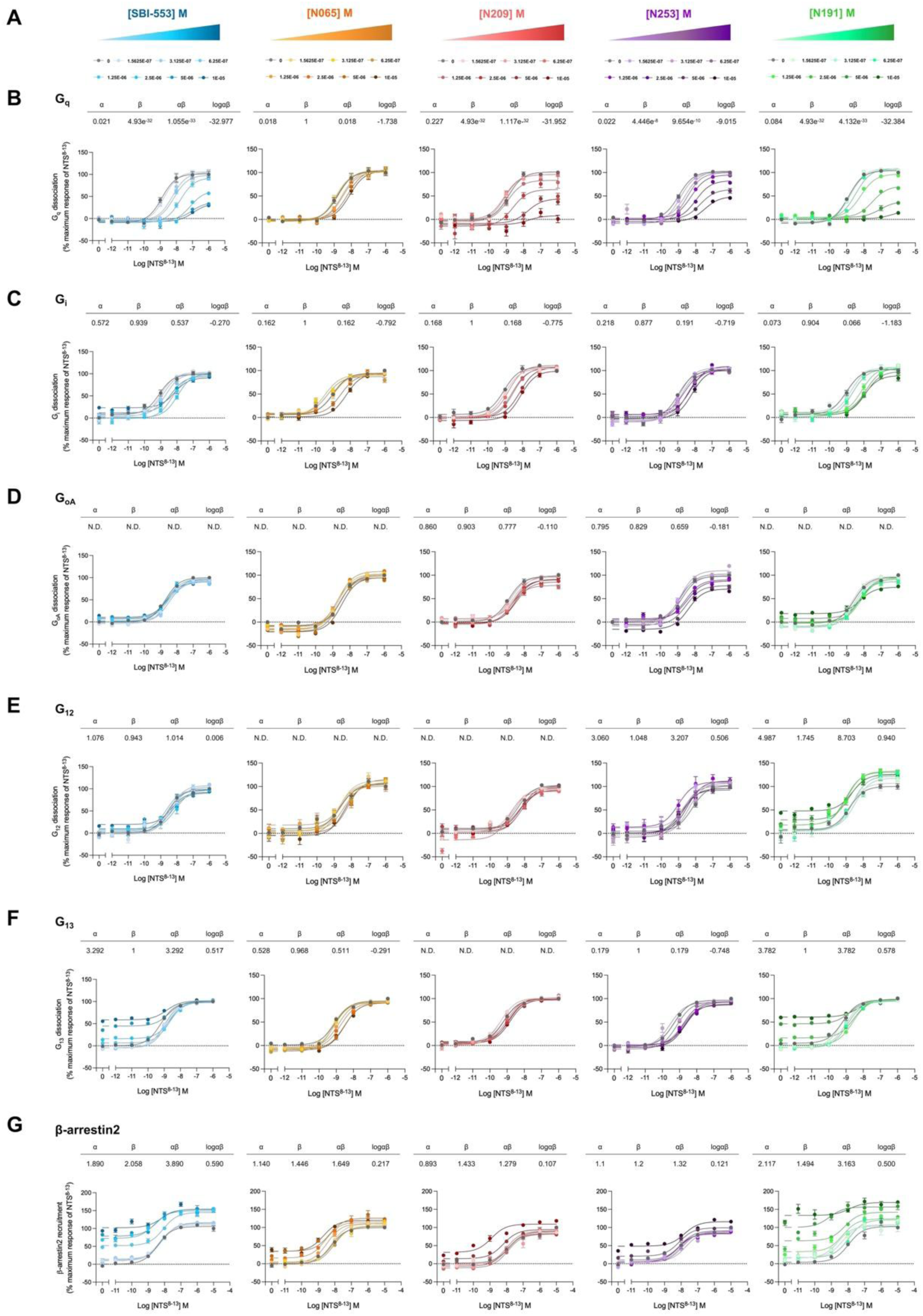
| Quantization of allosteric modulation by fitting the allosteric model across pathways. **(A)** Concentration scheme for 5 allosteric ligands. **(B–G)** Allosteric operational model fits to NTS^8–13^ concentration–response curves in the presence of increasing concentrations of intracellular ligands across pathways: (B) G_q_, (C) G_i_, (D) G_oA_, (E) G_12_, (F) G_13_, (G) β-arrestin2. ND indicates parameters not determined due to insufficient measurable modulation/response within the fitted conditions. Data are normalized to the maximal response of NTS^8–13^ in each pathway and presented as mean ± s.e.m. from three independent experiments.

**Extended data Fig. 3.**
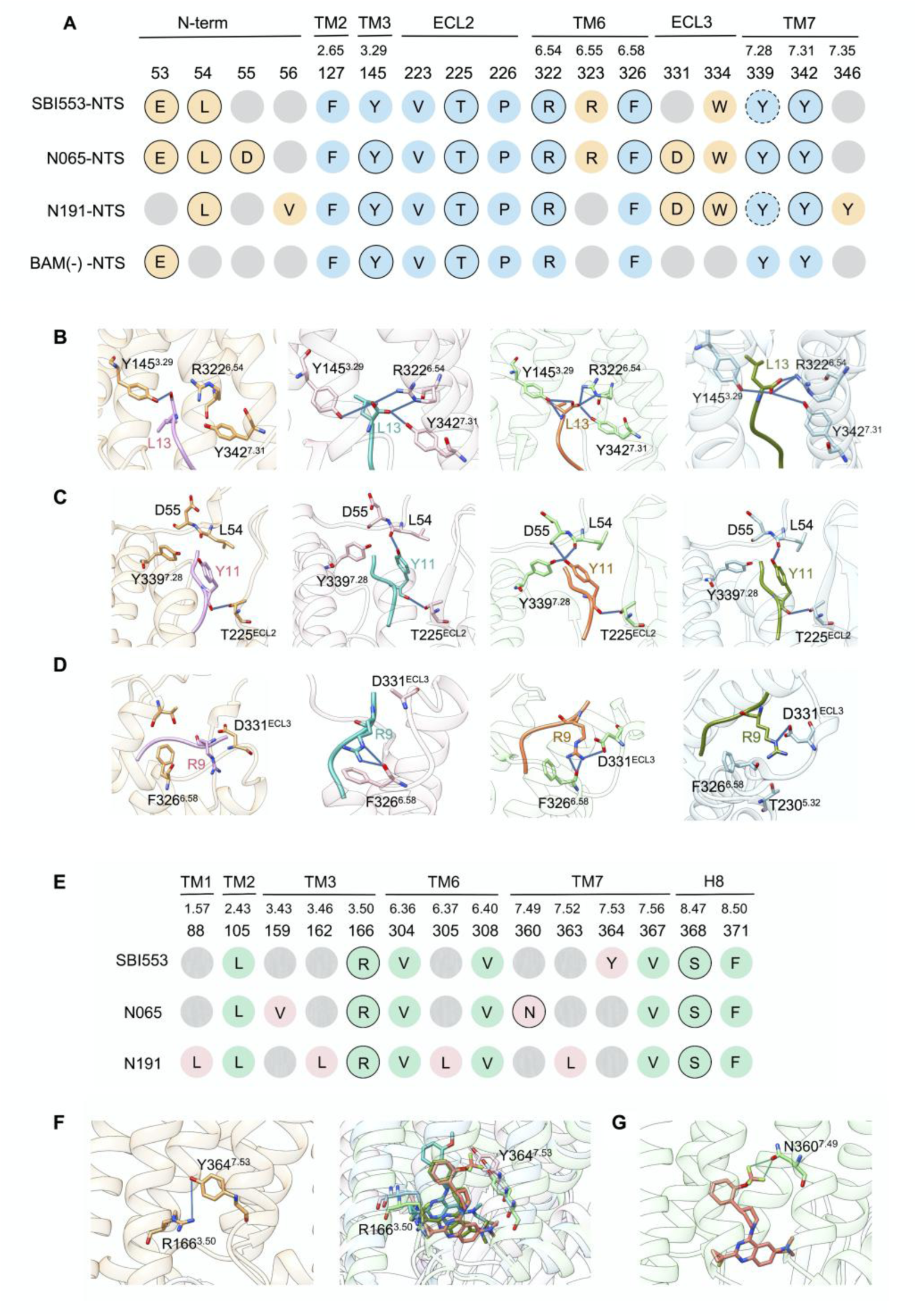
| Ligand-specific interaction fingerprints of NTS^8–13^ and allosteric modulators in NTSR1. **(A)** Interaction fingerprint map of NTS^8–13^ interactions in NTSR1 structures bound to SBI-553, N065, or N191, compared with the BAM-absent control structure (PDB: 6UP7). Interactions conserved across all four structures are shown in blue, whereas interactions unique to individual structures are shown in orange. Gray circles indicate the absence of contact. Solid outlines denote hydrogen-bond interactions, dashed outlines denote π-cation interactions, and circles without outlines denote hydrophobic interactions. **(B–D)** Comparison of key NTS-receptor interactions across the four NTSR1 structures. From left to right, close-up views show specific NTS–NTSR1 interactions in the BAM-absent, SBI-553-bound, N065-bound, and N191-bound structures, respectively. **(E)** Interaction fingerprint map of the intracellular allosteric pocket for SBI-553, N065, and N191. **(F)** Structural views illustrating a conserved polar anchoring interaction centered on R166^3.50^ and the adjacent Y364^7.53^ region within the intracellular pocket across the three modulators. **(G)** Close-up of N065 showing a ligand-specific short contact between its fluorine substituent and N360^7.49^, consistent with a halogen-bonding geometry. In panels (B–D), structures are arranged from left to right as BAM (–), SBI-553, N065, and N191; panels (F–G) follow the same color scheme. The orthosteric peptide and allosteric ligands are shown as sticks.

**Extended data Fig. 4.**
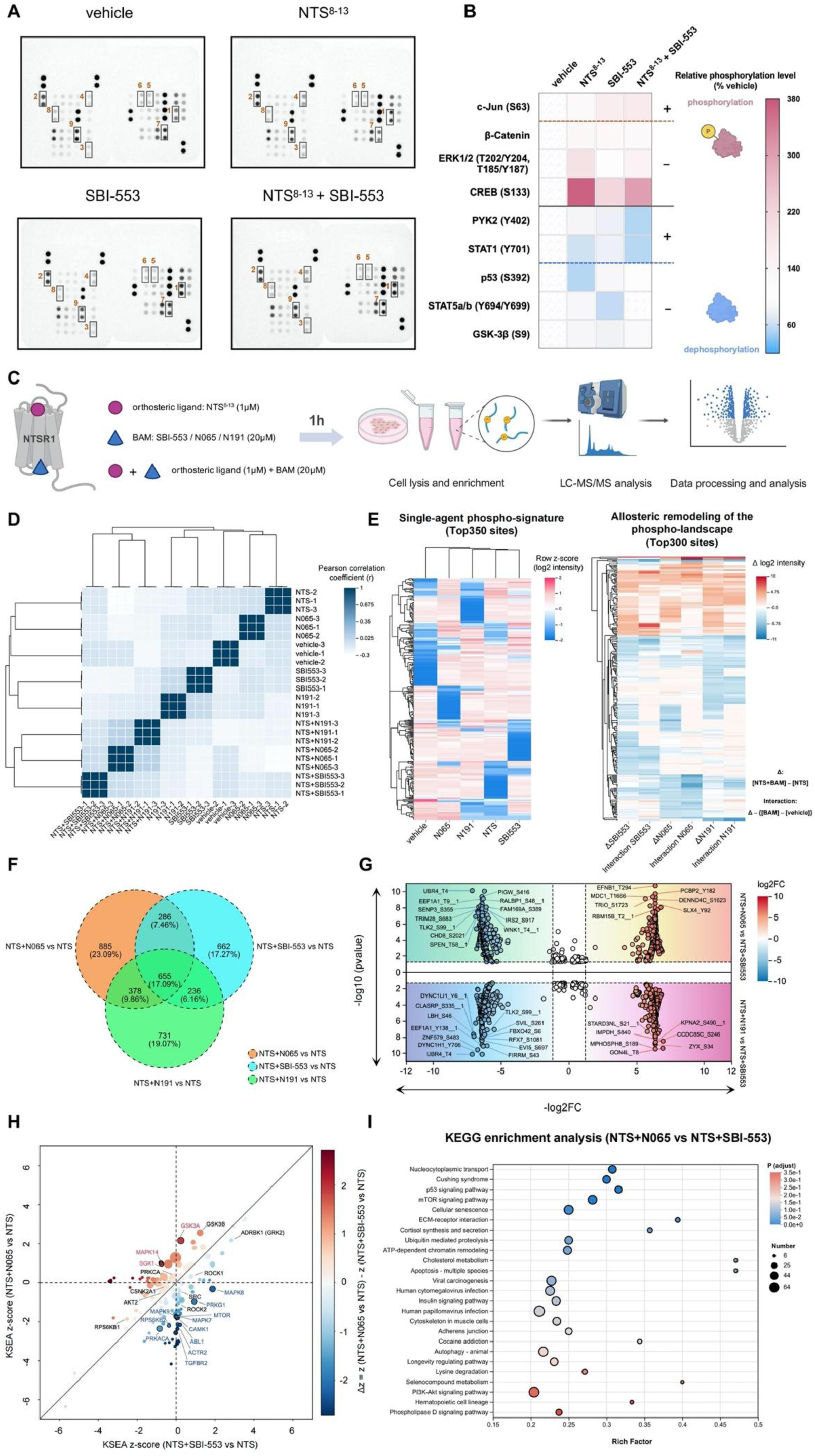
| Allosteric ligands shape distinct phospho-landscapes and drive allosteric remodeling. **(A)** Representative phospho-kinase array membranes for vehicle, NTS^8–13^, SBI-553 and NTS^8–13^ + SBI-553. **(B)** Semi-quantitative heatmap of selected nodes (relative to vehicle). Phosphorylation levels were quantified using Image J, normalized relative to the vehicle. Red represents phosphorylation, and blue represents dephosphorylation. “+” indicates an enhancement of NTS^8–13^ activity due to the introduction of SBI-553, and “-” indicates a weakening of NTS^8–13^ activity due to the introduction of SBI-553. **(C)** Phosphoproteomics workflow. **(D)** Sample–sample Pearson correlation matrix with hierarchical clustering, assessing within-group reproducibility and between-group separation. **(E)** Left, single-agent phospho-signatures (Top350 sites; row-scaled log2 intensities). Right, remodeling heatmap under the NTS background (Top300 sites; Δlog2 intensity) with definitions: Δ= [*NTS* + *BAM*] − [*NTS*]; *Interaction* = Δ − ([*BAM*] − [*ve*ℎ*icle*]), allowing separation of additive ligand effects from non-additive allosteric effects. **(F)** Venn diagram of differential phosphosites for NTS+BAM versus NTS, highlighting a shared core and ligand-specific branches. Differential sites were defined as |log_2_FC| > 1.2 and *p* < 0.05. **(G)** SBI-553-referenced volcano plots showing differential phosphosites between allosteric conditions under an NTS background, with representative sites annotated. The x-axis shows log_2_FC and the y-axis shows –log_10_(*p* value). Significant phosphosites were defined as |log_2_FC| > 1.2 and *p* < 0.05. **(H)** KSEA rewiring map: x-axis, KSEA z-score (NTS+SBI-553 vs NTS); y-axis, KSEA z-score (NTS+N065 vs NTS); color indicates Δ z-score. Only kinase sets with at least five matched substrate sites were included (m ≥5 and *p* < 0.05). Representative kinases are annotated to illustrate differential kinase-program remodeling across allosteric ligands. KEGG enrichment bubble plot showing significantly enriched pathways from the differential phosphoproteins in NTS+N065 vs NTS+SBI-553. The x-axis indicates rich factor, bubble size indicates the number of hits, and color indicates adjusted significance (*p_adj_*).

