## Supplementary Information for "Fine-tuning an intracellular allosteric pocket programs GPCR signaling bias"

33

### 34 **Chemistry**

35 1. Synthetic routes of final compounds N065, N191, N209 and N253 .....4

36 2. NMR, HRMS and HPLC spectra of final compounds N065, N191, N209 and N253 ..... 17

37

### 38 **Supplementary tables**

39 1. Table S1. Curve parameters and statistical comparisons presented in Figure 4 and Figure S23.26

40 2. Table S2. Curve parameters and statistical comparisons presented in Figure 4 and Figure S23.28

57 8. Fig. S8 | Surface expression level and NTS<sup>8-13</sup> agonist activity of NTSR1 mutants across

79

80

81

82

83

84

### Chemistry

#### Synthetic routes of final compounds N065, N191, N209 and N253

##### Scheme 1. Synthesis of final compound N065 <sup>a</sup>

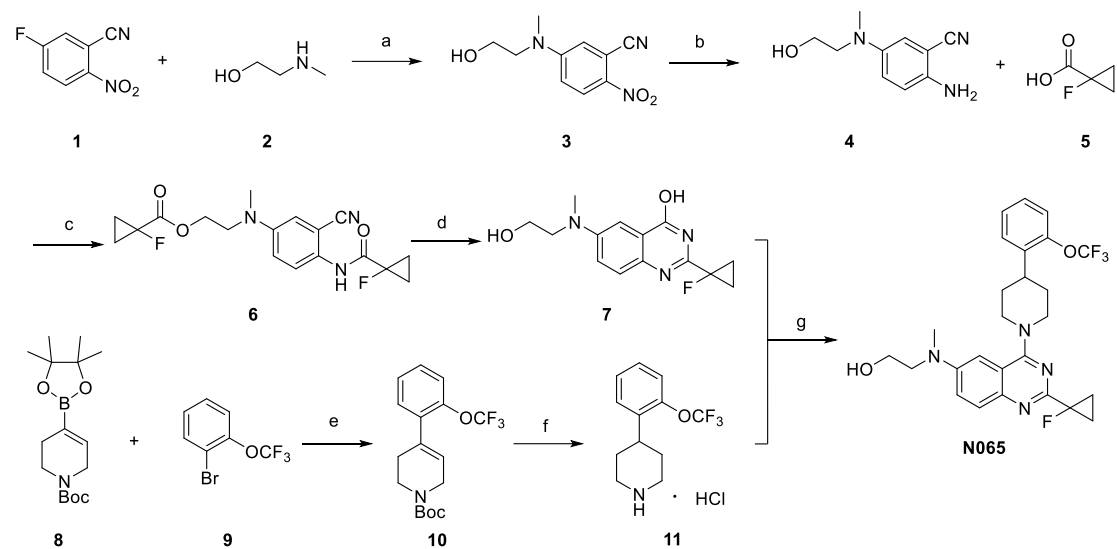

<sup>a</sup> Reagents and conditions: (a) TEA, IPA, rt; (b) Fe, NH<sub>4</sub>Cl, EtOH, MeOH, H<sub>2</sub>O, 80 °C; (c) TCFH, NMI, MeCN, rt; (d) NaOH, H<sub>2</sub>O<sub>2</sub>, EtOH, 85 °C; (e) Pd(dppf)Cl<sub>2</sub>, K<sub>2</sub>CO<sub>3</sub>, Dioxane, H<sub>2</sub>O, 85 °C; (f) H<sub>2</sub>, Pd/C, HCl/MeOH, rt; (g) BOP, DBU, MeCN, rt.

##### Synthesis of 5-((2-hydroxyethyl)(methyl)amino)-2-nitrobenzonitrile (intermediate 3)

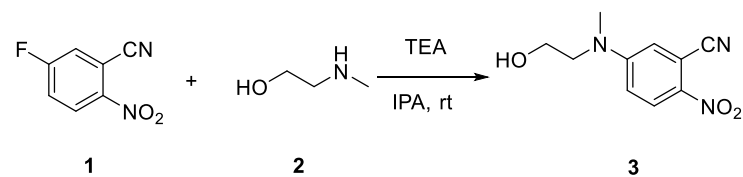

To a solution of 5-fluoro-2-nitrobenzonitrile (**1**, 5.0 g, 30 mmol) in isopropanol (IPA, 100 mL) were added 2-(methylamino)ethanol (**2**, 2.7 g, 36 mol) and triethylamine (TEA, 4.6 g, 45 mmol). The mixture was stirred at room temperature (rt) overnight. The resulting precipitate was collected by filtration to afford intermediate **3** as a yellow solid (6.3 g, 95.0% yield). <sup>1</sup>H NMR (400 MHz, DMSO-*d*<sub>6</sub>) $\delta$ 8.15 (d, *J* = 9.6 Hz, 1H), 7.30 (d, *J* = 2.9 Hz, 1H), 7.03 (dd, *J* = 9.7, 2.9 Hz, 1H), 4.91 – 4.82 (m, 1H), 3.61 (d, *J* = 2.7 Hz, 4H), 3.13 (s, 3H). MS(ESI<sup>+</sup>) *m/z*: 222.1 [M+H]<sup>+</sup>.

##### Synthesis of 2-amino-5-((2-hydroxyethyl)(methyl)amino)benzonitrile (intermediate 4)

To a solution of intermediate 3 (4.4 g, 20 mmol) in ethanol (50 mL), methanol (50 mL), and water (10 mL) was added iron powder (5.6 g, 100 mmol) and ammonium chloride (13.0 g, 200 mmol). The mixture was stirred at 80 o C for 4 h. The solvents were removed under reduced pressure, and the residue was diluted with ethyl acetate. The organic phase was washed sequentially with water and brine, dried over anhydrous Na2SO4, and concentrated under reduced pressure to afford compound 4 as a tan solid (3.4 g, 89.5% yield). 1 H NMR (400 MHz, DMSO-d6) δ 6.92 (dd, J = 9.1, 3.0 Hz, 1H), 6.73 (d, J = 9.1 Hz, 1H), 6.65 (d, J = 3.0 Hz, 1H), 5.22 (s, 2H), 4.56 (t, J = 5.4 Hz, 1H), 3.49 (q, J = 6.0 Hz, 2H), 3.22 (t, J = 6.2 Hz, 2H), 2.78 (s, 3H). MS (ESI) m/z: 192.1 [M+H]+ .

Synthesis of 2-((3-cyano-4-(1-fluorocyclopropane-1-carboxamido)phenyl)(methyl) amino)ethyl 1-fluorocyclopropane-1-carboxylate (intermediate 6)

To a solution of intermediate 4 (1.91 g, 10 mmol) and 1-fluorocyclopropane-1- carboxylic acid (5, 2.1 g, 20 mmol) in acetonitrile (50 mL) were added N,N,N',N'- tetramethylchloroformamidinium hexafluorophosphate (TCFH, 6.7 g, 24 mmol) and N-methylimidazole (NMI, 3.3 g, 40 mmol). The mixture was stirred at room temperature for 2 h. The solvent was removed under reduced pressure, and the residue was diluted with ethyl acetate. The organic phase was washed sequentially with water and brine, dried over anhydrous Na2SO4, and concentrated under reduced pressure. The residue was purified by silica gel column chromatography (petroleum ether/ethyl acetate = 2:1) to afford intermediate 6 as a pale yellow oil (3.1 g, 85.2% yield). 1 H NMR (400 MHz, DMSO-d6) δ 10.20 (s, 1H), 7.23 (d, J = 9.0 Hz, 1H), 7.10 (d, J = 3.0 Hz, 1H), 7.03 (dd, J = 9.0, 3.0 Hz, 1H), 4.31 (t, J = 5.3 Hz, 2H), 3.70 (t, J = 5.3 Hz,

2H), 2.93 (s, 3H), 1.49 – 1.32 (m, 4H), 1.31 – 1.23 (m, 2H), 1.21 – 1.12 (m, 2H). MS (ESI) $m/z$: 364.1 $[M+H]^+$.

Synthesis of 2-(1-fluorocyclopropyl)-6-((2-hydroxyethyl)(methyl)amino)quinazolin-4-ol (intermediate **7**)

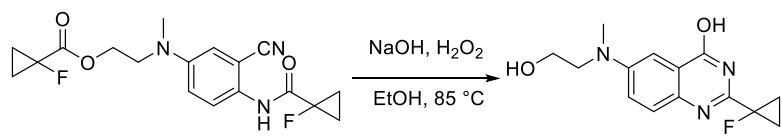

**6**

**7**

To a solution of intermediate **6** (1.45 g, 4 mmol) in ethanol (20 mL) were added NaOH (192 mg, 4.8 mmol) and $H_2O_2$ (4 mL). The mixture was stirred at 85 °C overnight. The solvent was removed under reduced pressure with the addition of silica gel. The resulting silica gel-adsorbed mixture was purified by flash column chromatography (petroleum ether/ethyl acetate = 1:1 to ethyl acetate) to afford intermediate **7** as a pale yellow solid (0.38 g, 34.2% yield). $^1H$ NMR (400 MHz, $DMSO-d_6$) $\delta$ 12.26 (s, 1H), 7.44 (d, $J$ = 9.0 Hz, 1H), 7.29 (dd, $J$ = 9.1, 3.1 Hz, 1H), 7.17 (d, $J$ = 3.0 Hz, 1H), 4.76 (s, 1H), 3.57 (t, $J$ = 6.0 Hz, 2H), 3.49 (t, $J$ = 6.0 Hz, 2H), 3.02 (s, 3H), 1.52 – 1.36 (m, 4H). MS (ESI) $m/z$: 278.1 $[M+H]^+$.

Synthesis of *tert*-butyl 4-(2-(trifluoromethoxy)phenyl)-3,6-dihydropyridine-1(2*H*)-carboxylate (intermediate **10**)

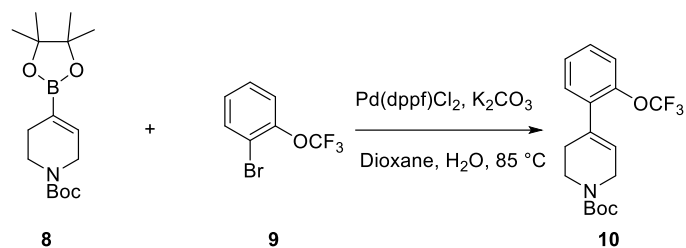

**8**

**9**

**10**

To a solution of *tert*-butyl 4-(4,4,5,5-tetramethyl-1,3,2-dioxaborolan-2-yl)-3,6-dihydropyridine-1(2*H*)-carboxylate (**8**, 1.32 g, 4.0 mmol) and 1-bromo-2-(trifluoromethoxy)benzene (**9**, 0.98 g, 4.8 mmol) in 1,4-dioxane (15 mL) and water (3 mL) were added $K_2CO_3$ (2.2 g, 16 mmol) and $Pd(dppf)Cl_2$ (0.15 g, 0.2 mmol) under a $N_2$ atmosphere. The mixture was purged with $N_2$ three times and then stirred at 85 °C overnight. The solvent was removed under reduced pressure, and the residue was diluted with ethyl acetate. The organic phase was washed sequentially with water and brine, dried over anhydrous Na<sub>2</sub>SO<sub>4</sub>, and concentrated under reduced pressure. The residue was purified by silica gel column chromatography (petroleum ether/ethyl acetate = 5:1) to afford intermediate **10** as a colorless oil (710 mg, 51.7% yield). <sup>1</sup>H NMR (400 MHz, DMSO-*d*<sub>6</sub>) $\delta$ 7.45 – 7.34 (m, 4H), 5.84 (s, 1H), 3.98 (d, *J* = 3.7 Hz, 2H), 3.51 (t, *J* = 5.6 Hz, 2H), 2.42 – 2.29 (m, 2H), 1.43 (s, 9H). MS (ESI) *m/z*: 344.1 [M+H]<sup>+</sup>.

Synthesis of 4-(2-(trifluoromethoxy)phenyl)piperidine hydrochloride (intermediate **11**)

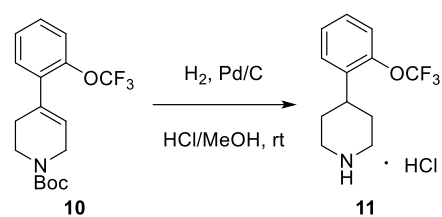

To a solution of intermediate **10** (686 mg, 2 mmol) in 2N HCl/MeOH (10 mL) was added 10% Pd/C (70 mg) under a N<sub>2</sub> atmosphere. The mixture was purged with H<sub>2</sub> three times and then stirred at room temperature overnight. The reaction mixture was filtered through a pad of Celite to remove the Pd/C, and the filtrate was concentrated under reduced pressure to afford intermediate **11** as an off-white solid (518 mg, 92.0% yield). <sup>1</sup>H NMR (400 MHz, DMSO-*d*<sub>6</sub>) $\delta$ 9.19 (br, 1H), 9.13 (br, 1H), 7.49 – 7.31 (m, 5H), 3.33 (d, *J* = 12.5 Hz, 2H), 3.20 (dtd, *J* = 12.4, 9.6, 8.9, 4.9 Hz, 1H), 3.04 (q, *J* = 12.0 Hz, 2H), 2.02 (qd, *J* = 13.1, 3.8 Hz, 2H), 1.78 (d, *J* = 13.6 Hz, 2H). MS (ESI) *m/z*: 246.1 [M+H]<sup>+</sup>.

Synthesis of 2-((2-(1-fluorocyclopropyl)-4-(4-(2-(trifluoromethoxy)phenyl)piperidin-1-yl)quinazolin-6-yl)(methyl)amino)ethan-1-ol (final product **N065**)

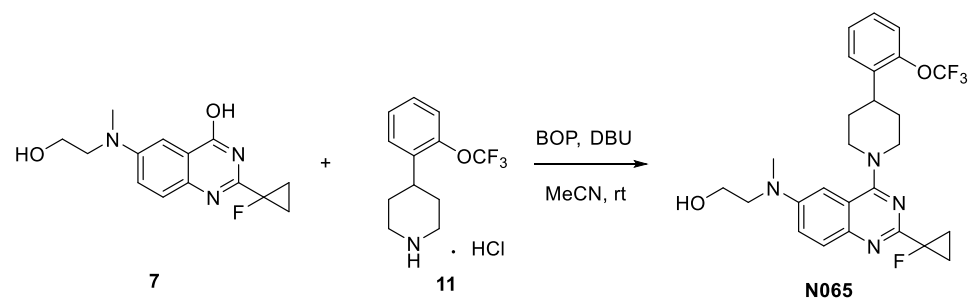

To a solution of intermediate **7** (110 mg, 0.4 mmol) and intermediate **11** (76 mg, 0.4 mmol) in acetonitrile (5 mL) were added BOP (265 mg, 0.6 mmol) and DBU (243 mg, 1.6 mmol). The mixture was stirred at room temperature for 4 h. The reaction mixture was diluted with ethyl acetate, and the organic phase was washed sequentially with water and brine, dried over anhydrous Na<sub>2</sub>SO<sub>4</sub>, and concentrated under reduced pressure. The residue was purified by silica gel column chromatography (petroleum ether/ethyl acetate = 1:1) to afford final product **N065** as a pale yellow solid (130 mg, 64.4% yield). <sup>1</sup>H NMR (400 MHz, DMSO-*d*<sub>6</sub>) δ 7.67 (d, *J* = 9.3 Hz, 1H), 7.57 – 7.51 (m, 1H), 7.47 (dd, *J* = 9.3, 2.8 Hz, 1H), 7.41 – 7.31 (m, 3H), 6.87 (d, *J* = 2.7 Hz, 1H), 4.67 (q, *J* = 5.0 Hz, 1H), 4.32 (d, *J* = 13.1 Hz, 2H), 3.63 (q, *J* = 5.7 Hz, 2H), 3.52 (t, *J* = 6.0 Hz, 2H), 3.23 – 3.17 (m, 1H), 3.12 (t, *J* = 11.9 Hz, 2H), 3.05 (s, 3H), 1.98 – 1.81 (m, 4H), 1.45 – 1.40 (m, 4H). <sup>13</sup>C NMR (101 MHz, DMSO-*d*<sub>6</sub>) δ 163.83, 157.73 (d, *J*<sub>CF</sub> = 20.7 Hz), 147.05, 146.40, 144.21, 138.23, 128.99, 128.89, 128.37, 128.31, 121.86, 121.20, 120.68 (q, *J*<sub>CF</sub> = 255.9 Hz), 116.52, 102.48, 79.08 (d, *J*<sub>CF</sub> = 219.2 Hz), 58.80, 54.97, 50.06, 39.18, 35.92, 32.25, 14.82 (d, *J*<sub>CF</sub> = 10.9 Hz). HRMS (ESI) *m/z*: calcd. for C<sub>26</sub>H<sub>29</sub>N<sub>4</sub>O<sub>2</sub>F<sub>4</sub> [M+H]<sup>+</sup>: 505.2221, found: 505.2239. HPLC purity: 99%.

**Scheme 2.** Synthesis of final compound **N191** <sup>a</sup>

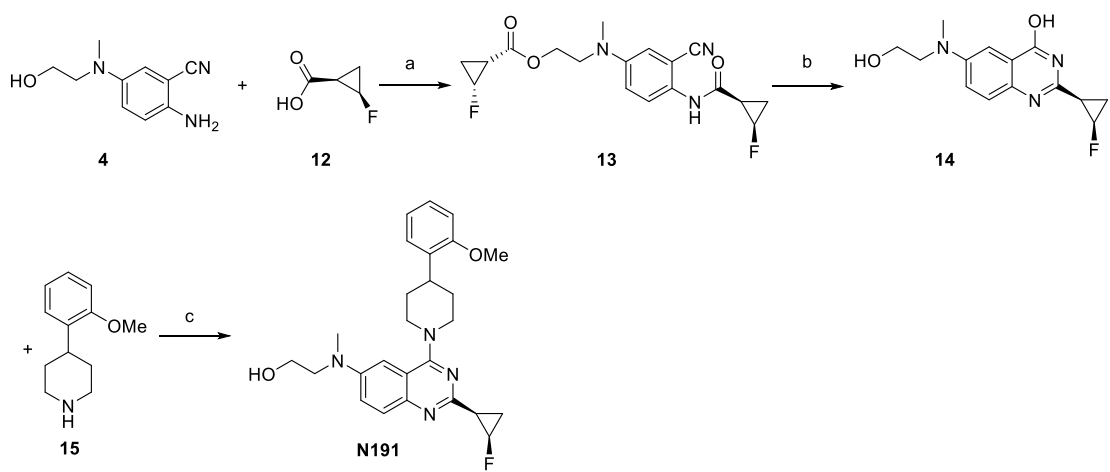

<sup>a</sup> Reagents and conditions: (a) TCFH, NMI, MeCN, rt; (b) NaOH, H<sub>2</sub>O<sub>2</sub>, EtOH, 85 °C; (c) BOP, DBU, MeCN, rt.

Synthesis of 2-((3-cyano-4-((1*R*,2*R*)-2-fluorocyclopropane-1-carboxamido)phenyl) (methyl)amino)ethyl (1*R*,2*R*)-2-fluorocyclopropane-1-carboxylate (intermediate **13**)

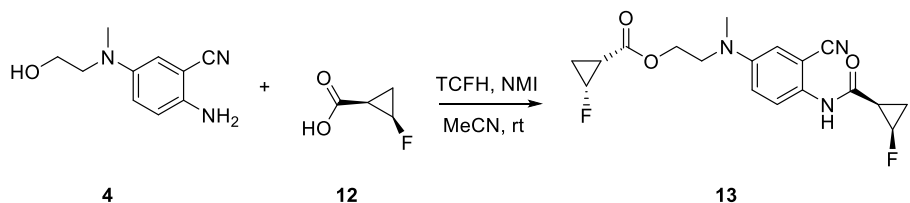

To a solution of intermediate **4** (765 mg, 4 mmol) and (1*R*,2*R*)-2-fluorocyclopropane-1-carboxylic acid (**12**, 838 mg, 8 mmol) in acetonitrile (30 mL) were added TCFH (1.97 g, 12 mmol) and NMI (3.3 g, 24 mmol). The mixture was stirred at room temperature for 2 h. The solvent was removed under reduced pressure, and the residue was diluted with ethyl acetate. The organic phase was washed sequentially with water and brine, dried over anhydrous Na<sub>2</sub>SO<sub>4</sub>, and concentrated under reduced pressure. The residue was purified by silica gel column chromatography (petroleum ether/ethyl acetate = 2:1) to afford intermediate **13** as a pale yellow solid (1.1 g, 75.8% yield). <sup>1</sup>H NMR (400 MHz, DMSO-*d*<sub>6</sub>) δ 10.04 (s, 1H), 7.27 (d, *J* = 9.0 Hz, 1H), 7.03 (d, *J* = 7.5 Hz, 2H), 5.00 (td, *J* = 6.1, 3.8 Hz, 1H), 4.84 (td, *J* = 6.2, 3.5 Hz, 1H), 4.23 (td, *J* = 5.7, 2.9 Hz, 2H), 3.65 (t, *J* = 5.6 Hz, 2H), 2.94 (s, 3H), 2.07 – 1.96 (m, 1H), 1.85 – 1.75 (m, 1H), 1.66 – 1.44 (m, 2H), 1.26 – 1.08 (m, 2H). MS (ESI) *m/z*: 364.1 [M+H]<sup>+</sup>.

Synthesis of 2-((1*R*,2*R*)-2-fluorocyclopropyl)-6-((2-hydroxyethyl)(methyl)amino)quinazolin-4-ol (intermediate **14**)

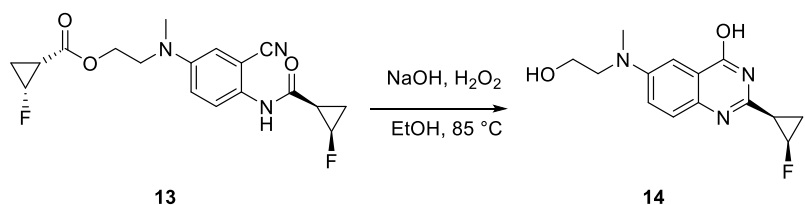

To a solution of intermediate **13** (1.45 g, 4 mmol) in ethanol (20 mL) were added NaOH (0.38 g, 9.6 mmol) and H<sub>2</sub>O<sub>2</sub> (8 mL). The mixture was stirred at 85 °C overnight. The solvent was removed under reduced pressure with the addition of silica gel. The resulting silica gel-adsorbed residue was purified by silica gel column chromatography (petroleum ether/ethyl acetate = 1:1 to ethyl acetate) to afford intermediate **14** as a pale yellow solid (0.31 g, 28.0% yield). <sup>1</sup>H NMR (400 MHz, DMSO-*d*<sub>6</sub>) δ 10.04 (s, 1H), 7.24 (d, *J* = 9.9 Hz, 1H), 7.02 – 6.94 (m, 2H), 4.92 (dtd, *J* = 66.1, 6.3, 3.7 Hz, 1H), 3.54 (t, *J* = 5.8 Hz, 2H), 3.41 (t, *J* = 6.0 Hz, 2H), 2.94 (s, 3H), 2.01 (dtd, *J* = 9.3, 6.9, 5.0 Hz, 1H), 1.60 (dtd, *J* = 23.1, 6.8, 3.6 Hz, 1H), 1.14 (ddt, *J* =

12.2, 9.1, 6.2 Hz, 1H). MS (ESI) $m/z$: 278.1 $[M+H]^+$.

Synthesis of 2-((2-((1*R*,2*R*)-2-fluorocyclopropyl)-4-(4-(2-methoxyphenyl)piperidin-1-yl)quinazolin-6-yl)(methyl)amino)ethan-1-ol (final product **N191**)

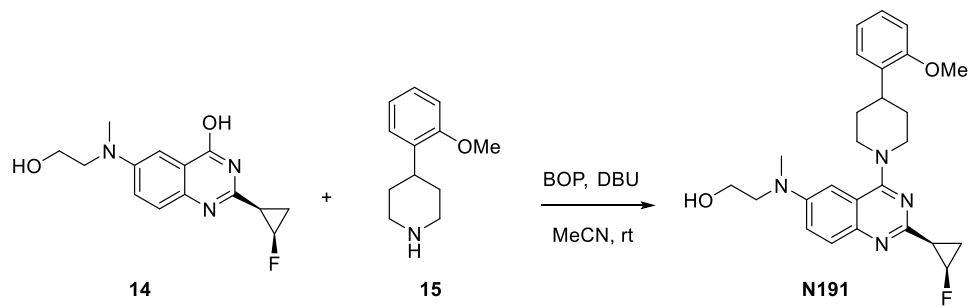

To a solution of intermediate **14** (110 mg, 0.4 mmol) and 4-(2-methoxyphenyl)piperidine (**15**, 76 mg, 0.4 mmol) in acetonitrile (5 mL) were added BOP (265 mg, 0.6 mmol) and DBU (243 mg, 1.6 mmol). The mixture was stirred at room temperature for 4 h. The reaction mixture was diluted with ethyl acetate, and the organic phase was washed sequentially with water and brine, dried over anhydrous $\text{Na}_2\text{SO}_4$, and concentrated under reduced pressure. The residue was purified by silica gel column chromatography (petroleum ether/ethyl acetate = 1:1) to afford the final product **N191** as a pale yellow solid (108 mg, 60.0% yield). $^1\text{H}$ NMR (400 MHz, $\text{DMSO}-d_6$) $\delta$ 7.61 (d, $J = 9.3$ Hz, 1H), 7.46 (dd, $J = 9.3, 2.7$ Hz, 1H), 7.25 – 7.18 (m, 2H), 6.98 (dd, $J = 8.3, 1.1$ Hz, 1H), 6.92 (td, $J = 7.4, 1.1$ Hz, 1H), 6.84 (d, $J = 2.8$ Hz, 1H), 4.98 (dtd, $J = 66.5, 6.3, 3.5$ Hz, 1H), 4.76 (s, 1H), 4.38 (t, $J = 11.0$ Hz, 2H), 3.81 (s, 3H), 3.61 (t, $J = 5.9$ Hz, 2H), 3.51 (t, $J = 5.9$ Hz, 2H), 3.28 – 3.11 (m, 4H), 3.03 (s, 3H), 2.33 (qd, $J = 7.4, 3.6$ Hz, 1H), 2.05 (dtd, $J = 22.9, 6.7, 3.5$ Hz, 1H), 1.85 (m, 3H), 1.22 (ddt, $J = 12.4, 9.6, 6.3$ Hz, 1H). $^{13}\text{C}$ NMR (101 MHz, $\text{DMSO}-d_6$) $\delta$ 163.34, 156.90, 155.75 (d, $J_{\text{CF}} = 3.5$ Hz), 146.72, 144.69, 133.77, 128.71, 127.58, 126.82, 121.65, 120.97, 116.48, 111.20, 102.43, 73.92 (d, $J_{\text{CF}} = 223.7$ Hz), 58.78, 55.79, 55.02, 50.62, 50.28, 39.20, 35.60, 32.08, 31.78, 24.70 (d, $J_{\text{CF}} = 10.6$ Hz), 10.88 (d, $J_{\text{CF}} = 9.8$ Hz). HRMS (ESI) $m/z$: calcd. for $\text{C}_{26}\text{H}_{32}\text{N}_4\text{O}_2\text{F}$ $[M+H]^+$: 451.2504, found: 451.2513. HPLC purity: 95%.

**Scheme 3.** Synthesis of final compound **N209**<sup>a</sup>

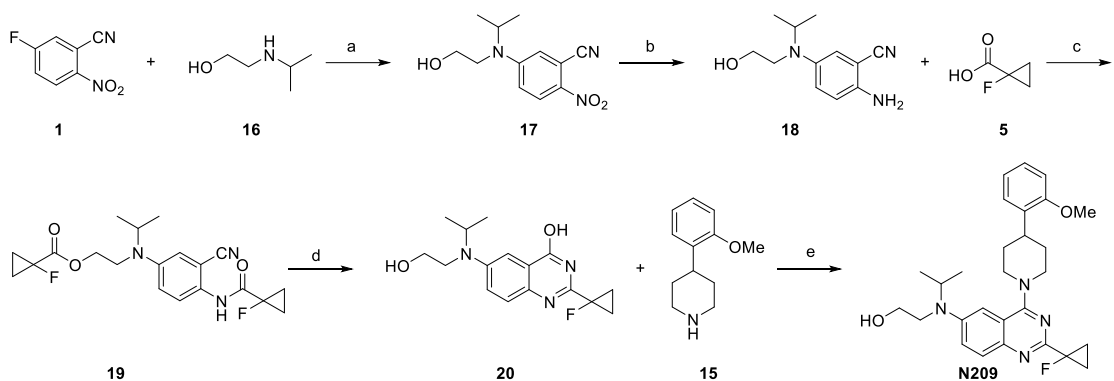

<sup>a</sup> Reagents and conditions: (a) TEA, IPA, reflux; (b) Fe, NH<sub>4</sub>Cl, EtOH, H<sub>2</sub>O, 80 °C; (c) TCFH, NMI, MeCN, rt; (d) NaOH, H<sub>2</sub>O<sub>2</sub>, EtOH, 85 °C; (e) BOP, DBU, MeCN, rt.

Synthesis of 5-((2-hydroxyethyl)(isopropyl)amino)-2-nitrobenzonitrile (intermediate **17**)

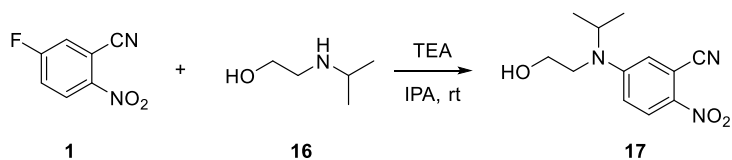

To a solution of 5-fluoro-2-nitrobenzonitrile (**1**, 1.2 g, 8 mmol) in isopropanol (10 mL) were added triethylamine (1.2 g, 12 mmol) and 2-(isopropylamino)ethan-1-ol (**16**, 1.0 g, 9.6 mmol). The mixture was stirred at reflux for 2 h. The solvent was removed under reduced pressure with the addition of silica gel. The resulting silica gel-adsorbed residue was purified by silica gel column chromatography (petroleum ether/ethyl acetate = 1:1) to afford intermediate **17** as a yellow solid (0.85 g, 42.6% yield). <sup>1</sup>H NMR (400 MHz, DMSO-*d*<sub>6</sub>) δ 8.16 (d, *J* = 9.7 Hz, 1H), 7.39 (d, *J* = 2.9 Hz, 1H), 7.15 (dd, *J* = 9.8, 3.0 Hz, 1H), 4.97 (t, *J* = 5.6 Hz, 1H), 4.31 (hept, *J* = 6.6 Hz, 1H), 3.57 – 3.50 (m, 2H), 3.46 (t, *J* = 6.2 Hz, 2H), 1.19 (d, *J* = 6.5 Hz, 6H). MS (ESI<sup>+</sup>) *m/z*: 250.1 [M+H]<sup>+</sup>.

Synthesis of 2-amino-5-((2-hydroxyethyl)(isopropyl)amino)benzonitrile (intermediate **18**)

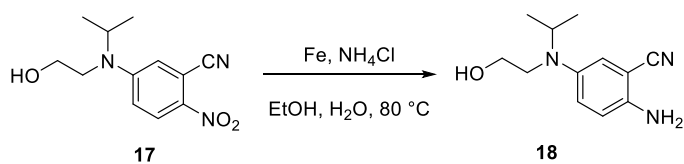

To a solution of intermediate **17** (0.50 g, 2 mmol) in ethanol (20 mL) and water (4 mL)

were added iron powder (0.56 g, 10 mmol) and ammonium chloride (1.3 g, 20 mmol). The mixture was stirred at 80 °C for 4 h. The solvent was removed under reduced pressure, and the residue was diluted with ethyl acetate. The organic phase was washed sequentially with water and brine, dried over anhydrous Na<sub>2</sub>SO<sub>4</sub>, and concentrated under reduced pressure to afford intermediate **18** as a pale yellow oil (0.42 g, 95.9% yield). <sup>1</sup>H NMR (400 MHz, DMSO-*d*<sub>6</sub>) δ 6.99 (dd, *J* = 9.1, 2.9 Hz, 1H), 6.78 (d, *J* = 2.9 Hz, 1H), 6.73 (d, *J* = 9.1 Hz, 1H), 5.34 (s, 2H), 4.59 (t, *J* = 5.6 Hz, 1H), 3.70 (hept, *J* = 6.5 Hz, 1H), 3.38 – 3.33 (m, 2H), 3.04 (t, *J* = 7.1 Hz, 2H), 1.02 (d, *J* = 6.5 Hz, 6H). MS (ESI) *m/z*: 220.1 [M+H]<sup>+</sup>.

Synthesis of 2-((3-cyano-4-((1-fluorocyclopropane-1-carboxamido)phenyl)(isopropyl)amino)ethyl 1-fluorocyclopropane-1-carboxylate (intermediate **19**)

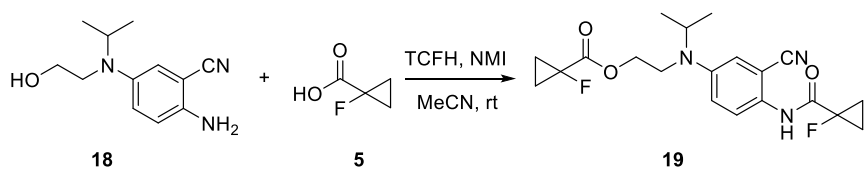

To a solution of intermediate **18** (0.22 g, 1.0 mmol) and 1-fluorocyclopropane-1-carboxylic acid (**5**, 0.23 g, 2.2 mmol) in acetonitrile (10 mL) were added TCFH (0.67 g, 2.4 mmol) and NMI (0.33 g, 4.0 mmol). The mixture was stirred at room temperature for 2 h. The solvent was removed under reduced pressure, and the residue was diluted with ethyl acetate. The organic phase was washed sequentially with water and brine, dried over anhydrous Na<sub>2</sub>SO<sub>4</sub>, and concentrated under reduced pressure. The residue was purified by silica gel column chromatography (petroleum ether/ethyl acetate = 2:1) to afford intermediate **19** as a pale yellow oil (0.32 g, 81.6% yield). <sup>1</sup>H NMR (400 MHz, DMSO-*d*<sub>6</sub>) δ 10.24 (s, 1H), 7.24 (d, *J* = 9.0 Hz, 2H), 7.21 (d, *J* = 2.9 Hz, 2H), 7.15 (dd, *J* = 9.1, 3.1 Hz, 1H), 4.23 (t, *J* = 6.2 Hz, 2H), 4.11 (hept, *J* = 6.6 Hz, 1H), 3.51 (t, *J* = 6.2 Hz, 2H), 1.50 – 1.39 (m, 2H), 1.32 – 1.25 (m, 2H), 1.14 (d, *J* = 6.5 Hz, 6H). MS (ESI) *m/z*: 392.2 [M+H]<sup>+</sup>.

Synthesis of 2-(1-fluorocyclopropyl)-6-((2-hydroxyethyl)(isopropyl)amino)quinazolin-4-ol (intermediate **20**)

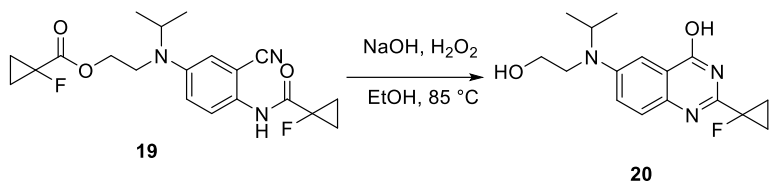

To a solution of intermediate **19** (0.39 g, 1.0 mmol) in ethanol (5 mL) were added NaOH (0.12 g, 3.0 mmol) and H<sub>2</sub>O<sub>2</sub> (3 mL). The mixture was stirred at 85 °C overnight. The solvent was removed under reduced pressure with the addition of silica gel. The resulting silica gel-adsorbed residue was purified by flash column chromatography (petroleum ether/ethyl acetate = 1:1 to ethyl acetate) to afford intermediate **20** as a pale yellow solid (0.12 g, 39.3% yield). <sup>1</sup>H NMR (400 MHz, DMSO-*d*<sub>6</sub>) δ 12.24 (s, 1H), 7.44 (d, *J* = 9.1 Hz, 1H), 7.34 (dd, *J* = 9.1, 3.1 Hz, 1H), 7.29 (d, *J* = 3.0 Hz, 1H), 4.85 (t, *J* = 5.5 Hz, 1H), 4.14 (p, *J* = 6.6 Hz, 1H), 3.51 (q, *J* = 7.1 Hz, 2H), 3.30 (t, *J* = 7.2 Hz, 2H), 1.52 – 1.37 (m, 4H), 1.17 (d, *J* = 6.5 Hz, 6H). MS (ESI) *m/z*: 306.2 [M+H]<sup>+</sup>.

Synthesis of 2-((2-(1-fluorocyclopropyl)-4-(4-(2-methoxyphenyl)piperidin-1-yl)quinazolin-6-yl)(isopropyl)amino)ethan-1-ol (final product **N209**)

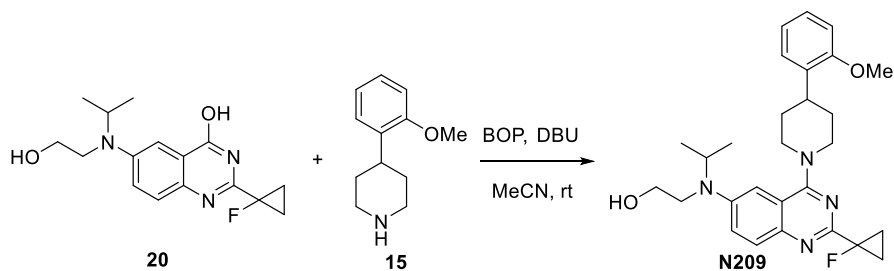

To a solution of intermediate **20** (122 mg, 0.4 mmol) and 4-(2-methoxyphenyl)piperidine (**15**, 76 mg, 0.4 mmol) in acetonitrile (5 mL) were added BOP (265 mg, 0.6 mmol) and DBU (243 mg, 1.6 mmol). The mixture was stirred at room temperature for 4 h. The reaction mixture was diluted with ethyl acetate, and the organic phase was washed sequentially with water and brine, dried over anhydrous Na<sub>2</sub>SO<sub>4</sub>, and concentrated under reduced pressure. The residue was purified by silica gel column chromatography (petroleum ether/ethyl acetate = 1:1) to afford the final product **N209** as a pale yellow solid (98 mg, 51.2% yield). <sup>1</sup>H NMR (400 MHz, DMSO-*d*<sub>6</sub>) δ 7.69 (d, *J* = 9.3 Hz, 1H), 7.52 (dd, *J* = 9.4, 2.5 Hz, 1H), 7.26 – 7.16 (m, 2H), 7.00 – 6.91 (m, 3H), 4.85 (t, *J* = 5.3 Hz, 1H), 4.27 (d, *J* = 12.9 Hz,

2H), 4.13 (hept, 6.4 Hz, 1H), 3.81 (s, 3H), 3.55 (q, $J = 6.5$ Hz, 2H), 3.32 (d, $J = 6.9$ Hz, 2H), 3.20 (h, $J = 7.6, 6.4$ Hz, 1H), 3.12 – 3.04 (m, 2H), 1.93 – 1.76 (m, 4H), 1.47 – 1.39 (m, 4H), 1.18 (d, $J = 6.5$ Hz, 6H). $^{13}\text{C}$ NMR (101 MHz, DMSO-$d_6$) $\delta$ 163.97, 157.90 (d, $J_{\text{CF}} = 20.7$ Hz), 156.93, 146.38, 144.29, 133.67, 129.02, 127.61, 126.84, 122.42, 120.98, 116.59, 111.20, 103.57, 79.07 (d, $J_{\text{CF}} = 218.5$ Hz), 60.07, 55.78, 50.47, 48.32, 46.87, 35.68, 31.89, 20.14, 14.82 (d, $J_{\text{CF}} = 10.8$ Hz). HRMS (ESI) $m/z$: calcd. for $\text{C}_{28}\text{H}_{35}\text{FN}_4\text{O}_2$ $[\text{M}+\text{H}]^+$: 479.2817, found: 479.2796. HPLC purity: 95%.

**Scheme 4.** Synthesis of final compound **N253**<sup>a</sup>

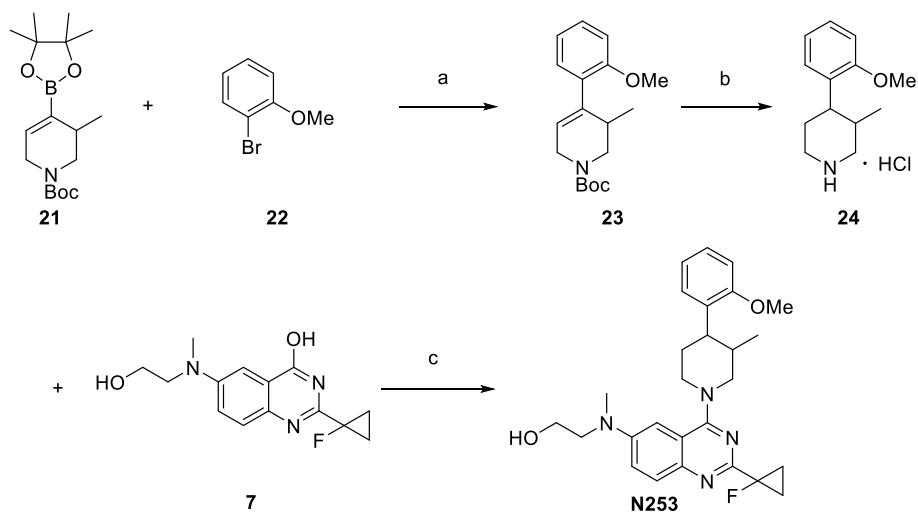

<sup>a</sup> Reagents and conditions: (a) $\text{Pd}(\text{dppf})\text{Cl}_2$, $\text{K}_2\text{CO}_3$, Dioxane, $\text{H}_2\text{O}$, $85^\circ\text{C}$; (b) $\text{H}_2$, $\text{Pd/C}$, $\text{HCl/MeOH}$, rt; (c) BOP, DBU, MeCN, rt.

**Synthesis of *tert*-butyl 4-(2-methoxyphenyl)-3-methyl-3,6-dihydropyridine-1(2H)-carboxylate (intermediate **23**)**

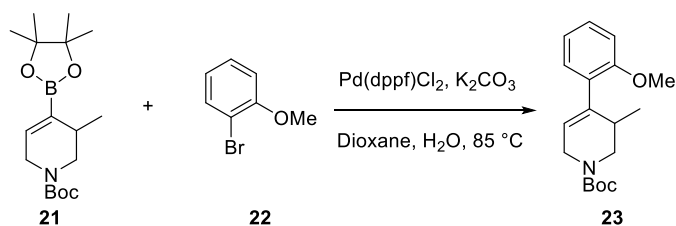

To a solution of *tert*-butyl 3-methyl-4-(4,4,5,5-tetramethyl-1,3,2-dioxaborolan-2-yl)-3,6-dihydropyridine-1(2H)-carboxylate (**21**, 97 mg, 0.3 mmol) and 1-bromo-2-methoxybenzene (**22**, 67 mg, 0.36 mmol) in 1,4-dioxane (10 mL) and water (2 mL) were added $\text{K}_2\text{CO}_3$ (83 mg, 0.6 mmol) and $\text{Pd}(\text{dppf})\text{Cl}_2$ (22 mg, 0.03 mmol) under a $\text{N}_2$ atmosphere. The mixture was purged with $\text{N}_2$ three times and then stirred at 85 °C overnight. The solvent was removed under reduced pressure, and the residue was diluted with ethyl acetate. The organic phase was washed sequentially with water and brine, dried over anhydrous Na<sub>2</sub>SO<sub>4</sub>, and concentrated under reduced pressure. The residue was purified by silica gel column chromatography (petroleum ether/ethyl acetate = 5:1) to afford intermediate **23** as a colorless oil (50 mg, 55.6% yield). <sup>1</sup>H NMR (400 MHz, DMSO-*d*<sub>6</sub>) δ 7.40 (ddd, *J* = 8.8, 7.5, 1.7 Hz, 1H), 7.24 (dd, *J* = 7.5, 1.7 Hz, 1H), 7.09 (dd, *J* = 8.4, 1.0 Hz, 1H), 7.00 (td, *J* = 7.5, 1.0 Hz, 1H), 5.84 (s, 1H), 3.86 (dd, *J* = 12.9, 4.4 Hz, 1H), 3.81 (s, 3H), 3.72 (dd, *J* = 12.9, 5.7 Hz, 1H), 3.48 – 3.38 (m, 1H), 3.34 – 3.28 (m, 1H), 3.21 – 3.10 (m, 1H), 1.48 (s, 9H), 0.90 (d, *J* = 7.0 Hz, 3H). MS (ESI) *m/z*: 304.2 [M+H]<sup>+</sup>.

Synthesis of 4-(2-methoxyphenyl)-3-methylpiperidine hydrochloride (intermediate **24**)

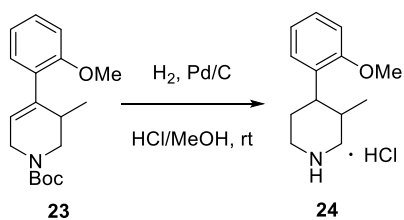

To a solution of intermediate **23** (45 mg, 0.15 mmol) in 2N HCl/MeOH (5 mL) was added 10% Pd/C (5 mg) under a N<sub>2</sub> atmosphere. The mixture was purged with H<sub>2</sub> three times and then stirred at room temperature overnight. The reaction mixture was filtered through a pad of Celite to remove the Pd/C, and the filtrate was concentrated under reduced pressure to afford intermediate **24** as an off-white solid (33 mg, 90.9% yield). <sup>1</sup>H NMR (400 MHz, DMSO-*d*<sub>6</sub>) δ 9.30 (s, 1H), 8.61 (s, 1H), 7.23 (td, *J* = 8.4, 7.9, 1.7 Hz, 1H), 7.04 – 6.91 (m, 3H), 3.79 (s, 3H), 3.42 – 3.24 (m, 4H), 3.01 (q, *J* = 12.0 Hz, 1H), 2.39 – 2.30 (m, 1H), 2.21 (qd, *J* = 13.3, 4.1 Hz, 1H), 1.66 – 1.54 (m, 1H), 0.74 (d, *J* = 7.4 Hz, 3H). MS (ESI) *m/z*: 206.1 [M+H]<sup>+</sup>.

Synthesis of 2-((2-(1-fluorocyclopropyl)-4-(4-(2-methoxyphenyl)-3-methylpiperidin-1-yl)quinazolin-6-yl)(methyl)amino)ethan-1-ol (final product **N253**)

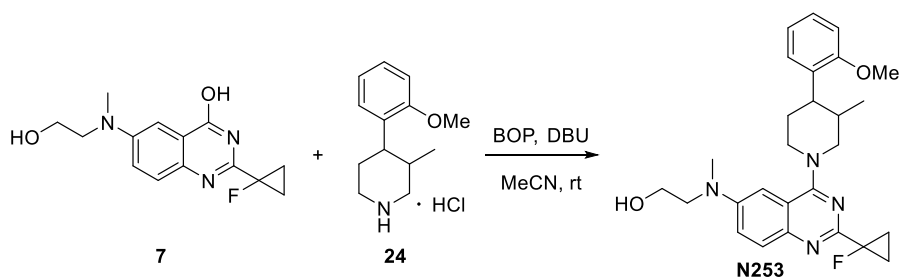

To a solution of intermediate **7** (33 mg, 0.12 mmol) and intermediate **24** (29 mg, 0.12 mmol) in acetonitrile (5 mL) were added BOP (80 mg, 0.18 mmol) and DBU (73 mg, 0.48 mmol). The mixture was stirred at room temperature for 4 h. The reaction mixture was diluted with ethyl acetate, and the organic phase was washed sequentially with water and brine, dried over anhydrous Na<sub>2</sub>SO<sub>4</sub>, and concentrated under reduced pressure. The residue was purified by silica gel column chromatography (petroleum ether/ethyl acetate = 1:1) to afford final product **N253** as a pale yellow solid (36 mg, 64.6% yield). <sup>1</sup>H NMR (400 MHz, DMSO-*d*<sub>6</sub>) δ 7.69 (d, *J* = 9.3 Hz, 1H), 7.48 (dd, *J* = 9.4, 2.7 Hz, 1H), 7.22 (td, *J* = 7.8, 1.7 Hz, 1H), 7.12 (dd, *J* = 7.7, 1.7 Hz, 1H), 6.99 (dd, *J* = 8.4, 1.2 Hz, 1H), 6.94 (td, *J* = 7.4, 1.2 Hz, 1H), 6.90 (d, *J* = 2.8 Hz, 1H), 4.79 (s, 1H), 4.34 – 4.21 (m, 2H), 3.80 (s, 3H), 3.61 (t, *J* = 5.8 Hz, 2H), 3.55 – 3.50 (m, 2H), 3.28 (d, *J* = 12.7 Hz, 2H), 3.12 (t, *J* = 12.3 Hz, 1H), 3.05 (s, 3H), 2.42 – 2.25 (m, 2H), 1.59 (dd, *J* = 13.1, 3.1 Hz, 1H), 1.51 – 1.40 (m, 4H), 0.74 (d, *J* = 6.9 Hz, 3H). <sup>13</sup>C NMR (101 MHz, DMSO-*d*<sub>6</sub>) δ 164.81, 157.76 (d, *J*<sub>CF</sub> = 20.7 Hz), 157.02, 147.01, 144.19, 131.89, 128.93, 127.91, 127.59, 121.77, 120.46, 116.67, 111.00, 102.47, 79.07 (d, *J*<sub>CF</sub> = 218.3 Hz), 58.76, 56.39, 55.75, 54.95, 50.37, 39.42, 38.76, 32.21, 24.03, 14.89 (d, *J*<sub>CF</sub> = 11.3 Hz), 14.77 (d, *J*<sub>CF</sub> = 11.5 Hz), 12.45. HRMS (ESI) *m/z*: calcd. for C<sub>27</sub>H<sub>33</sub>FN<sub>4</sub>O<sub>2</sub> [M+H]<sup>+</sup>: 465.2660, found: 465.2635. HPLC purity: 90%.

To a solution of intermediate **7** (33 mg, 0.12 mmol) and intermediate **24** (29 mg, 0.12 mmol) in acetonitrile (5 mL) were added BOP (80 mg, 0.18 mmol) and DBU (73 mg,

0.48 mmol). The mixture was stirred at room temperature for 4 h. The reaction

mixture was diluted with ethyl acetate, and the organic phase was washed sequentially

with water and brine, dried over anhydrous $\text{Na}_2\text{SO}_4$, and concentrated under reduced

pressure. The residue was purified by silica gel column chromatography (petroleum

ether/ethyl acetate = 1:1) to afford final product **N253** as a pale yellow solid (36 mg,

64.6% yield). $^1\text{H}$ NMR (400 MHz, $\text{DMSO}-d_6$) $\delta$ 7.69 (d, $J = 9.3$ Hz, 1H), 7.48 (dd, $J$

= 9.4, 2.7 Hz, 1H), 7.22 (td, $J = 7.8, 1.7$ Hz, 1H), 7.12 (dd, $J = 7.7, 1.7$ Hz, 1H), 6.99

(dd, $J = 8.4, 1.2$ Hz, 1H), 6.94 (td, $J = 7.4, 1.2$ Hz, 1H), 6.90 (d, $J = 2.8$ Hz, 1H), 4.79

(s, 1H), 4.34 – 4.21 (m, 2H), 3.80 (s, 3H), 3.61 (t, $J = 5.8$ Hz, 2H), 3.55 – 3.50 (m,

2H), 3.28 (d, $J = 12.7$ Hz, 2H), 3.12 (t, $J = 12.3$ Hz, 1H), 3.05 (s, 3H), 2.42 – 2.25 (m,

2H), 1.59 (dd, $J = 13.1, 3.1$ Hz, 1H), 1.51 – 1.40 (m, 4H), 0.74 (d, $J = 6.9$ Hz, 3H).

<sup>13</sup>C NMR (101 MHz, DMSO-*d*<sub>6</sub>) δ 164.81, 157.76 (d, *J*<sub>CF</sub> = 20.7 Hz), 157.02, 147.01,

144.19, 131.89, 128.93, 127.91, 127.59, 121.77, 120.46, 116.67, 111.00, 102.47,

79.07 (d, $J_{\text{CF}} = 218.3$ Hz), 58.76, 56.39, 55.75, 54.95, 50.37, 39.42, 38.76, 32.21,

24.03, 14.89 (d, $J_{\text{CF}} = 11.3$ Hz), 14.77 (d, $J_{\text{CF}} = 11.5$ Hz), 12.45. HRMS (ESI) $m/z$:

calcd. for $\text{C}_{27}\text{H}_{33}\text{FN}_4\text{O}_2$ $[\text{M}+\text{H}]^+$: 465.2660, found: 465.2635. HPLC purity: 90%.

376 **NMR, HRMS and HPLC spectra of final compounds N065, N191, N209 and**
 377 **N253**

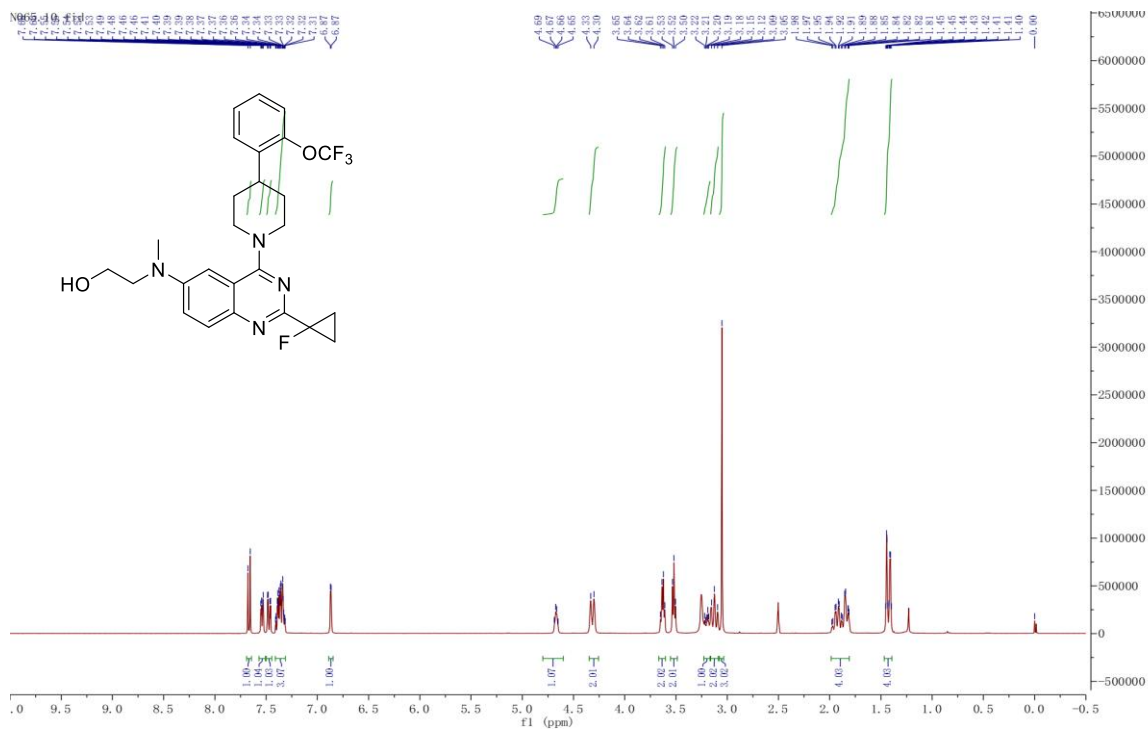

378
379 <sup>1</sup>H NMR spectrum of compound N065
380
381

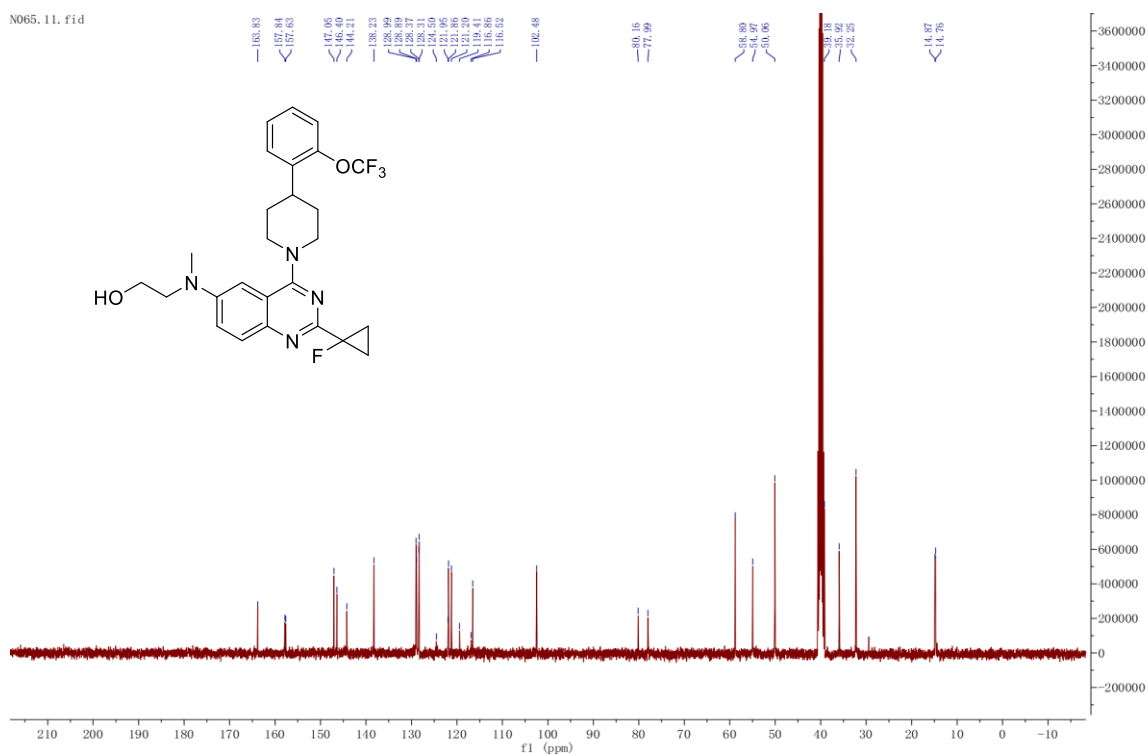

382
383 <sup>13</sup>C NMR spectrum of compound N065

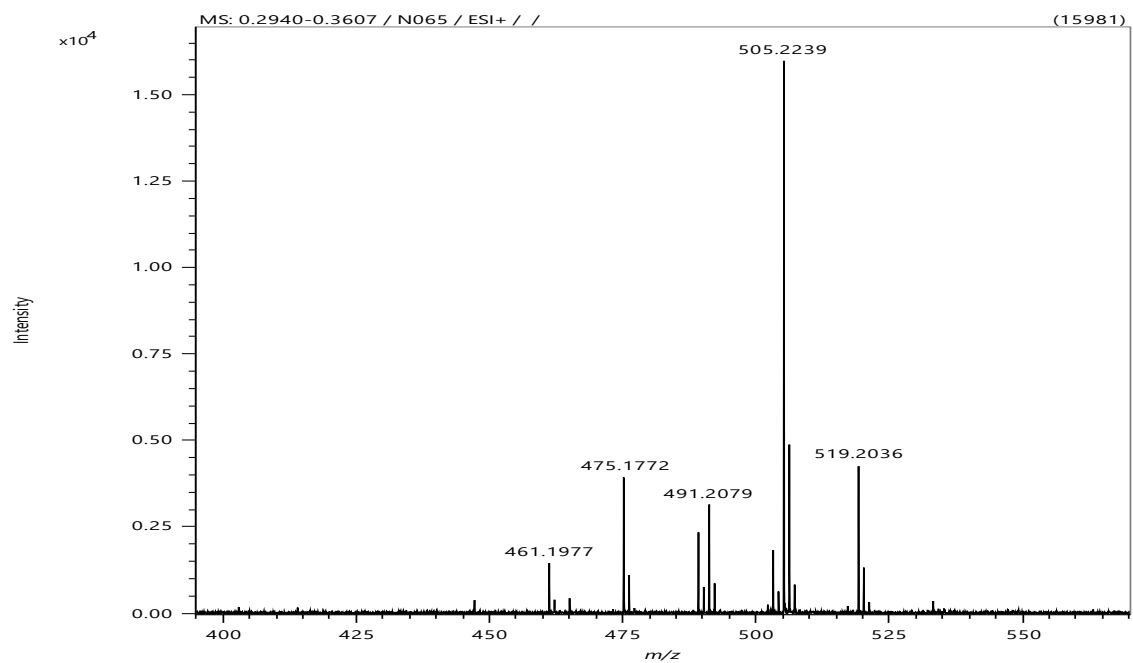

HRMS spectrum of compound N065

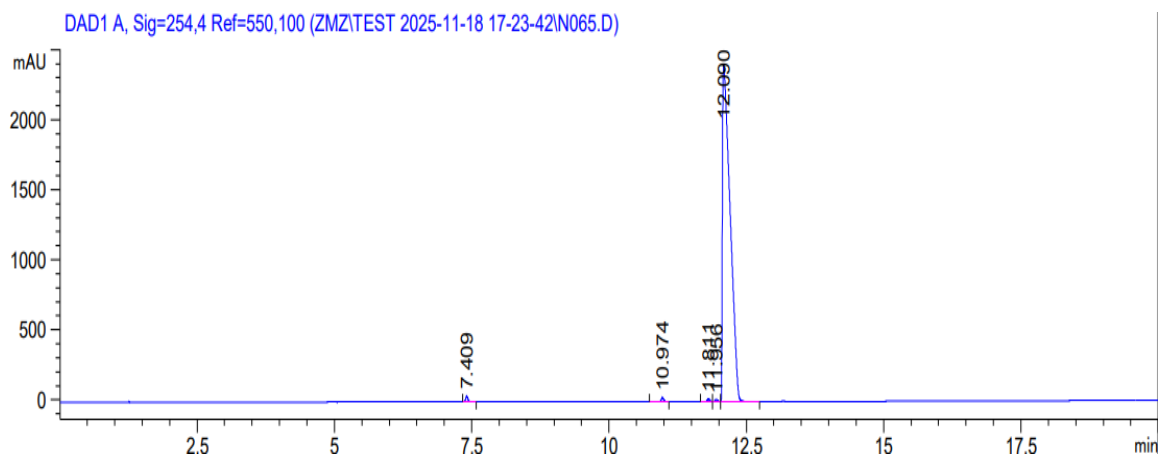

| Peak # | RetTime [min] | Type | Width [min] | Area [mAU*s] | Height [mAU] | Area % |
| --- | --- | --- | --- | --- | --- | --- |
| 1 | 7.409 | BB | 0.0416 | 110.04398 | 41.24390 | 0.4653 |
| 2 | 10.974 | BB | 0.0483 | 89.36971 | 28.72192 | 0.3779 |
| 3 | 11.811 | BV | 0.0479 | 61.78386 | 19.79717 | 0.2612 |
| 4 | 11.956 | VV | 0.0451 | 50.16656 | 17.15553 | 0.2121 |
| 5 | 12.090 | VB | 0.1395 | 2.33389e4 | 2398.61108 | 98.6835 |

HPLC spectrum of compound N065

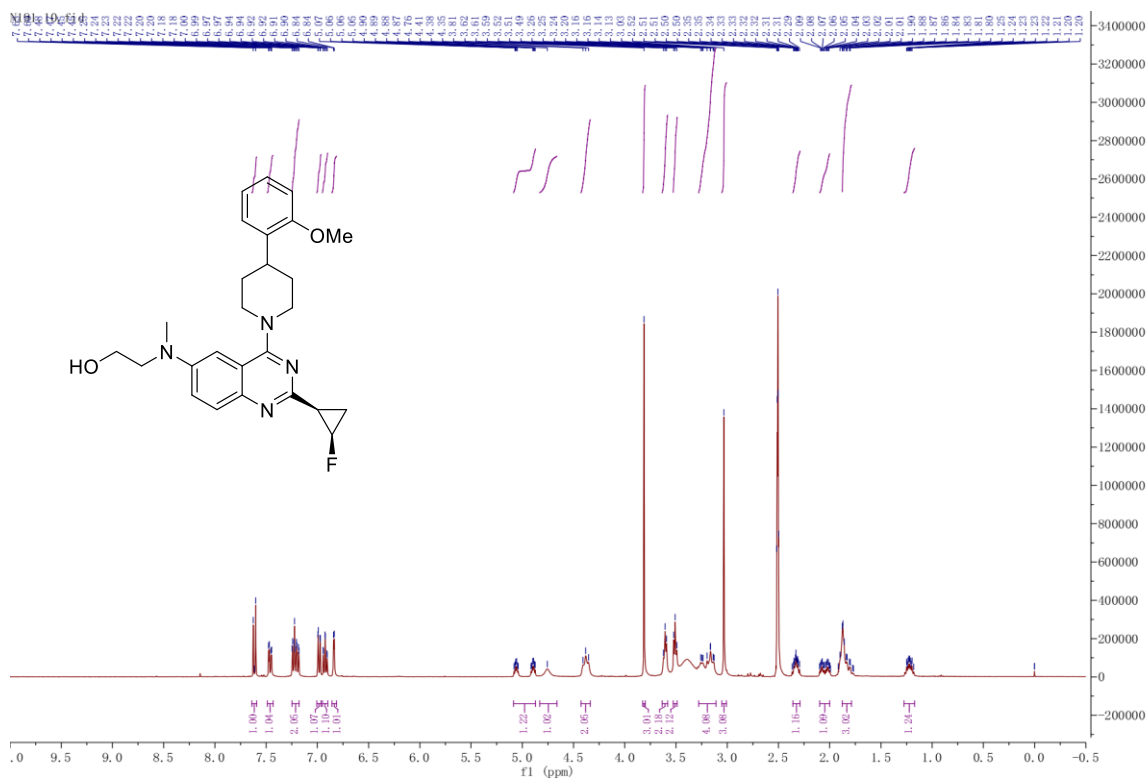

<sup>1</sup>H NMR spectrum of compound N191

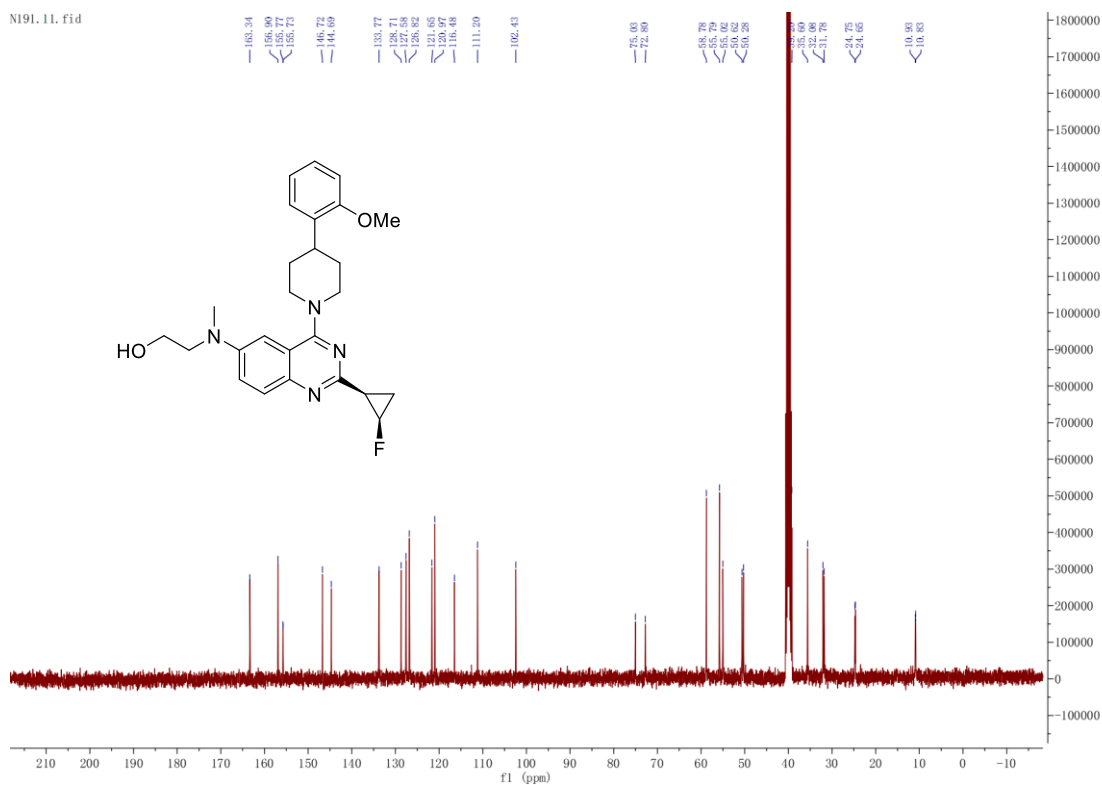

<sup>13</sup>C NMR spectrum of compound N191

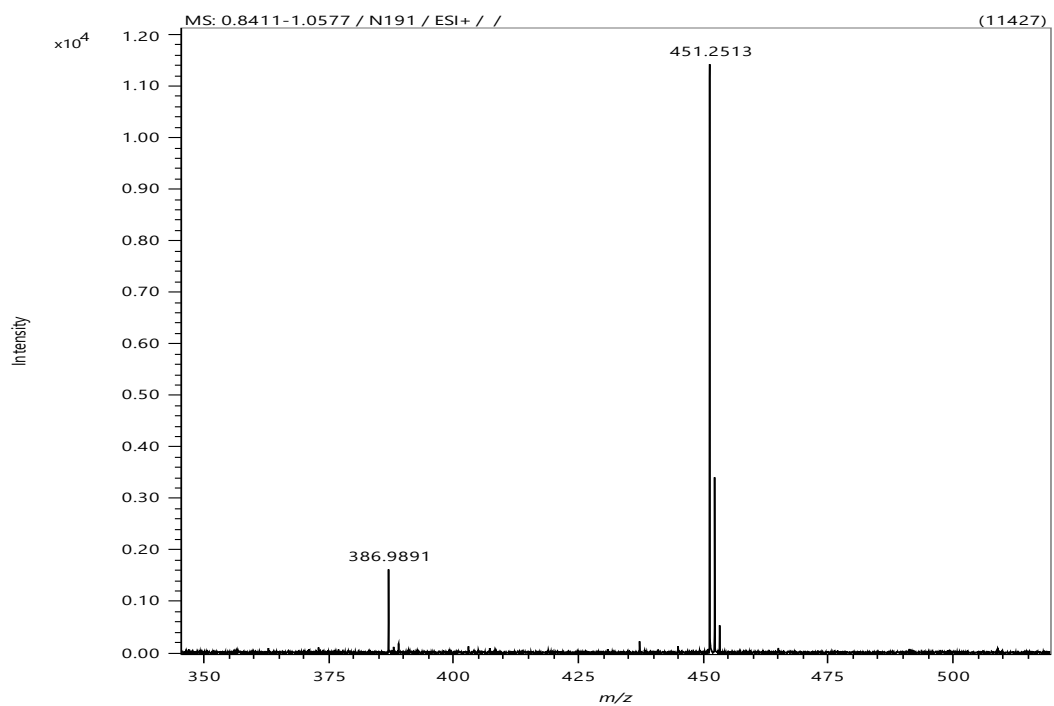

HRMS spectrum of compound N191

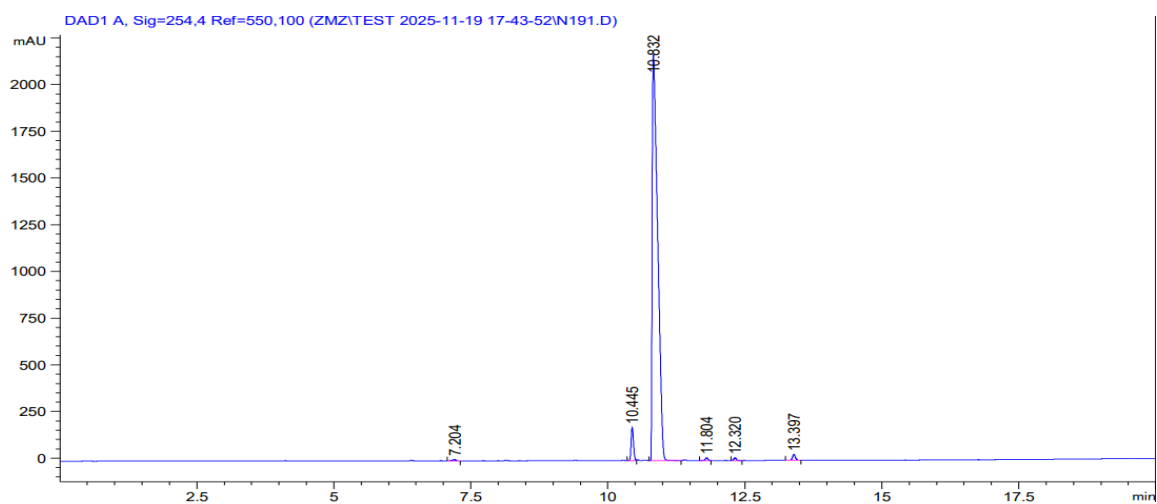

| Peak # | RetTime [min] | Type | Width [min] | Area [mAU*s] | Height [mAU] | Area % |
| --- | --- | --- | --- | --- | --- | --- |
| 1 | 7.204 | BB | 0.0593 | 33.00202 | 8.05110 | 0.2101 |
| 2 | 10.445 | VV | 0.0463 | 527.81628 | 176.93069 | 3.3599 |
| 3 | 10.832 | BB | 0.1063 | 1.49378e4 | 2171.24609 | 95.0879 |
| 4 | 11.804 | VV | 0.0532 | 48.91503 | 13.83471 | 0.3114 |
| 5 | 12.320 | VV | 0.0511 | 46.63409 | 14.09264 | 0.2969 |
| 6 | 13.397 | BB | 0.0564 | 115.29121 | 31.71788 | 0.7339 |

HPLC spectrum of compound N191

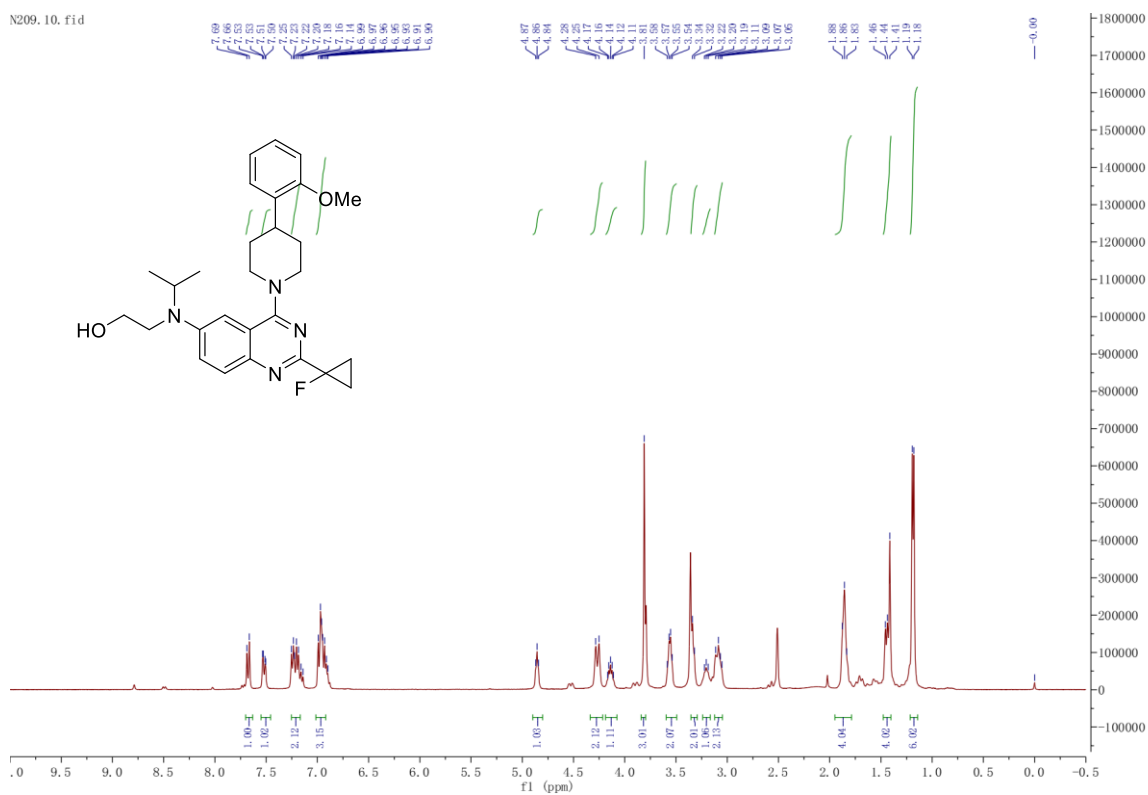

<sup>1</sup>H NMR spectrum of compound N209

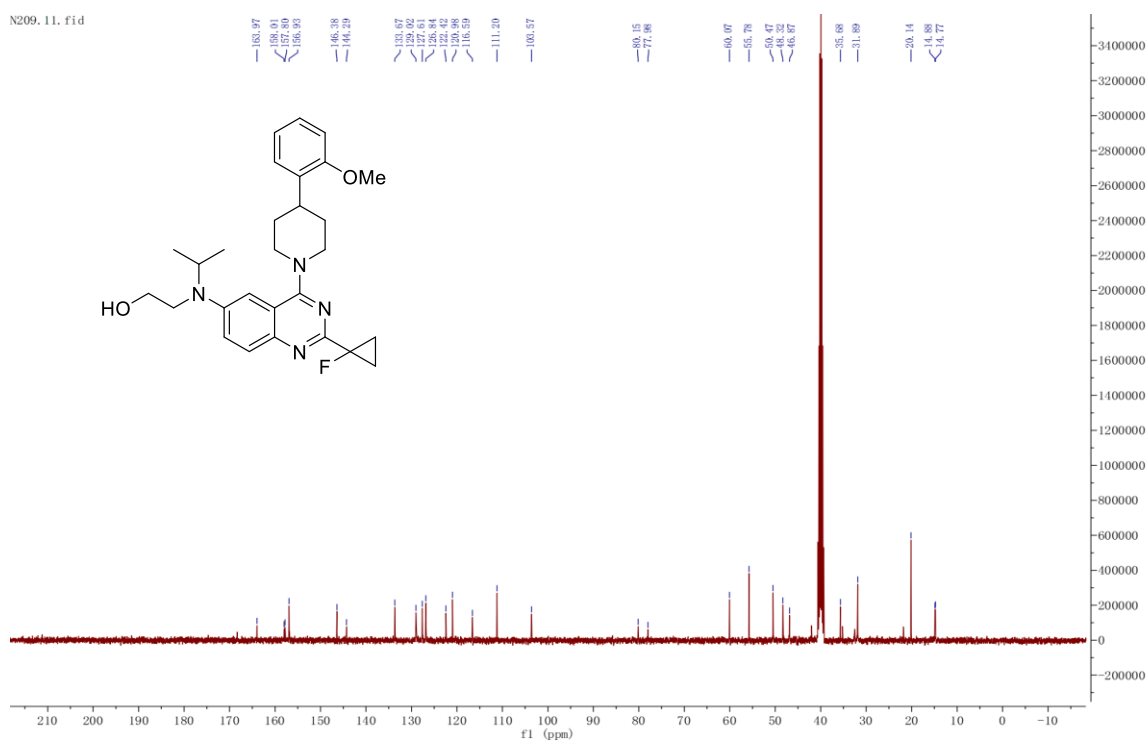

<sup>13</sup>C NMR spectrum of compound N209

HRMS spectrum of compound N209

| Peak # | RetTime [min] | Type | Width [min] | Area [mAU*s] | Height [mAU] | Area % |
| --- | --- | --- | --- | --- | --- | --- |
| 1 | 8.342 | BB | 0.0444 | 36.01479 | 12.56520 | 0.6882 |
| 2 | 9.100 | BB | 0.0541 | 135.53703 | 38.91961 | 2.5898 |
| 3 | 9.807 | BB | 0.0521 | 19.87400 | 5.93741 | 0.3798 |
| 4 | 10.444 | BB | 0.0488 | 17.10046 | 5.49857 | 0.3268 |
| 5 | 11.067 | VB | 0.0499 | 32.86466 | 10.12265 | 0.6280 |
| 6 | 11.457 | BB | 0.0856 | 4947.73389 | 887.12366 | 94.5409 |
| 7 | 12.215 | VB | 0.0514 | 17.86478 | 5.35915 | 0.3414 |

HPLC spectrum of compound N209

427

428

429

HRMS spectrum of compound N253

430

431

432

| Peak # | RetTime [min] | Type | Width [min] | Area [mAU*s] | Height [mAU] | Area % |
| --- | --- | --- | --- | --- | --- | --- |
| 1 | 3.772 | VB | 0.0381 | 40.85174 | 14.19983 | 0.3911 |
| 2 | 8.358 | VV | 0.0533 | 41.21448 | 11.92830 | 0.3945 |
| 3 | 10.851 | BB | 0.0522 | 61.68922 | 18.10708 | 0.5905 |
| 4 | 11.165 | VB | 0.0529 | 345.84116 | 98.71672 | 3.3106 |
| 5 | 11.368 | BV | 0.0527 | 491.85608 | 146.42406 | 4.7083 |
| 6 | 11.614 | VV | 0.1155 | 9365.51660 | 1180.86255 | 89.6520 |
| 7 | 12.607 | VB | 0.0622 | 99.55255 | 24.07893 | 0.9530 |

433

434

HPLC spectrum of compound N253

435

436

437

438 HPLC spectrum of TAMRA-PEG6-Arg-Arg-Pro-Tyr-Ile-Leu (provided by TGpeptide company)

454 **Table S1. Curve parameters and statistical comparisons presented in Figure 4 and Figure**
455 **S23 -Supporting Figure 4 and Figure S23.**

456

| Figure | Panel | Experiment | Statistical Test | F Statistics | Multiple Comparisons | N |
| --- | --- | --- | --- | --- | --- | --- |
| 4 | B | Core body temperature - N065, SBI-553 | Two-way, repeated measures ANOVA | $F_{\text{interaction}}(12, 208) = 7.989, p < 0.0001$<br>$F_{\text{time}}(1.811, 94.20) = 92.58, p < 0.0001$<br>$F_{\text{treatment}}(3, 52) = 23.28, p < 0.0001$ | *$p < 0.05$, **$p < 0.01$, ***$p < 0.001$, ****$p < 0.0001$, as compared to Vehicle at doses and indicated time points | 10-16 mice/treatment |
| 4 | C | Open field test - N065, SBI-553 | Two-way, repeated measures ANOVA | $F_{\text{time}}(4.337, 125.5) = 283.4, p < 0.0001$<br>$F_{\text{treatment}}(2, 29) = 2.317, p = 0.1166$<br>$F_{\text{interaction}}(34, 492) = 1.129, p = 0.2868$ | *$p < 0.05$, as compared to Vehicle at doses and indicated time points | 10-11 mice/treatment |
| 4 | D | Pole descent test - N065, SBI-553 | One-way ANOVA on ranks | $F(2, 28) = 0.9628, p = 0.3941$ | No treatment effect | 10-11 mice/treatment |
| S23 | A | Wire hang test - N065, SBI-553 | One-way ANOVA on ranks | All the values are the same, so no statistical test is needed. | No treatment effect | 10-11 mice/treatment |
| 4 | E | Contact righting - N065, SBI-553 | One-way ANOVA on ranks | Kruskal-Wallis statistic=2.035, $p = 0.3615$ | No treatment effect | 10-11 mice/treatment |
| 4 | F | Systolic blood pressure - N065, SBI-553 | One-way ANOVA on ranks | Kruskal-Wallis statistic=0.5368, $p = 0.7646$ | No treatment effect | 10-11 mice/treatment |
| S23 | B | Diastolic blood pressure - N065, SBI-553 | One-way ANOVA on ranks | Kruskal-Wallis statistic=1.853, $p = 0.3960$ | No treatment effect | 10-11 mice/treatment |
| S20 | C | Heart ratio - N065, SBI-553 | One-way ANOVA on ranks | $F(2, 27) = 0.6803, p = 0.5149$ | No treatment effect | 9-11 mice/treatment |
| 4 | G | Flow - N065, SBI-553 | One-way ANOVA on ranks | $F(2, 28) = 3.368, p = 0.0489$ | *$p < 0.05$, as compared to Vehicle | 10-11 mice/treatment |
| S23 | D | Volume - N065, SBI-553 | One-way ANOVA on ranks | $F(2, 28) = 1.742, p = 0.1936$ | No treatment effect | 10-11 mice/treatment |
| S23 | E | VO<sub>2</sub> - N065, SBI-553 | Two-way, repeated measures ANOVA | $F_{\text{interaction}}(88, 528) = 0.9332, p = 0.6486$;<br>$F_{\text{time}}(44, 528) = 2.424$;<br>$F_{\text{treatment}}(2, 12) = 0.1799, p = 0.8376$; | No treatment effect | 5 mice/treatment |
| S23 | F | VCO<sub>2</sub> - N065, SBI-553 | Two-way, repeated measures ANOVA | $F_{\text{interaction}}(88, 528) = 0.7884, p = 0.9160$;<br>$F_{\text{time}}(44, 528) = 3.516$;<br>$F_{\text{treatment}}(2, 12) = 0.1385, p = 0.8720$; | No treatment effect | 5 mice/treatment |
| 4 | H | RER - N065, SBI-553 | Two-way, repeated measures ANOVA | $F_{\text{time}}(2.928, 11.71) = 11.64, p = 0.0008$<br>$F_{\text{treatment}}(1.052, 4.208) = 2.618, p = 0.1782$<br>$F_{\text{interaction}}(3.671, 14.69) = 1.668, p = 0.2127$ | No treatment effect | 5 mice/treatment |
| S23 | G | Heat - N065<br>SBI-553 | Two-way,<br>repeated measures<br>ANOVA | $F_{\text{interaction}}(88, 528) = 0.8069, p=0.8933$<br>$F_{\text{time}}(44, 528)=2.372$<br>$F_{\text{treatment}}(2, 12)=1.164, p=0.3450$ | No treatment effect | 5 mice/<br>treatment |
| 4 | I | RER - N065<br>SBI-553 | One-way ANOVA<br>on ranks | $F(2, 12) = 4.053, p=0.0452$ | *$p<0.05$, as compared to SBI-553 | 5 mice/<br>treatment |

<sup>1</sup>Post-hoc Tukey multiple comparison test for parametric tests and Dunn's multiple comparisons test for the Kruskal-Wallis test on ranks.

**Table S2. Curve parameters and statistical comparisons presented in Figure 4 and Figure S23 -Supporting Figure 4 and Figure S23.**

| Figure | Panel | Experiment | Statistical Test | F Statistics | Multiple Comparisons<sup>1</sup> | N |
| --- | --- | --- | --- | --- | --- | --- |
| 4 | J | Food intake - N065, SBI-553 | One-way ANOVA on ranks | Kruskal-Wallis statistic=13.85, p=0.0010 | **p<0.01, N065 as compared to vehicle;<br>##p<0.01, N065 as compared to SBI-553; | 10 mice/treatment |
| 4 | K | Food intake - N065 | Multiple tests | First 1.5h: t=3.226; df=18.00;<br>Fourth 1.5h: t=3.218; df=18.00; | **p<0.01, N065 as compared to vehicle; | 10 mice/treatment |
| S23 | H | Food intake - N065 | Unpaired tests | t=2.856, df=18; | *p<0.05, N065 as compared to vehicle; | 10 mice/treatment |
| 4 | L | Body weight change - N065, SBI-553 | Two-way, repeated measures ANOVA | F<sub>interaction</sub>(34, 391)=2.288, P<0.0001; F<sub>time</sub>(17, 391)=279.9, P<0.0001; F<sub>treatment</sub>(2, 23)=4.849, P=0.0175; | **p<0.01, SBI-553 as compared to vehicle;<br>#p<0.05, ##p<0.01, ###p<0.001, SBI-553 as compared to N065; | 8-9 mice/treatment |
| 4 | M | Body weight - N065, SBI-553 | One-way ANOVA on ranks | F(2, 22)=3.435, p=0.0503 | #p<0.05, SBI-553 as compared to N065; | 8-9 mice/treatment |
| S23 | K | Fat - N065, SBI-553 | One-way ANOVA on ranks | F(2, 22)=1.038, p=0.3710 | No treatment effect | 8-9 mice/treatment |
| S23 | I | Fat mass - N065, SBI-553 | One-way ANOVA on ranks | F(2, 24)=0.1674, p=0.8468 | No treatment effect | 9 mice/treatment |
| S23 | J | Lean mass - N065, SBI-553 | One-way ANOVA on ranks | F(2, 24)=0.1848, p=0.8324 | No treatment effect | 9 mice/treatment |

<sup>1</sup>Post-hoc Tukey multiple comparison test for parametric tests and Dunn's multiple comparisons test for the Kruskal-Wallis test on ranks.

**Table S3. Curve parameters and statistical comparisons presented in Figure 5 - Supporting Figure 5.**

| Figure | Panel | Experiment | Statistical Test | F Statistics | Multiple Comparisons<sup>1</sup> | N |
| --- | --- | --- | --- | --- | --- | --- |
| 5 | B | Effect of administered N065, SBI-553 on PD149163-induced hypothermia | Two-way, repeated measures ANOVA | $F_{\text{interaction}}(45, 504)=38.00, p<0.0001$<br>$F_{\text{time}}(1.563, 87.53)=347.8, p<0.0001$<br>$F_{\text{treatment}}(5, 56)=76.74, p<0.0001$ | *$p<0.05$, **$p<0.01$, ***$p<0.001$, ****$p<0.0001$, as compared to treatment Vehicle + Vehicle treatment at indicated N065 and SBI-553 doses and time points | 9-11 mice/treatment |
| 5 | D | Effect of administered N065, SBI-553 on low-dose LPS-induced hypothermia | Two-way, repeated measures ANOVA | $F_{\text{interaction}}(40, 2820)=13.06, p<0.0001$<br>$F_{\text{time}}(2.373, 334.6)=564.7, p<0.0001$<br>$F_{\text{treatment}}(2, 141)=12.26, p<0.0001$ | *$p<0.05$, **$p<0.01$, ***$p<0.001$, ****$p<0.0001$, as compared to treatment LPS treatment at indicated N065 and SBI-553 doses and time points | 39-53 mice/treatment |
| 5 | E | Effects of N065 and SBI-553 on survival rate reduction induced by low-dose LPS | Log-rank (Mantel-Cox) test | Chi square=13.85, $p=0.0010$ | $P=0.0041$, LPS+N065 as compared to LPS<br>$P=0.0026$, LPS+SBI-553 as compared to LPS<br>$P=0.5570$, LPS+N065 as compared to LPS+SBI-553 | 39-53 mice/treatment |
| 5 | G | Effect of administered N065, SBI-553 on high-dose LPS-induced hypothermia | Two-way, repeated measures ANOVA | $F_{\text{interaction}}(38, 1216)=6.862, p<0.0001$<br>$F_{\text{time}}(3.583, 229.3)=1108, p<0.0001$<br>$F_{\text{treatment}}(2, 64)=8.395, p=0.0006$ | *$p<0.05$, **$p<0.01$, ***$p<0.001$, as compared to LPS treatment at indicated N065 and SBI-553 doses and time points | 21-23 mice/treatment |
| 5 | H | Effects of N065 and SBI-553 on survival rate reduction induced by high-dose LPS | Log-rank (Mantel-Cox) test | Chi square=14.87, $p=0.0006$ | $p=0.0002$, LPS+N065 as compared to LPS<br>$p=0.0663$, LPS+SBI-553 as compared to LPS<br>$p=0.0284$, LPS+N065 as compared to LPS+SBI-553 | 21-23 mice/treatment |
| 5 | J | Effects of N065 on survival rate reduction induced by high-dose LPS at 30 °C | Two-way, repeated measures ANOVA | $F_{\text{time}}(24, 549) = 89.96, p<0.0001$;<br>$F_{\text{treatment}}(1, 38) = 1.575, p=0.2172$;<br>$F_{\text{interaction}}(24, 549) = 2.840, p<0.0001$ | *$p<0.05$, **$p<0.01$, as compared to LPS treatment at indicated N065 doses and time points | 1-20 mice/treatment |
| 5 | K | | Log-rank (Mantel-Cox) test | Chi square=0.7297, $p=0.3930$ | $P=0.3930$, as compared to LPS treatment at indicated N065 doses and time points | 1-20 mice/treatment |

<sup>1</sup>Post-hoc Tukey multiple comparison test for parametric tests and Dunn's multiple comparisons test for the Kruskal-Wallis test on ranks.

**Table S4. Curve parameters and statistical comparisons presented in Figure S24–Supporting Figure S24.**

| Figure | Panel | Experiment | Statistical Test | F Statistics | Multiple Comparisons<sup>1</sup> | N |
| --- | --- | --- | --- | --- | --- | --- |
| S24 | A | Latency to platform - N065, SBI-553 | Two-way, repeated measures ANOVA | $F_{\text{interaction}}(8, 116)=0.6953, P=0.6951$; $F_{\text{time}}(3.463, 100.4)=12.14, P<0.0001$; $F_{\text{treatment}}(2, 29)=0.9307, P=0.4057$ | *$p<0.05$, SBI-553 as compared to vehicle at doses and indicated time points | 10-11 mice/treatment |
| S24 | B | Spontaneous Alternation (%) - N065, SBI-553 | One-way ANOVA on ranks | Kruskal-Wallis statistic=7.016, $P=0.03$ | *$p<0.05$, SBI-553 as compared to vehicle | 23 mice/treatment |
| S24 | C | Total no. of arm entries - N065, SBI-553 | One-way ANOVA on ranks | $F(2, 66) = 16.79, P<0.0001$ | ***$p<0.001$, ****$p<0.0001$, as compared to vehicle | 23 mice/treatment |
| S24 | D | Discrimination index - N065, SBI-553 | One-way ANOVA on ranks | $F(2, 27) = 0.4472, P=0.6441$ | No treatment effect | 9-11 mice/treatment |
| S24 | E | Time in open arms - N065, SBI-553 | One-way ANOVA on ranks | $F(2, 29) = 0.1294, P=0.8791$ | No treatment effect | 10-11 mice/treatment |
| S24 | F | Post test consumption - N065, SBI-553 | One-way ANOVA on ranks | Kruskal-Wallis statistic=12.12, $P=0.0023$ | **$p<0.01$, N065 as compared to vehicle; #$p<0.05$, N065 as compared to SBI-553; | 9-11 mice/treatment |
| S24 | G | Latency to first approach - N065, SBI-553 | One-way ANOVA on ranks | $F(2, 24) = 0.6117, P=0.5507$ | No treatment effect | 8-10 mice/treatment |
| S24 | H | Immobility time - N065, SBI-553 | One-way ANOVA on ranks | $F(2, 27) = 2.011, P=0.1534$ | No treatment effect | 10 mice/treatment |

<sup>1</sup>Post-hoc Tukey multiple comparison test for parametric tests and Dunn's multiple comparisons test for the Kruskal-Wallis test on ranks.

**Table S5. Cryo-EM data collection, refinement and validation statistics**

|  | N065 | N191 | NA-SBI553-2 |
| --- | --- | --- | --- |
| **Data collection and processing** |  |  |  |
| Magnification | 165K | 165K | 105K |
| Voltage (kV) | 300 | 300 | 300 |
| Electron exposure (e-/Å<sup>2</sup>) | 50 | 50 | 50 |
| Defocus range (μm) | 0.8-1.8 | 0.8-1.8 | 0.8-1.8 |
| Pixel size (Å) | 0.73 | 0.73 | 0.824 |
| Symmetry imposed | C1 | C1 | C1 |
| Final particle images (no.) | 180083 | 248821 | 968592 |
| Map resolution (Å) | 3.35 | 3.11 | 2.72 |
| FSC threshold | 0.143 | 0.143 | 0.143 |
| Map resolution range (Å) | 2.5-5 | 2.5-5 | 2-4 |
| **Refinement** |  |  |  |
| Model resolution (Å) | 3.5 | 3.5 | 3.1 |
| FSC threshold | 0.5 | 0.5 | 0.5 |
| Model composition |  |  |  |
| Non-hydrogen atoms | 8604 | 8568 | 8572 |
| Protein residues | 1109 | 1109 | 1110 |
| Ligands | 2 | 2 | 2 |
| *B* factors (Å<sup>2</sup>) |  |  |  |
| Protein | 57.42 | 76.34 | 73.65 |
| Ligand | 94.18 | 87.21 | 61.61 |
| R.m.s. deviations |  |  |  |
| Bond lengths (Å) | 0.002 | 0.002 | 0.003 |
| Bond angles (°) | 0.547 | 0.540 | 0.603 |
| Validation |  |  |  |
| MolProbity score | 1.74 | 1.64 | 1.76 |
| Clashscore | 6.64 | 7.49 | 7.26 |
| Poor rotamers (%) | 0.54 | 0.75 | 0.84 |
| Ramachandran plot |  |  |  |
| Favored (%) | 95.78 | 96.51 | 95.50 |
| Allowed (%) | 4.22 | 3.3 | 4.50 |
| Disallowed (%) | 0 | 0.18 | 0 |

**Table S6. Summary of pharmacokinetic parameters of N065 and SBI-553 in male ICR mice following intravenous (i.v.) and intraperitoneal (i.p.) administration.**

| Compound & Route | Matrix | T<sub>max</sub> (h) | C<sub>max</sub>(ng/mL) | AUC<sub>last</sub>(hr*ng/mL) | T<sub>1/2</sub> (h) | V<sub>ss</sub> (L/kg) | CL (mL/min/kg) | B/P Ratio |
| --- | --- | --- | --- | --- | --- | --- | --- | --- |
| N065 (IV 1 mg/kg) | Plasma | - | 1545 | 744 | 2.76 | 1.96 | 22.4 | - |
| N065 (IP 12 mg/kg) | Plasma | 1.00 | 1250 | 7666 | 3.66 | - | - | 0.36 (1h) |
|  | Brain | 1.00 | 403 | - | - | - | - |  |
| SBI-553 (IV 1 mg/kg) | Plasma | - | 890 | 339 | 0.96 | 1.84 | 49.6 | - |
| SBI-553 (IP 12 mg/kg) | Plasma | 0.50 | 1212 | 4428 | 3.38 | - | - | 0.90 (1h) |
|  | Brain | 0.50 | 1001 | - | - | - | - |  |

Plasma and brain exposures were evaluated following i.v. (1 mg/kg) and i.p. (12 mg/kg) administration. For i.v. groups, C<sub>max</sub> is presented as the initial plasma concentration (C<sub>0</sub>), and T<sub>max</sub> is not applicable. Whole brains were collected only from the i.p. groups at 0.5 and 1 h post-dose; therefore, brain parameters and the brain-to-plasma (B/P) ratio are not available for the i.v. groups. The B/P ratio was calculated as the concentration ratio of brain to plasma at 1 h post-dose. T<sub>max</sub>, time to reach C<sub>max</sub>; C<sub>max</sub>, maximum concentration; AUC<sub>0-t</sub>, area under the concentration-time curve from time zero to the last measurable time point; T<sub>1/2</sub>, terminal elimination half-life; B/P, brain-to-plasma; not determined.

**Fig. S1 | Validation of TAMRA-PEG6-NTS<sup>8-13</sup>.**

**(A)** Chemical structure of the orthosteric fluorescent tracer.

**(B)** Live-cell imaging validating tracer specificity. Bright-field and fluorescence images acquired 1 min after tracer addition and 5 min after washout in cells lacking (–) or expressing (+) NTSR1. Receptor-dependent fluorescence is observed in NTSR1-positive cells with persistent punctate signal after washout, consistent with receptor-associated tracer.

**Fig. S2 | Sequence and structural comparison of SBI-553-bound NTSR1–β-arrestin complexes.**

(A) Sequence alignment of human β-arrestin1 (βarr1, UniProt P49407) and β-arrestin2 (βarr2, UniProt P32121), with 75.00% identity and 86.68% similarity. Identical residues are highlighted in red.

(B) Structural superposition of three reported NTSR1–βarr1–SBI-553 complexes (PDB: 8ZYU, 8ZYY, and 8ZYT) and the NTSR1–βarr2–SBI-553 complex determined in this study, aligned by NTSR1. The inset shows the superimposed SBI-553 molecules.

(C–F) SBI-553 binding sites in complexes 8ZYU (C), 8ZYY (D), 8ZYT (E), and the NTSR1–βarr2 complex (F). Protein residues with at least one heavy atom within 4 Å

of any SBI-553 heavy atom are shown as sticks.

**(G)** Comparison of SBI-553 binding-pocket residues across the four complexes, defined using a 5 Å heavy-atom distance cutoff. Each dot denotes a pocket residue in the corresponding complex, grouped by NTSR1 and β-arrestin.

**Fig. S3 | Cryo-EM processing of the NTSR1-βarr2-SBI-553 complex.** Representative micrographs, 2D classes, and refinement workflow; final map resolution 2.88 Å.

**Fig. S4 | Data processing of the NTSR1-βarr2-N065 complex.** Representative 2D classes and refinement pipeline; final map resolution 2.54 Å.

**Fig. S5 | Data processing of the NTSR1-βarr2-N191 complex.** Reconstruction

reached 2.63 Å with clear density for TM6–TM7 and the βarr2 finger loop.

**SBI-553-bound NTSR1/ $\beta$ arr2**

**N065-bound NTSR1/ $\beta$ arr2**

**N191-bound NTSR1/ $\beta$ arr2**

**Fig. S6 | Representative cryo-EM densities for the NTSR1–β-arrestin2 complexes bound to SBI-553, N065 or N191.**

Shown are density maps and fitted atomic models for individual NTSR1 transmembrane helices (TM1–TM6 and TM7–H8), the β-arrestin2 finger loop, the orthosteric peptide NTS<sup>8–13</sup> and the corresponding intracellular allosteric modulators, illustrating the local map quality supporting model assignment. Two views related by a 70° rotation are shown for NTS<sup>8–13</sup> and each intracellular modulator to provide complementary visualization of the surrounding density. Rows correspond to the SBI-553-, N065- and N191-bound complexes, respectively. Density is displayed as a gray mesh around the fitted atomic models.

**Fig. S7 | Comparison of intracellular ligand-binding pockets in NTSR1–Gi/o and NTSR1–β-arrestin2 complexes.** Comparison of binding-pocket residues surrounding SBI-553 and its derivatives N065, N191, N209, and N253 in Gi/o-coupled (upper five rows) and β-arrestin2 (βarr2)-coupled (lower five rows) NTSR1 complexes. Pocket residues were defined by a minimum heavy-atom distance of 5 Å between each protein residue and the corresponding ligand. X indicates a protein residue within this cutoff. The three cryo-EM structures of NTSR1–βarr2 complexes bound to SBI-553, N065, and N191 determined in this study share a conserved receptor pocket involving L105<sup>2.43</sup>, V159<sup>3.43</sup>, R166<sup>3.50</sup>, V304<sup>6.36</sup>, V308<sup>6.40</sup>, and Y364<sup>7.53</sup>, but differ in peripheral pocket residues and βarr2 finger-loop contributions. L88<sup>1.57</sup> and L252<sup>5.54</sup> are present in the N065 and N191 pockets, whereas N370<sup>8.49</sup> is observed in the SBI-553 pocket. The βarr2 finger-loop residue N.s5s6.05L is included in the SBI-553 and N065 pockets but not in the N191 pocket. Residues are grouped by NTSR1, Gi/o, and βarr2. Complexes labeled “dock” correspond to molecular docking models.

**Fig. S8 | Surface expression level and NTS<sup>8-13</sup> agonist activity of NTSR1 mutants across pathways.**

**(A)** Surface expression level of mutants (normalized to WT).

**(B–E)** NTS<sup>8-13</sup> agonist activity of NTSR1 mutants across four pathways.

### SBI-553

**Fig. S9 | SBI-553 agonist activity of NTSR1 mutants across pathways.**

**Fig. S10 | N065 agonist activity of NTSR1 mutants across pathways.**

**Fig. S11 | N191 agonist activity of NTSR1 mutants across pathways.**

**Fig. S12 | Structural mapping of transducer-sensitive residues identified by alanine scanning onto NTSR1 structures in different ligand-bound states.**

Residues whose alanine substitutions produced loss-of-function (LOF) effects across four transducer pathways, gain-of-function (GOF) effects in G-protein signaling, or bidirectional, transducer-dependent regulation are indicated by orange triangles, purple triangles and blue diamonds, respectively. The dashed line denotes the approximate boundary of the intracellular receptor–transducer interface. The orthosteric agonist NTS<sup>8-13</sup> is shown in blue, and the intracellular modulators SBI-553 and N191 are shown in cyan and green, respectively. W316<sup>6.48</sup>, located outside the immediate intracellular ligand-contact region, is highlighted as a conserved activation-motif residue exhibiting transducer-dependent functional effects. The spatial distribution of these residues indicates that functionally sensitive positions extend beyond the immediate BAM-binding interface and occupy multiple regions of the intracellular receptor core.

**Fig. S13 | Systematic classification of mutation-dependent changes in intracellular allosteric modulation across $G_q$, $G_i$, $G_{13}$ and β-arrestin2 signaling.**

Alanine mutants were classified relative to the corresponding wild-type response according to whether allosteric modulation was weakened, enhanced, shifted to a different regulatory mode, or reversed in direction. Donut plots summarize the number of mutants in each category for SBI-553/N191 and N065 across the indicated signaling pathways. Representative schematic concentration–response curves illustrate the corresponding changes in modulatory behavior, including attenuation or enhancement of NAM or ago-PAM activity, transitions between modulatory modes, and reversal of modulation direction. Structural maps show the spatial distribution of residues associated with each response category within the NTSR1 intracellular region. Selected residues exhibiting prominent mode shifts or reversals, including W316<sup>6,48</sup>, D112<sup>2,50</sup>, N256<sup>5,58</sup> and G301<sup>6,33</sup>, are indicated. These data demonstrate that the functional consequences of alanine substitution are both ligand- and pathway-dependent, revealing a non-uniform contribution of intracellular receptor residues to BAM-mediated signaling modulation.

**Fig. S14–15 | Allosteric effect curves of SBI-553, N065 and N191 from the NTSR1 mutants in G<sub>q</sub> signaling pathway.**

**Fig. S16–17 | Allosteric effect curves of SBI-553, N065 and N191 from the NTSR1 mutants in G<sub>i</sub> signaling pathway.**

**Fig. S18–19 | Allosteric effect curves of SBI-553, N065 and N191 from the NTSR1 mutants in G<sub>13</sub> signaling pathway.**

654

**Fig. S20–21 | Allosteric effect curves of SBI-553, N065 and N191 from the NTSR1 mutants in β-arrestin2 signaling pathway.**

**Fig. S22 | Pharmacokinetic parameters of N065 and SBI-553 in ICR mice.**

**(A)** Plasma concentration-time curves of N065 and SBI-553 following i.v. administration at 1 mg/kg (n = 3 per time point).

**(B)** Plasma concentration-time curves of N065 and SBI-553 following i.p. administration at 12 mg/kg (n = 3 per time point).

**(C)** Brain concentration-time curves of N065 and SBI-553 following i.p. administration at 12 mg/kg. Brain tissues were collected only at 0.5 and 1 h post-dose (n = 3 per time point).

**(D)** Brain-to-plasma (B/P) ratio of N065 and SBI-553 following i.p. administration at 12 mg/kg (n = 3 per time point).

Data are presented as mean ± SEM. Blood samples were collected via the retro-orbital venous plexus at 0.083, 0.25, 0.5, 1, 2, 4, 8, and 24 h post-dose.

**Fig. S23 | Effects of N065 and SBI-553 on diverse phenotypes in mice.**

**(A)** Wire hang test. Time the mouse hung on and explored the underside of the wire grid before releasing into cage below. Animals received vehicle, N065 or SBI-553 (i.p.) 30 min prior to testing.

**(B-D)** Diastolic blood pressure (mmHg) (B), heart rate (bpm) (C) and blood volume (μL) (D) was continuously recorded by tail-cuff system CODA in mice 30 min after treatment with vehicle, N065 or SBI-553 (i.p.). Baseline was set to 0.

**(E-G)** Metabolic cage analysis of mice treated with vehicle, N065 or SBI-553. Oxygen consumption (E), CO<sub>2</sub> output (F) and energy expenditure (G).

**(H)** Regular food intake in C57BL/6 mice were evaluated under fed states. Mice were administered N065 (12 mg/kg), SBI-553 (12 mg/kg), or vehicle 30 min before the feeding test.

**(I-K)** Body composition analysis generated by echoMRI scan of DIO mice treated with vehicle, N065 or SBI-553 from day 34 of HFD diet. Percentage of fat mass (I), percentage of lean mass (J) and Fat mass (g) (K).

For details on statistical comparisons, see Table S1 and Table S2.

**Fig. S24 | Effects of N065 and SBI-553 on behaviors in male mice.**

**(A)** Morris water maze test. Animals received vehicle, N065 or SBI-553 (i.p.) 1 h prior to testing. Latency to platform was recorded.

**(B-C)** Y-maze task. Animals received vehicle, N065 or SBI-553 (i.p.) 1 h prior to testing. (B) Spontaneous alternation. (C) Total arm entries.

**(D)** Novel object recognition test. Animals received vehicle, N065 or SBI-553 (i.p.) 1 h prior to testing. Discrimination index (exploration novel object- exploration same object)/total exploration time) is tested.

**(E)** Elevated plus maze test. Animals received vehicle, N065 or SBI-553 (i.p.) 1 h prior to testing. Time spent in the open arms was recorded.

**(F-G)** Novelty-suppressed feeding test. Animals received vehicle, N065 or SBI-553 (i.p.) 1 h prior to testing. (F) Amount of consumed food within 10 min. (G) latency period of first feeding.

**(H)** Forced swim test. Animals received vehicle, N065 or SBI-553 (i.p.) 1 h prior to

716 testing. Immobility time was recorded.
717 For details on statistical comparisons, see Table S4.
718

**Fig. S25 | Global QC, differential statistics, KSEA, and KEGG analyses of the phosphoproteomic dataset.**

**(A)** Within-group coefficient of variation (CV) distributions across treatment conditions, showing the reproducibility of phosphoproteomic quantification.

**(B)** Numbers of differential phosphosites across pairwise comparisons.

**(C)** Circular volcano plot summarizing global differential phosphosite distributions across multiple comparisons. Significant up- and down-regulated sites were defined as $|\log_2FC| > 1.2$ and $p < 0.05$; remaining sites were classified as non-significant (NS).

**(D)** KSEA activity heatmap showing kinase enrichment z-scores across multiple comparisons. Only kinase sets with at least five matched substrate sites were retained ($m \geq 5$), and significantly enriched kinase sets ($p < 0.05$) are shown. Row clustering was used to compare kinase-program similarities and differences across conditions.

**(E)** Classification of allosteric modulation of NTS-enriched pathway sets. Significantly enriched KEGG pathways relative to vehicle were classified as retained, gained, or lost. Retained pathways overlapped with the NTS program, gained pathways were newly enriched under co-treatment, and lost pathways were enriched in the NTS condition but absent from the corresponding co-treatment.

**(F–H)** KEGG enrichment bubble plots for the three co-treatment groups relative to NTS: (F) NTS+SBI-553 vs NTS, (G) NTS+N065 vs NTS, (H) NTS+N191 vs NTS. The x-axis indicates rich factor, bubble size indicates the number of hits, and color indicates *padj*.
